# Aging compromises Zebrafish caudal fin regeneration by disrupting Regenerative gene networks and Cellular metabolism

**DOI:** 10.64898/2026.03.24.713633

**Authors:** PV Anusha, Qasid Ahmed, PV Athira, Anushka Arvind, Izzath Fathima, PS Basil, Mohammed Ghalib Enayathullah, Mohammed Mundhir, Ifra Iyoob, Sarena Banu, Jessica Bharathi, Sirin Bano, Saloni Garg, Jameela Bano, Samreen Fatma, Mohammed Lukman Rafi, Cader O Salma, Jahangir Alom, Mohammed Arsalan, Hari Krishna, Shiv Pratap Singh Yadav, Mohammed M Idris

**Author notes:** Corresponding Author Dr. Mohammed M Idris Chief Scientist, CSIR-CCMB, Hyderabad, India.

## Abstract

Zebrafish are widely recognized as a powerful vertebrate model for studying epimorphic regeneration due to their remarkable ability to restore complex tissues. However, regenerative efficiency declines with age, potentially due to alterations in gene regulatory networks and cellular metabolism. In the present study, we investigated the molecular and bioenergetic basis of age-associated regenerative decline by comparing young adult (<1 year) and old adult (>3 years) zebrafish during caudal fin regeneration. To further examine the contribution of mitochondrial function, mitochondrial dysfunction was experimentally induced using rotenone (20 nM), a mitochondrial Complex I inhibitor. Regenerative progression was assessed morphologically at 12hpa, 1dpa, 2dpa, 3dpa, and 7dpa, revealing a pronounced delay in fin regrowth in aged and rotenone-treated fish compared with young controls. Behavioral analysis indicated subtle but non-significant changes across experimental groups. Gene expression analysis using quantitative real-time PCR revealed age- and mitochondria-associated dysregulation of key regenerative gene families involved in developmental patterning, extracellular matrix organization, cellular signaling, and mitochondrial metabolism. Proteomic profiling further identified differential expression of proteins associated with mitochondrial bioenergetics, extracellular matrix remodeling, and signaling pathways required for blastema formation and tissue outgrowth. Ultrastructural examination by transmission electron microscopy revealed pronounced mitochondrial abnormalities, including enlarged mitochondria with fragmented or disrupted cristae, in aged and rotenone-treated regenerating tissues. Collectively, our integrative analysis establishes a mechanistic link between aging, mitochondrial dysfunction, and compromised regenerative capacity in zebrafish. The findings provide broader insights into metabolic constraints underlying age-related decline in regenerative potential in vertebrates.

## Introduction

Regeneration is a complex biological process that enables organisms to restore lost or damaged tissues through highly coordinated cellular and molecular mechanisms. While higher vertebrates exhibit limited regenerative capacity, certain lower vertebrates, such as zebrafish (*Danio rerio*), possess an exceptional ability to regenerate various organs, including the heart, spinal cord, and appendages [1, 2]. Zebrafish caudal fin regeneration, in particular, serves as a robust model of epimorphic regeneration due to its reproducible sequence of healing events such as wound closure, blastema formation, proliferative outgrowth, and tissue differentiation into bone, epidermis, and vasculature [3, 4, 5].

Several signaling pathways and gene families have been implicated in zebrafish fin regeneration, including Wnt/β-catenin, fibroblast growth factor (FGF), bone morphogenetic proteins (BMPs), and homeobox (HOX) genes, among others [6, 7]. Our previous study has revealed a distinct expression pattern across several gene families, including HOX, Keratin, TMEM, and SLC genes in young zebrafish during fin regeneration. These gene sets were found to play central roles in tissue patterning, structural remodeling, membrane transport, and cellular signaling during regeneration. Importantly, their temporal expression correlated strongly with the stages of regenerative outgrowth, suggesting a dynamic and tightly regulated genetic program [5]. However, despite extensive studies in young adult zebrafish, the impact of aging on these regenerative processes and their underlying molecular dynamics remains poorly characterized. In the present study, we build on this foundation to examine whether the same gene expression programs are preserved, attenuated, or misregulated in aged zebrafish. Our objective is to delineate the molecular footprint of aging in the regenerative landscape.

Aging is associated with a decline in regenerative potential, often linked to changes in gene expression, impaired cellular signaling, and reduced proliferative capacity [8]. Understanding how aging affects key gene families involved in fin regeneration can provide valuable insights into the mechanisms underlying age-related decline in tissue repair. In this study, we systematically analyzed the expression of multiple gene families, including SRY-related HMG-box (SOX) transcription factors, Homeobox (HOX) genes, Muscle segment homeobox (MSX) genes along with their interacting partners, Keratin, Protein Kinases, Laminin, Transmembrane (TMEM) genes, Collagen, Solute carrier (SLC), Neuronal, Notch, Mitochondrial genes and aging-associated genes, in regenerating caudal fin tissue of young and old zebrafish.

The gene families assessed in this study are central to diverse but interconnected to biological functions critical for regeneration. One of the hallmark transcriptional regulators during fin regeneration is the MSX gene family. MSX homeobox genes, particularly *msxb*, are transiently expressed during the early post-amputation phase, especially during blastema formation and later becomes restricted to the distal-most blastema (DMB), a zone characterized by slow-cycling or non-proliferating cells, where they modulate apoptosis, dedifferentiation, and proliferation. Given that MSX homeodomain proteins are known transcriptional repressors associated with preserving cells in an undifferentiated state, *msxb* likely contributes to creating a regulatory environment essential for proper regenerative patterning and directional growth. It has been proposed that these non-cycling *msxb*-expressing cells at the DMB may provide spatial cues necessary for organizing the regenerative outgrowth and ensuring proper tissue architecture during fin regrowth [2, 9]. HOX genes, that encode homeodomain transcription factors are shown to be expressed in regenerating tissues including zebrafish fins. The expression of the *Hox* genes during a regeneration event represent re-patterning of the tissues, analogous to their role in embryonic development [10]. In our previous study, genes such as *hoxa10b*, *hoxa13b*, *hoxc13b*, and *hoxd10a* showed prominent expression from wound closure to regenerative outgrowth [6]. SOX genes encode a family of transcription factors and belong to a superfamily of genes characterized by a homologous sequence called the HMG-box, that is highly conserved throughout eukaryotic species. The SOX genes are involved in the regulation of diverse growth-related and developmental processes, such as chondrogenesis, neurogenesis, and sex determination and differentiation. The genes such as *sox9a* and *sox10*, are critical for stem cell maintenance and lineage specification, and are reactivated during regeneration to maintain progenitor identity and promote differentiation [11]. Keratin genes, which encode intermediate filament proteins in epithelial cells, are essential for re-epithelialization and wound closure. Collagens, primarily *col1a1a*, *col5a1*, and *col12a1a*, form the structural scaffold of the ECM. Proper collagen synthesis, cross-linking, and degradation are essential for tissue remodeling. TMEM genes, although diverse in function, are increasingly recognized for their roles in membrane signaling, ion transport, and homeostasis. Notably, the *tmem9* gene was found to be downregulated during zebrafish caudal fin tissue regeneration [6].

Notch signaling, another critical juxtacrine pathway, has been implicated in regulating cell fate during development and regeneration. Its activation has been observed in blastema cells, influencing proliferation and maintenance of undifferentiated states [12]. Meanwhile, Kinases are involved in signalling pathways which regulate diverse cellular functions such as cell cycle progression, apoptosis, metabolism, differentiation, cell morphology, cell migration and secretion of cellular proteins. Laminins, components of the extracellular matrix (ECM), are central to establishing a regenerative niche. They support cell adhesion, migration, and survival during re-epithelialization and blastema proliferation. Laminins also cross-talk with integrin and Fgf signaling, thereby integrating environmental and intrinsic cues. The laminin beta 1a (lamb1a) gene, is crucial for establishing polarity in basal epithelial cells, inducing and maintaining basal epithelial markers, and localizing signaling receptors during fin regeneration [13]. Solute carrier (SLC) genes code for membrane-bound transporters which carry amino acids, neurotransmitters, lipids, and other substrates across all cellular and organelle membranes with the exception of the nuclear membrane. There are several subfamilies of SLC genes, of which control the transport of substances like amino acids, calcium, potassium, sodium, sulfur compounds, folic acid, nucleotide-sugars, bile acid, L-gluatmine, and zinc ions. These processes are essential in providing zebrafish with the necessary nutrients and maintaining homeostasis in order to initiate and follow through with all steps of regeneration. In addition, various genes have been found to play a role in melanin pigmentation and melanosome biogenesis, one of the final steps of caudal fin restoration. The knockdown of some of these genes, including slc25a1a and slc25a1b have been found to result in abnormal fin morphology, emphasizing the importance of SLC genes in the regenerative process. Neuronal genes are also included as the functional recovery of fin depends not only on morphological restoration but also on sensory integration and neuronal growth.

Among the various regulatory networks involved in caudal fin regeneration, mitochondrial dynamics, and bioenergetics have emerged as critical factors as mitochondria generate oxidative metabolic signals essential for cellular homeostasis and tissue [14]. Mitochondria, being the powerhouse of the cell, are essential for cellular energy production, providing a continuous energy supply necessary for the growth, development, and regeneration of zebrafish’s regenerating fin tissue. Key mitochondrial genes involved in oxidative phosphorylation (OXPHOS) such as mt-nd1, nd4, nd4L, nd5, nd6, mt-co1, co2, and mt-cyb encode components of the electron transport chain (ETC) and are shown to fluctuate during the regeneration process, reflecting the high energy demand and metabolic reprogramming required for cell proliferation, differentiation, and tissue remodeling. The mt-nd1, nd4, nd5, and nd6 genes encode Complex I subunits, which initiate electron transfer from NADH to ubiquinone, establishing the proton gradient for ATP synthesis. Their upregulation during regeneration highlights the tight regulation of mitochondrial bioenergetics to meet energy demands [15, 16]. The mt-cyb gene encodes a Complex III subunit, facilitating electron transfer between ubiquinol and cytochrome c, crucial for respiratory efficiency and ATP generation [17, 18]. The mt-co1 and mt-co2 genes encode Complex IV subunits, which transfer electrons to oxygen, forming water and contributing to ATP production. Cytochrome c oxidase is crucial for maintaining the mitochondrial membrane potential and ensuring efficient ATP production [19, 20].

Finally, our study included aging-associated genes such as *igf1*, *igfbp2a*, *tert*, *hsp70*, and *atm,* that serve as key regulators of cellular growth, genomic maintenance, and stress resilience. Their expression modulates pathways involved in insulin-like growth factor signaling, telomere preservation, proteostasis, and DNA damage response, all of which are critical for sustaining regenerative capacity. The altered regulation of these genes in aged zebrafish may underlie the decline in cellular proliferation, increased susceptibility to oxidative stress, and reduced structural restoration observed during fin regeneration.

Given the intimate relationship between mitochondrial function and cellular plasticity, we hypothesized that aging-associated decline in regenerative capacity may be mechanistically linked to altered mitochondrial bioenergetics. To test this, mitochondrial dysfunction was experimentally induced using rotenone, a well-characterized Complex I inhibitor known to elevate oxidative stress and disrupt mitochondrial membrane potential [21]. Studies show that rotenone exposure in zebrafish embryos causes high mortality and developmental abnormalities, including cardiac edema, tail deformities, loss of equilibrium, delayed somite formation, impaired yolk sac absorption, and lack of pigmentation [22, 23]. As a mitochondrial Complex I inhibitor, rotenone increases reactive oxygen species (ROS) production via microglial NADPH oxidase, depletes antioxidant enzymes like glutathione, elevates lipid peroxidation, and triggers mitochondrial membrane depolarization [24].

By integrating morphological assessment, behavioral analysis, quantitative gene expression profiling, proteomic characterization, and ultrastructural examination through transmission electron microscopy, this study delineates the molecular and bioenergetic landscape underlying age-dependent decline in zebrafish caudal fin regeneration. Collectively, our findings provide a comprehensive framework linking developmental gene network dysregulation and mitochondrial dysfunction to impaired regenerative competence with aging, offering broader insights into the metabolic control of tissue repair in vertebrates.

## Materials and Methods

### Animal Maintenance and Sample Collection

Wild-type zebrafish (*Danio rerio*) were maintained under standard laboratory conditions at 28°C with a 14:10-hour light-dark cycle and fed a controlled diet consisting of dry feed and live brine shrimp. Adult zebrafish were categorized into two groups based on age: young (<1 year; 9–12 months) and old (>3 years; 36–48 months). Caudal fin amputations were performed on anesthetized fish (0.02% MS-222) using a sterile scalpel blade following established protocols. Approximately 50% of the caudal fin was removed. Regenerating fin tissues were collected at six time points: 0 hours (0hpa), 12 hours (12hpa), 1 day (1dpa), 2 days (2dpa), 3 days (3dpa), and 7 days (7dpa) post-amputation. Each experimental group consisted of five animals per time point, performed in two independent batches. All experiments were conducted in accordance with institutional animal ethical guidelines and approved protocols (IAEC/CCMB/Protocol #50/2013).

### Rotenone Treatment

Prior to experimental analysis, various rotenone concentrations were tested, and 20 nM rotenone was determined to be effective without causing mortality, whereas higher concentrations resulted in lethality. To induce mitochondrial dysfunction, zebrafish were exposed to 20 nM rotenone dissolved in system water for 48 hours prior to caudal fin amputation. Rotenone treatment was continued throughout the entire regeneration period (up to 7 days post-amputation, dpa). Control fish were maintained under identical conditions without rotenone treatment. Following exposure, fins were amputated as described above, and tissues were collected at corresponding post-amputation time points for molecular, proteomic, and ultrastructural analyses.

### Phenotypic Analysis of Regeneration

Regenerated caudal fins were captured at each time point using a stereomicroscope. Fin regeneration was assessed by measuring the regenerated length of fin lobe and cleft, using ImageJ software. Quantitative measurements were statistically analyzed to compare regeneration efficiency across young, aged, and rotenone-treated groups.

### Behavioral Analysis

To assess the behavioral responses associated with caudal fin amputation and subsequent regeneration, we employed the Novel Tank Test (NTT), a validated method for analyzing locomotor activity and exploratory behavior in zebrafish. Behavioral analysis was conducted on both young and old zebrafish at six post-amputation time points: 0h, 12h, 1d, 2d, 3d, and 7d, alongside an unamputated group serving as a control. At each time point, n = 5 individual fish were analyzed. For each fish, a 3-minute video recording was obtained in a standardized novel tank environment under uniform lighting and noise conditions. Behavioral parameters such as total distance moved, average velocity, freezing duration, and zone preference (top vs. bottom dwell time) were quantified using EthoVision XT software (Noldus Information Technology). Both individual fish behavior and group-averaged trends were analyzed for each time point and age group. This behavioral assay aimed to capture potential age- and regeneration-associated differences in activity and anxiety-like behavior in zebrafish, providing an additional functional perspective alongside molecular and proteomic analyses.

### NGS Data Analysis

Gene families analyzed in the present study were selected based on our previously published transcriptomic analysis of zebrafish caudal fin regeneration [5]. Next-generation sequencing was performed using the Illumina HiSeq 2000 platform on total RNA isolated from regenerating caudal fin tissues collected at multiple post-amputation time points. De novo transcriptome assembly, functional annotation using BLASTx against the *Danio rerio* database, and differential expression analysis using FPKM normalization were carried out. Zero hours post-amputation (0h) was used as the baseline control. Transcripts showing >1 log-fold change were considered significantly differentially expressed. Based on these data, regeneration-associated gene families were identified, and selected genes were further evaluated in the present study to determine age- and mitochondria-associated alterations in gene expression.

### Quantitative Real-Time PCR (qRT-PCR) Analysis

Total RNA was extracted from regenerating fin tissues of young, old and rotenone treated fishes (n=5) at six time points (0hpa, 12hpa, 1dpa, 2dpa, 3dpa, and 7dpa) using TRIzol reagent (Invitrogen, USA). cDNA synthesis was performed using 2μg of total RNA per sample with the iScript cDNA Synthesis Kit (Bio-Rad, USA). Gene expression analysis was conducted for regeneration-associated gene families including SOX, HOX, MSX, MSX interacting partners, Keratin, Kinase, Laminin, TMEM, Collagen, Solute Carrier (SLC), Neuronal genes, Aging-associated genes, and mitochondrial genes. Quantitative real-time PCR was performed using SYBR Green Master Mix (Bio-Rad, USA) on a QuantStudio 6 Flex Real-Time PCR System (Applied Biosystems, USA). Primers were designed using Primer3 software based on transcriptomic sequences and NCBI *Danio rerio* database annotations. The ODC gene was used as the internal reference gene. All reactions were performed in biological and technical duplicates. Relative gene expression was calculated using the 2^(-ΔΔCt) method, with 0h post-amputation samples serving as the baseline control.

### Protein Extraction and Quantification

Total protein from regenerating caudal fin tissues of young and old fishes (n=5) was extracted using a urea-based protein solubilization buffer containing protease inhibitors [25]. Lysates were centrifuged to remove debris, and supernatants were collected. Protein concentration was determined using the Amido Black assay with bovine serum albumin (BSA) as the standard.

### Proteomic Analysis

Total protein from regenerating caudal fin tissues at different time points (0hpa, 12hpa, 1dpa, 2dpa, 3dpa, and 7dpa) was subjected to quantitative proteomic analysis using both label-free quantification (LFQ) and isobaric tags for relative and absolute quantitation (iTRAQ). For iTRAQ-based proteomics, protein samples were resolved on 10% SDS-PAGE, digested using trypsin, and labeled with iTRAQ 4-plex reagents (AB Sciex, USA). Labeled peptides were purified using C-18 spin columns (Thermo Scientific, USA). For label-free quantification, trypsin-digested peptides were purified using C-18 spin columns prior to analysis. Purified peptides were reconstituted in 5% acetonitrile (ACN) and 0.2% formic acid and analyzed using an LC-MS/MS Orbitrap Velos Nano analyzer (Thermo Scientific). Raw data were processed using Proteome Discoverer 2.3 (Thermo Fisher Scientific, USA) with the Sequest HT search algorithm and a 1% false discovery rate (FDR) cutoff. *Danio rerio* protein sequences were used as the reference database. Proteins exhibiting at least one log-fold differential expression between age groups and time points were considered significant for further analysis.

### Transmission Electron Microscopy (TEM)

Regenerating fin tissues from young, old, and rotenone-treated fish were rinsed with 1X PBS and fixed in 2.5% glutaraldehyde at 4°C overnight, followed by washing with PBS and post-fixation in 2% osmium tetroxide (OsO_4_) for 60-120 minutes. Tissues were subsequently stained with Uranyless for 45 minutes at room temperature. Samples were dehydrated through a graded ethanol series (50%, 70%, 95%, and 100%) and treated with propylene oxide (2 × 5 minutes), followed by infiltration with epoxy resin–propylene oxide mixtures (1:1, 2:1, 3:1, and pure resin). Samples were embedded in epoxy resin mixture (ERL 4221, DER 736, NSA, and DMAE) and polymerized at 70°C for 8 hours or 60 °C for 14 hours. Embedded samples were trimmed to expose the blastema region. Ultrathin sections were cut using an ultramicrotome, collected on copper grids, and counterstained with Uranyless and lead citrate. Sections were examined using a Talos L 120C Transmission Electron Microscope equipped with a Ceta camera. Images were analyzed using Velox software. Mitochondrial morphology was evaluated across young, aged, and rotenone-treated zebrafish to assess ultrastructural changes during regeneration and aging.

### Data Analysis and Pathway Enrichment

Differentially expressed genes and proteins were analyzed using STRING database for Protein-Protein Interaction Networks and Functional Enrichment Analysis, providing insights into key molecular pathways underlying age-associated differences in zebrafish caudal fin regeneration. The data generated from the study were analysed using Python Program for visualization. Heatmaps and hierarchical clustering were generated using Heatmapper portal (www.heatmapper.ca) to visualize expression dynamics across time points.

### Data submission and Statistical Analysis

The Proteomic data generated both from LFQ and iTRAQ were submitted to the Proteome Xchange consortium via the PRIDE partner repository with the dataset Accession No. PXD074633 and No. PXD074680 respectively. All data are presented as mean ± SEM. Statistical analyses were performed using Student’s t-test. A p-value < 0.05 was considered statistically significant.

## Results

### Age-associated attenuation of caudal fin regeneration in zebrafish

Morphological assessment of caudal fin regeneration revealed clear differences in regenerative capacity among young (<12 months), aged (>36 months), and rotenone-treated zebrafish across different time points post-amputation (Figure 1). In young fish, rapid regenerative outgrowth was observed, with wound closure and initiation of blastema formation occurring as early as 1–2 days post-amputation (dpa). By 7dpa, significant regrowth was evident, characterized by restoration of both lobes and clefts of the fin. In contrast, aged zebrafish displayed markedly reduced regenerative progression, with slower wound healing and limited blastema formation. At 7dpa, fins from aged fish showed incomplete regrowth and reduced structural recovery compared to young fish at 7dpa. Rotenone-treated zebrafish exhibited impairment in regeneration. Mitochondrial inhibition resulted in delayed wound closure and minimal regenerative outgrowth during early stages, with significantly reduced fin length and incomplete structural restoration at later stages. Quantitative measurements of regenerated fin length in both lobe and cleft regions confirmed these observations. Young zebrafish displayed significantly greater regenerative indices across later time points, whereas aged and rotenone-treated fish showed reduced regenerative growth (Figure 1). These findings demonstrate that aging and mitochondrial dysfunction both negatively influence regenerative efficiency in zebrafish caudal fins.

**Figure 1.**
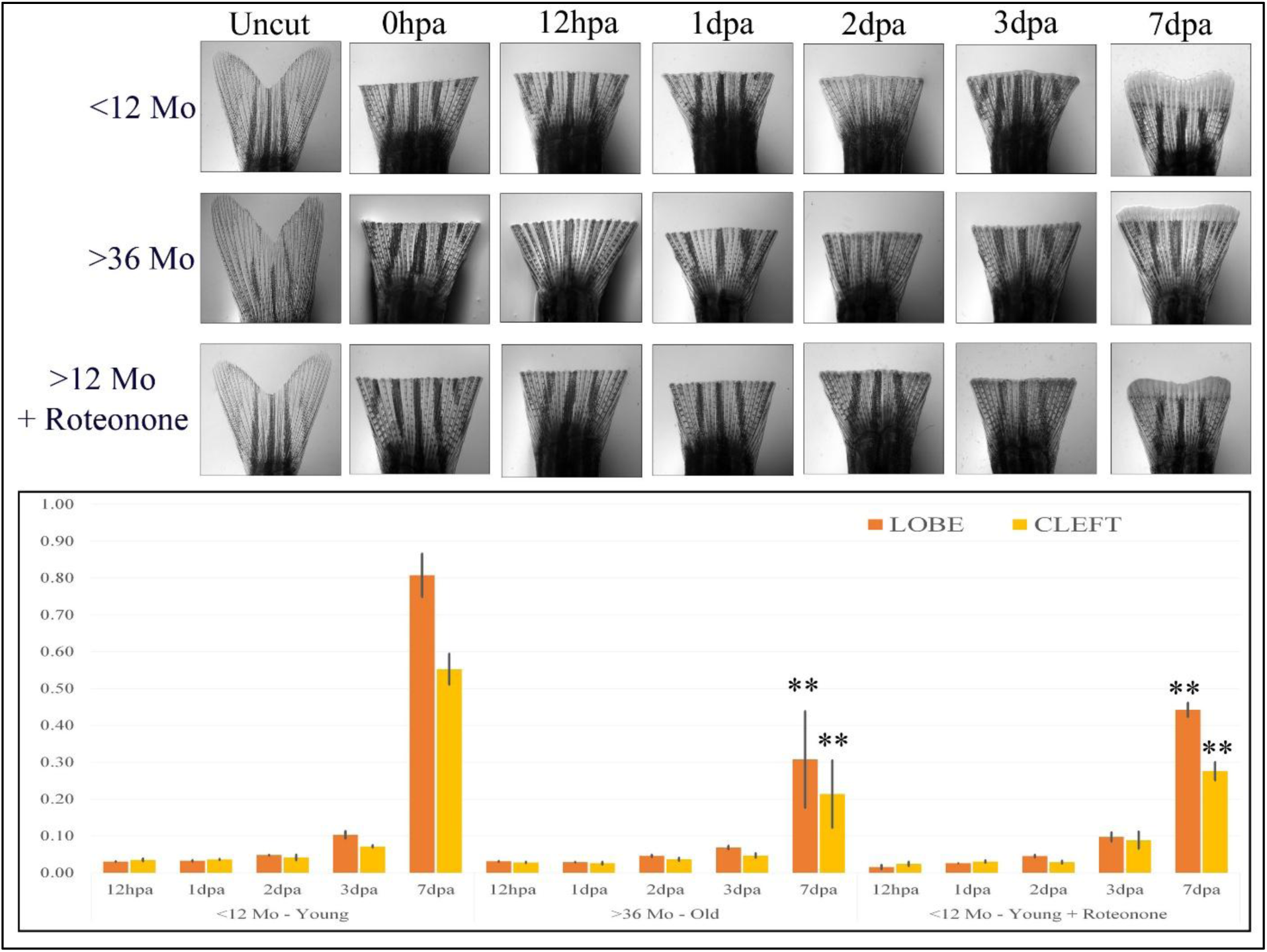
Caudal fin regeneration dynamics in zebrafish. (a) Representative images showing caudal fin regeneration in young (<12 months), old (>36 months), and rotenone-treated young zebrafish at 0 hours post-amputation (hpa; control), 12 hpa, and 1, 2, 3, and 7 days post-amputation (dpa). (b) Quantitative analysis of regenerated fin length in the lobe and cleft regions at 12 hpa and 1, 2, 3, and 7 dpa across experimental groups. Data are presented as mean ± SEM. Statistical significance is indicated (*p* < 0.01).

### Behavioural performance is impaired in aged zebrafish during regeneration

To evaluate the functional impact of aging and regeneration on locomotion, we performed a Novel Tank Test (NTT) to assess exploratory behaviour, including total distance travelled and average velocity, in young and old zebrafish (Figure 2). In control (unamputated) groups, young zebrafish exhibited significantly higher locomotor activity compared to aged fish. Following fin amputation, both groups showed an initial fluctuation in activity at early time points (0h–12h post-amputation), reflecting the immediate impact of injury. However, old zebrafish exhibited reduced distance travelled and lower velocity during intermediate time points (1d—2d post-amputation), and demonstrated a progressive recovery of swimming performance from 2dpa onwards, regaining activity levels comparable to controls by 7dpa. In contrast, young zebrafish exhibited persistently stable distance travelled and lower velocity throughout all the time points, with only fluctuation during intermediate time points. These results indicate that aging has a measurable impact on locomotor recovery following caudal fin amputation, with older fish showing a delayed but eventual restoration of swimming performance, while younger fish maintained relatively stable activity patterns across the regenerative timeline. Since the NTT is also widely used as an indicator of anxiety-like behavior in zebrafish [26], the observed fluctuations in locomotor parameters, particularly reduced exploratory activity in aged fish during early and intermediate phases of regeneration may further reflect an age-related increase in stress or anxiety responses following injury. Thus, the behavioral data suggest that regeneration is not only a structural process but also functionally linked to the modulation of stress and exploratory responses, both of which are influenced by aging.

**Figure 2.**
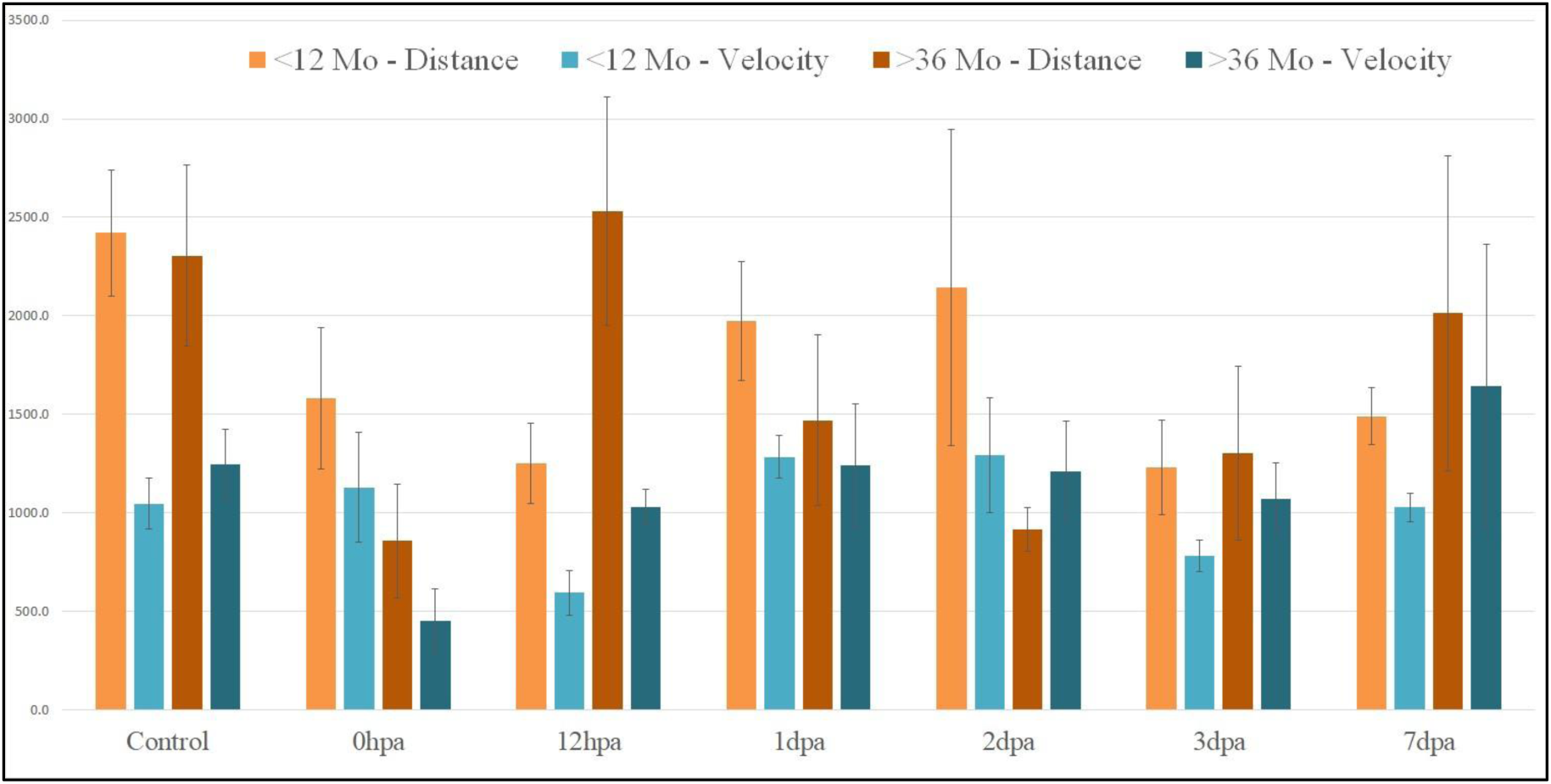
Behavioral assessment during caudal fin regeneration. Bar graphs showing total distance moved (cm) and average velocity (cm/s) during a 2-minute Novel Tank Test in young and old zebrafish at control (0 hpa) and subsequent regeneration time points. The y-axis represents distance moved and velocity. Data are presented as mean ± SEM. Statistical significance is indicated (*p* < 0.01).

### Differential gene expression highlights age-associated molecular changes during fin regeneration

To investigate the molecular basis of impaired regeneration in aged zebrafish, we analyzed the expression of key gene families known to regulate regeneration, including SOX, HOX, MSX, MSX interacting partners, Keratin, Kinase, Laminin, TMEM, Collagen, Solute Carrier (SLC), Neuronal genes, Aging-associated genes, and mitochondrial genes. Heatmap clustering and radial distribution analysis (Figure 3 and Table 1) revealed distinct transcriptional profiles between young and old zebrafish across regeneration time points. In young fish, dynamic upregulation of regenerative markers such as *hoxa13b*, *msxb*, *sox9a*, *keratin*, and *collagen* genes was observed, consistent with active tissue remodeling and blastema formation. In contrast, aged zebrafish displayed attenuated or delayed expression of these genes, particularly within the HOX and MSX families, suggesting disrupted re-patterning and progenitor maintenance.

**Figure 3.**
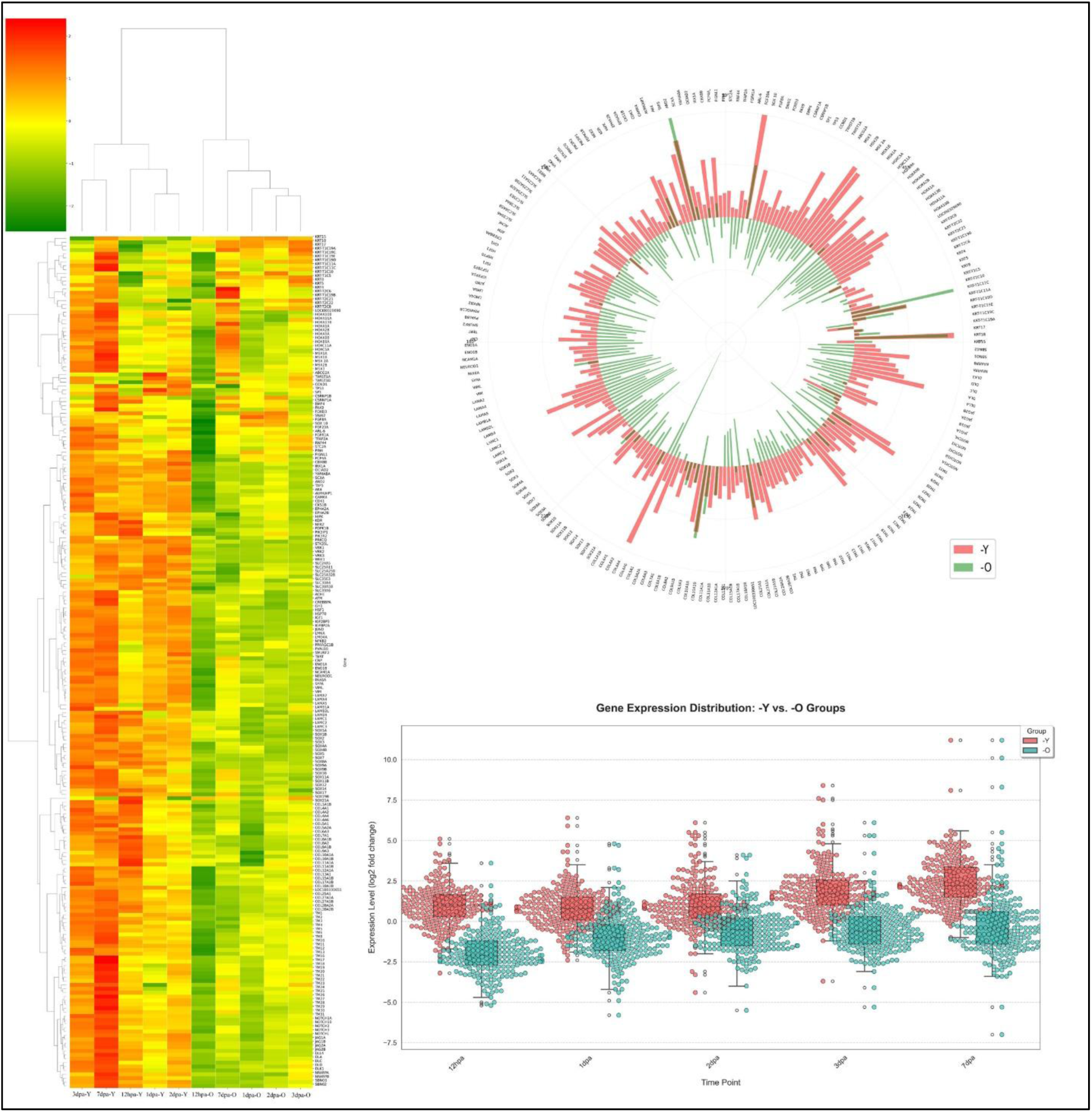
Differential gene expression during caudal fin regeneration. Comparative gene expression analysis of regeneration-associated genes at 12 hpa and 1, 2, 3, and 7 dpa relative to control (0 hpa) in young (<12 months) and old (>36 months) zebrafish. (a) Heatmap of gene expression profiles. (b) Polar plot representation. (c) Swarm plot visualization. Analyses were performed using R software.

**Table 1:** List of Genes validated based on Quantitative real-time PCR.

| S.No | Genes | 12hpa - Y | 1dpa - Y | 2dpa - Y | 3dpa - Y | 7dpa - Y | 12hpa - O | 1dpa - O | 2dpa - O | 3dpa - O | 7dpa - O |
| --- | --- | --- | --- | --- | --- | --- | --- | --- | --- | --- | --- |
| 1 | KRT15 | -3.2 | -0.9 | -0.7 | -0.5 | 0.9 | -1.0 | 1.3 | 0.6 | 1.6 | 1.3 |
| 2 | KRT18 | 3.7 | 6.4 | 5.5 | 5.7 | 3.6 | 2.2 | 4.8 | 3.9 | 5.3 | 3.8 |
| 3 | KRT17 | -1.9 | -0.6 | 1.0 | 0.7 | 2.0 | -1.2 | 1.4 | 1.4 | 2.2 | 1.7 |
| 4 | KRT-T1C19A | -1.5 | 0.6 | 0.9 | 0.5 | 1.6 | -1.2 | 0.8 | 0.4 | 1.7 | 0.4 |
| 5 | KRT-T1C19C | -2.2 | -0.4 | -4.4 | 0.5 | -0.1 | -0.5 | -2.3 | -0.4 | 0.4 | -0.6 |
| 6 | KRT-T1C19E | -0.8 | -1.5 | 0.3 | 1.1 | 1.9 | -2.0 | -2.1 | -2.8 | 0.3 | 0.7 |
| 7 | KRT-T1C19D | -0.5 | 0.8 | 0.6 | 0.4 | 2.1 | 1.0 | 1.7 | 1.2 | 2.5 | 2.5 |
| 8 | KRT-T1C11A | 0.2 | 3.8 | 4.0 | 3.8 | 2.1 | 0.8 | 2.4 | 2.1 | 3.0 | 3.5 |
| 9 | KRT-T1C11C | -0.4 | 0.5 | 2.0 | 1.4 | 3.8 | 3.6 | 4.7 | 4.1 | 5.3 | 4.8 |
| 10 | KRT-T1C10 | 1.3 | 1.6 | 0.8 | 0.8 | 1.1 | -0.9 | 0.4 | 0.2 | 0.0 | -1.2 |
| 11 | KRT-T1C5 | -1.2 | 0.7 | -0.2 | -0.1 | 2.7 | -2.7 | 0.9 | 0.4 | 1.6 | 0.2 |
| 12 | KRT8 | -0.4 | 1.9 | 0.9 | 2.3 | 1.8 | -1.4 | 0.4 | -0.7 | 0.1 | 0.5 |
| 13 | KRT5 | -1.8 | -0.2 | 0.3 | 0.8 | 0.9 | -2.5 | 0.2 | -0.3 | 0.7 | 0.3 |
| 14 | KRT4 | -0.4 | 2.3 | 1.4 | 0.6 | 1.2 | -0.9 | 0.7 | 0.2 | 0.5 | -0.4 |
| 15 | KRT-T2C6 | -1.3 | 1.2 | 1.2 | 0.4 | 1.5 | -2.4 | -0.3 | -1.4 | -0.1 | -1.1 |
| 16 | KRT-T1C19B | -0.9 | -0.7 | 0.8 | 0.5 | 1.7 | -1.9 | -0.4 | 0.8 | 0.9 | 0.6 |
| 17 | KRT-T2C21 | -1.7 | -1.7 | -1.6 | -0.1 | 1.7 | -3.5 | -2.0 | -1.7 | -0.8 | -0.4 |
| 18 | KRT-T2C22 | -0.7 | 0.3 | 0.4 | 0.3 | 2.8 | -2.3 | 0.7 | -0.2 | 0.9 | 0.3 |
| 19 | KRT-T2C8 | 0.7 | 0.8 | 2.2 | 2.3 | 3.3 | -3.4 | -4.2 | -1.9 | -2.4 | -1.8 |
| 20 | LOC100329690 | 1.9 | 2.3 | 2.1 | 2.1 | 2.4 | -2.8 | -1.9 | -1.5 | -1.4 | -1.7 |
| 21 | HOXA10B | 1.0 | 1.9 | 3.1 | 4.0 | 4.6 | -4.3 | -0.4 | -1.0 | -1.2 | 1.1 |
| 22 | HOXA11A | 1.5 | 2.7 | 4.1 | 4.3 | 4.4 | -4.0 | -1.0 | -1.6 | -1.7 | -0.4 |
| 23 | HOXA13B | 1.0 | 1.8 | 2.7 | 2.6 | 3.1 | -3.3 | -2.0 | -2.5 | -2.3 | 0.0 |
| 24 | HOXA1A | 1.4 | 2.5 | 3.2 | 3.4 | 4.0 | -5.0 | -1.6 | -2.0 | -2.5 | -0.7 |
| 25 | HOXA2B | 0.7 | 1.8 | 2.6 | 2.4 | 2.5 | -4.3 | -0.5 | -1.3 | -1.8 | -0.3 |
| 26 | HOXA9A | 0.7 | 1.5 | 2.6 | 2.4 | 2.8 | -3.3 | -1.1 | -1.6 | -1.4 | -0.3 |
| 27 | HOXA9B | 1.3 | 2.3 | 3.2 | 3.8 | 4.5 | -3.8 | -1.3 | -1.5 | -2.1 | 0.0 |
| 28 | HOXB9A | -0.8 | 3.1 | 0.8 | 0.2 | 0.6 | -1.3 | 0.1 | 0.1 | 1.0 | 2.1 |
| 29 | HOXC11A | 1.1 | 2.0 | 2.5 | 2.4 | 3.2 | -3.2 | -2.2 | -2.4 | -2.6 | -1.3 |
| 30 | HOXC5A | 1.3 | 3.5 | 4.8 | 4.2 | 4.7 | -3.9 | -4.9 | -5.5 | -2.6 | -0.6 |
| 31 | MSX1A | 1.1 | 1.6 | 3.2 | 2.1 | 1.5 | -2.9 | 2.1 | 1.0 | 0.3 | 0.5 |
| 32 | MSX1B | 0.9 | 1.2 | 0.4 | 2.8 | 4.9 | -1.3 | -2.2 | -2.2 | -1.3 | 2.4 |
| 33 | MSX 2A | 2.8 | 4.3 | 5.3 | 5.4 | 4.6 | -1.2 | -0.9 | -1.2 | -1.4 | 0.2 |
| 34 | MSX2B | 2.2 | 4.3 | 5.5 | 6.0 | 5.3 | -1.6 | -1.1 | -1.5 | -1.5 | 0.1 |
| 35 | MSX3 | 0.6 | 1.9 | 3.0 | 3.6 | 3.4 | -2.0 | -0.4 | -0.8 | -0.8 | 1.3 |
| 36 | ABCG2A | -0.4 | 1.2 | 0.8 | 1.4 | 2.6 | -1.3 | -0.9 | -1.9 | -1.2 | -1.2 |
| 37 | TWIST1A | 1.4 | 0.6 | 0.9 | 2.6 | 2.8 | -2.5 | -1.0 | -1.4 | -1.1 | -0.3 |
| 38 | TWIST1B | 4.1 | 2.8 | 3.1 | 4.3 | 3.6 | -2.4 | -0.7 | -1.0 | -0.5 | 0.6 |
| 39 | CCND1 | 1.7 | 0.5 | 0.7 | 1.9 | 1.3 | -2.3 | -0.3 | -1.8 | -1.5 | -0.7 |
| 40 | TP53 | -0.1 | 1.2 | 0.5 | 1.4 | 2.1 | -1.8 | -0.2 | -1.8 | -1.0 | -0.8 |
| 41 | SP1 | 0.5 | 1.4 | 0.8 | 1.2 | 2.4 | -1.1 | -0.7 | -1.7 | -0.8 | -0.4 |
| 42 | CSRNP1B | 1.5 | 1.5 | 1.0 | 1.0 | 0.6 | -2.8 | -1.1 | -1.6 | -1.9 | -2.0 |
| 43 | CSRNP1A | 1.8 | 0.1 | -0.2 | 1.2 | 1.4 | -2.5 | -2.1 | -2.6 | -2.3 | -2.4 |
| 44 | BMP4 | 0.2 | 1.3 | 1.0 | 2.8 | 3.9 | -0.8 | -0.6 | -1.7 | -0.2 | 1.3 |
| 45 | PAX9 | 0.3 | 0.6 | 0.5 | 1.2 | 1.7 | -0.9 | -1.1 | -1.8 | -1.2 | -1.1 |
| 46 | FOXD3 | 1.9 | 1.3 | -0.2 | 1.8 | 0.3 | -2.7 | -2.3 | -1.7 | -2.1 | -2.0 |
| 47 | SNAI2 | 1.2 | 0.8 | 0.3 | 1.3 | 1.9 | -2.3 | -0.9 | -2.3 | -1.8 | -1.2 |
| 48 | FGF8A | 1.3 | 1.4 | 2.5 | 3.1 | 4.9 | -5.2 | -1.5 | -1.4 | -1.5 | 0.6 |
| 49 | FGF20A | 4.2 | 5.9 | 5.3 | 7.6 | 4.2 | 0.9 | 1.1 | 2.3 | 4.2 | 3.5 |
| 50 | ARL-6 | 0.5 | 1.0 | 0.8 | 1.3 | 1.2 | -0.2 | -0.9 | -1.0 | -1.1 | -0.1 |
| 51 | FGFR1A | 0.9 | 1.4 | 0.9 | 1.6 | 2.2 | -1.1 | -1.8 | -1.3 | -1.7 | -1.1 |
| 52 | TFAP2A | 0.4 | 0.4 | 0.1 | 0.9 | 0.4 | -2.1 | -1.3 | -0.5 | -1.3 | -0.8 |
| 53 | RNF44 | 0.5 | 0.3 | -0.4 | 0.9 | 1.2 | -2.6 | -2.2 | -1.4 | -1.8 | -1.1 |
| 54 | STC2A | 2.1 | 1.5 | 0.5 | 1.1 | 1.6 | -0.6 | -0.4 | 0.1 | -1.1 | 0.6 |
| 55 | PIN4 | 1.3 | 1.1 | 1.1 | 0.9 | 1.8 | -1.1 | -0.9 | -0.1 | -0.4 | 1.0 |
| 56 | FIGNL1 | 1.1 | 0.2 | 1.7 | 1.6 | 1.0 | -2.8 | 0.3 | 0.4 | -1.4 | -1.1 |
| 57 | PCP4A | 0.8 | 1.8 | 3.3 | 4.9 | 4.6 | -3.1 | -0.7 | 0.4 | 1.0 | 5.5 |
| 58 | CBX8B | 2.0 | 0.5 | 0.6 | 0.8 | 1.6 | -1.4 | -0.1 | 0.1 | -0.3 | -0.4 |
| 59 | IRX1A | 3.3 | 2.0 | 1.6 | 4.0 | 4.2 | -0.8 | 1.1 | 1.4 | 0.6 | -0.1 |
| 60 | OCIAD2 | -0.5 | -0.2 | -0.1 | 1.6 | 2.8 | -3.6 | -2.8 | -1.5 | -1.1 | -1.0 |
| 61 | YWHABA | 0.7 | 0.5 | 0.8 | 2.5 | 2.6 | -2.2 | -1.7 | -0.9 | -0.3 | -0.8 |
| 62 | SCXA | 2.7 | 1.1 | 1.0 | 1.7 | 2.3 | -0.1 | 0.8 | 1.4 | 0.6 | 0.2 |
| 63 | AND2 | 0.7 | 0.0 | 0.9 | 8.4 | 11.2 | 1.3 | 1.7 | 4.1 | 6.1 | 10.1 |
| 64 | TAF5 | -0.1 | 2.8 | 4.2 | 5.6 | 5.4 | -1.5 | -0.7 | -0.6 | 0.3 | -0.4 |
| 65 | AK4 | 0.0 | -0.5 | 0.8 | 1.4 | 3.1 | -2.5 | -2.5 | -1.0 | -1.2 | -0.9 |
| 66 | AURKAIP1 | 1.8 | 0.9 | 2.2 | 2.4 | 3.8 | -1.5 | -0.6 | 0.0 | -0.6 | 0.1 |
| 67 | CAMK4 | -0.2 | -1.0 | -0.1 | 0.2 | 2.3 | -3.3 | -2.2 | -1.0 | -0.6 | -1.4 |
| 68 | CDK1 | 0.6 | 0.7 | 2.7 | 3.9 | 4.9 | -0.3 | 0.5 | 2.0 | 1.6 | 2.8 |
| 69 | CKS1B | 1.3 | 0.9 | 3.2 | 4.1 | 4.8 | -1.7 | 0.2 | 0.9 | 1.2 | 2.1 |
| 70 | EPHA2A | 1.3 | 1.3 | 1.8 | 1.6 | 2.5 | 0.1 | -0.1 | 0.6 | 0.5 | 0.5 |
| 71 | EPHA2B | 0.1 | -0.5 | 0.0 | 0.4 | 2.4 | -1.8 | -1.7 | -1.7 | -1.2 | -0.5 |
| 72 | HIPK | -0.3 | -0.5 | 0.1 | 0.8 | 2.9 | -2.0 | -0.8 | 0.4 | 0.8 | 0.1 |
| 73 | KDR | 0.1 | -0.2 | 0.8 | 1.4 | 2.8 | -1.8 | -0.8 | -0.4 | 0.1 | 1.0 |
| 74 | NEK2 | 0.5 | -0.2 | 1.6 | 1.8 | 4.1 | -2.0 | -1.1 | 0.3 | 0.4 | 0.6 |
| 75 | PDPK1B | -0.2 | -0.2 | 0.3 | 0.5 | 2.7 | -1.8 | -1.2 | -0.7 | -0.7 | -0.7 |
| 76 | PIK3IP1 | 1.1 | -1.9 | 0.1 | 2.0 | 4.8 | -3.2 | -1.2 | -1.1 | -0.3 | 0.7 |
| 77 | PIK3R2 | 0.6 | 0.4 | 1.4 | 1.7 | 3.8 | -2.2 | -1.9 | -1.0 | -0.9 | -0.3 |
| 78 | PRKCQ | -0.6 | -0.8 | -0.6 | 0.0 | 2.2 | -2.3 | -1.7 | -0.8 | -1.0 | -1.2 |
| 79 | STK35L | 0.8 | -0.4 | 0.2 | 0.9 | 3.0 | -2.7 | -2.5 | -1.7 | -1.8 | -1.2 |
| 80 | VRK1 | 1.2 | 0.1 | 1.9 | 2.1 | 3.9 | -2.2 | -1.0 | 0.2 | -0.1 | 0.4 |
| 81 | VRK2 | 0.5 | -1.3 | 1.0 | 1.5 | 3.5 | -2.3 | -1.2 | -0.1 | -0.8 | -0.2 |
| 82 | VRK3 | 0.5 | 0.7 | 1.2 | 1.5 | 3.8 | -1.1 | -1.2 | 0.0 | 0.0 | 0.5 |
| 83 | WEE1 | 1.9 | 0.5 | 2.2 | 2.3 | 4.0 | -1.7 | -0.8 | 0.2 | -0.3 | 0.1 |
| 84 | SLC24A5 | -1.7 | -2.1 | -3.3 | -0.3 | 1.3 | -2.0 | -1.5 | -0.3 | 0.0 | 0.2 |
| 85 | SLC25A11 | 0.8 | 1.2 | 1.4 | 2.5 | 3.3 | -1.9 | 0.4 | 0.3 | 0.9 | -0.4 |
| 86 | SLC25A25B | 2.1 | 0.8 | 1.0 | 2.1 | 2.4 | -0.5 | -0.6 | 0.1 | 0.4 | -0.2 |
| 87 | SLC25A32B | 1.5 | 0.6 | 1.2 | 2.0 | 2.3 | -1.3 | -0.2 | 0.1 | 0.7 | -0.6 |
| 88 | SLC35E3 | 2.2 | 0.7 | 0.6 | 2.4 | 2.2 | -1.4 | -1.5 | -0.5 | 0.1 | -0.2 |
| 89 | SLC38A4 | -1.0 | -1.3 | 0.1 | 0.6 | 1.2 | -2.6 | -2.0 | -1.8 | -1.0 | -2.5 |
| 90 | SLC38A5B | -0.3 | -1.5 | -0.4 | 0.6 | 1.2 | -3.4 | -1.9 | -2.4 | -0.9 | -2.4 |
| 91 | SLC39A6 | 1.2 | 0.6 | 0.8 | 1.8 | 2.5 | -2.4 | -0.9 | -0.9 | -0.1 | -1.6 |
| 92 | ACHE | 0.7 | -1.6 | 0.2 | 1.3 | 2.3 | -3.1 | -2.0 | -1.7 | -1.0 | -3.2 |
| 93 | ATM | 1.2 | 0.9 | 1.5 | 1.4 | 2.6 | -1.5 | -1.1 | -0.7 | -0.4 | -1.4 |
| 94 | CREBBPA | 2.1 | 0.7 | 1.7 | 1.6 | 3.5 | -1.9 | -1.5 | -0.5 | 0.2 | -1.2 |
| 95 | GH1 | 2.5 | 1.1 | 0.8 | 0.3 | 1.5 | -0.6 | -3.7 | -0.6 | -0.5 | -1.1 |
| 96 | HSF1 | 1.7 | 0.7 | 0.7 | 1.4 | 2.6 | -0.8 | -0.9 | -0.2 | 0.6 | -0.7 |
| 97 | HSP70 | 2.4 | 2.2 | 2.3 | 2.2 | 0.6 | 0.2 | -0.2 | 0.5 | -0.6 | -1.1 |
| 98 | IGF1 | 1.6 | 0.7 | 0.2 | 1.5 | 2.2 | -4.1 | -2.1 | -1.6 | -0.8 | -3.0 |
| 99 | IGF2BP3 | 0.7 | -1.2 | 0.5 | 1.6 | 1.9 | -0.6 | -0.4 | 0.3 | 0.9 | 0.6 |
| 100 | IGFBP2A | 1.7 | 0.1 | -0.4 | 1.4 | 1.6 | -2.6 | -2.0 | -1.6 | -1.1 | -2.8 |
| 101 | JUND | -0.6 | -0.1 | 0.7 | -0.6 | 0.8 | -0.6 | -0.4 | -0.5 | -0.1 | -1.9 |
| 102 | LMNA | 1.2 | 1.1 | 0.6 | 1.9 | 3.1 | -2.2 | -0.6 | 0.0 | 0.2 | -0.7 |
| 103 | LMO4A | 0.9 | -0.6 | -0.2 | 1.3 | 2.1 | -1.9 | -1.0 | -0.3 | 0.2 | -0.4 |
| 104 | NFKB2 | 1.6 | 0.6 | 0.9 | 1.8 | 3.9 | -3.8 | -1.2 | -0.5 | -0.2 | -1.7 |
| 105 | PPARGC1B | 1.5 | 0.0 | -0.1 | 1.2 | 2.0 | -3.1 | -2.1 | -1.5 | -0.9 | -2.7 |
| 106 | PVALB8 | 2.5 | -0.5 | -0.9 | 0.4 | 0.2 | -1.0 | -4.2 | -1.1 | -1.1 | -1.0 |
| 107 | SMURF2 | 1.0 | 0.5 | 0.6 | 1.8 | 2.7 | -2.3 | -1.5 | -0.8 | -0.5 | -1.2 |
| 108 | TERT | 0.2 | 0.5 | 0.0 | 1.4 | 1.9 | -2.3 | -0.5 | -0.6 | -0.8 | -2.3 |
| 109 | CNP | 0.6 | 1.7 | 1.2 | 2.9 | 3.2 | -2.6 | -0.9 | -1.4 | -0.8 | -2.3 |
| 110 | ENO1A | 0.1 | -0.3 | -0.5 | 1.2 | 1.7 | -3.6 | -0.7 | -1.4 | -1.5 | -2.6 |
| 111 | ENO1B | 1.7 | 1.7 | 0.0 | 1.7 | 1.9 | -3.4 | -1.5 | -0.1 | 0.6 | -2.1 |
| 112 | NCAM1A | 0.8 | -0.2 | 0.9 | 1.8 | 2.6 | -3.3 | -2.4 | -1.4 | -0.6 | -0.3 |
| 113 | NEUROD1 | 3.1 | 0.9 | -0.8 | 0.1 | 0.5 | -0.9 | -3.4 | -0.1 | -0.5 | -2.0 |
| 114 | PAX6A | 2.7 | 1.0 | 0.0 | 0.9 | 1.3 | -1.7 | -3.4 | -1.8 | -0.5 | -0.8 |
| 115 | SYPA | 3.7 | 1.4 | -2.0 | 0.5 | -1.0 | 1.6 | -0.4 | 1.1 | 2.0 | 1.0 |
| 116 | VIML | 0.9 | -0.3 | 1.0 | 0.0 | 0.6 | -2.5 | -1.7 | -3.0 | -1.2 | -3.4 |
| 117 | VIM | 0.7 | 0.7 | 0.5 | 1.0 | 2.1 | -5.1 | -1.0 | -1.9 | -0.7 | -2.8 |
| 118 | LAMA2 | -0.3 | 0.5 | -0.3 | 1.1 | 2.1 | -3.9 | -3.0 | -2.9 | -3.1 | -0.4 |
| 119 | LAMA4 | -0.7 | 0.2 | 0.0 | 1.0 | 0.8 | -3.1 | -1.8 | -2.5 | -2.0 | -0.4 |
| 120 | LAMA5 | 0.3 | 0.7 | 0.8 | 2.6 | 2.7 | -4.0 | -3.2 | -3.0 | -1.9 | -1.0 |
| 121 | LAMB1A | 1.0 | 0.6 | 1.2 | 3.6 | 4.7 | -3.3 | -1.7 | -0.9 | 0.2 | 1.8 |
| 122 | LAMB2L | 0.7 | 1.6 | 1.8 | 2.3 | 2.3 | -4.4 | -1.0 | -0.9 | -2.4 | -1.5 |
| 123 | LAMB4 | 2.8 | 1.8 | 2.8 | 4.3 | 4.3 | -4.5 | -1.2 | -1.5 | -2.6 | -5.3 |
| 124 | LAMC1 | 0.8 | 1.2 | 2.0 | 2.8 | 2.4 | -3.3 | -1.8 | -1.3 | -1.9 | -0.5 |
| 125 | LAMC2 | 1.0 | 2.0 | 1.4 | 1.6 | 0.7 | -1.8 | -1.9 | -2.3 | -1.6 | -1.3 |
| 126 | LAMC3 | -0.4 | -0.7 | -0.5 | 1.7 | 2.3 | -4.3 | -3.7 | -2.5 | -4.1 | -0.6 |
| 127 | SOX1A | 1.7 | 0.2 | -0.6 | 0.1 | 0.8 | -2.4 | -3.8 | -2.9 | -2.4 | -3.4 |
| 128 | SOX1B | 2.0 | 0.0 | -1.2 | 0.2 | 0.8 | -2.6 | -4.4 | -3.1 | -2.7 | -2.7 |
| 129 | SOX2 | 2.9 | 1.9 | 1.9 | 3.2 | 3.1 | -2.1 | -2.8 | -2.5 | -2.1 | -2.6 |
| 130 | SOX3 | -0.2 | -0.4 | -0.1 | -0.4 | 0.8 | -1.2 | -1.5 | -1.6 | -0.6 | -1.3 |
| 131 | SOX4A | 1.2 | 1.6 | 2.0 | 2.9 | 2.3 | -1.7 | -0.7 | 0.1 | -0.4 | 0.8 |
| 132 | SOX4B | 1.1 | 1.5 | 1.1 | 1.5 | 2.3 | -1.6 | -1.7 | -1.4 | -0.5 | -1.1 |
| 133 | SOX5 | 0.3 | 1.0 | 1.1 | 1.3 | 1.9 | -2.1 | -1.5 | -1.5 | -0.7 | -1.6 |
| 134 | SOX7 | 1.1 | 0.0 | -0.8 | 0.2 | 1.0 | -2.0 | -2.7 | -1.9 | -1.4 | -2.1 |
| 135 | SOX8A | 2.0 | -0.2 | -2.0 | 0.0 | 0.6 | -2.2 | -3.9 | -2.6 | -2.4 | -3.2 |
| 136 | SOX9A | 0.3 | 0.4 | 1.0 | 1.5 | 0.4 | -2.4 | -1.2 | -0.8 | -1.8 | -1.3 |
| 137 | SOX9B | -0.9 | 1.1 | 1.0 | 1.2 | 1.0 | 0.0 | 0.5 | 1.7 | 0.4 | 1.6 |
| 138 | SOX10 | 1.1 | 1.6 | 1.8 | 2.2 | 1.2 | -3.3 | -0.2 | 0.0 | -1.2 | -1.5 |
| 139 | SOX11A | 1.6 | 2.1 | 2.0 | 2.8 | 1.3 | -0.5 | 0.6 | 1.1 | 0.5 | 0.3 |
| 140 | SOX11B | -0.1 | 0.3 | 0.9 | 1.8 | 1.0 | -2.2 | 0.0 | 0.2 | -0.4 | 0.3 |
| 141 | SOX12 | 0.2 | 0.8 | 1.0 | 1.4 | 0.7 | -2.8 | -0.7 | -0.8 | -1.9 | -1.7 |
| 142 | SOX14 | 0.8 | 0.9 | 1.2 | 1.5 | 0.6 | -1.9 | -0.7 | -0.5 | -1.4 | -1.5 |
| 143 | SOX17 | 1.2 | 0.3 | 1.0 | 1.4 | 0.7 | -2.6 | 1.2 | 1.4 | 0.0 | 0.4 |
| 144 | SOX19B | 1.3 | 1.8 | 2.0 | 2.5 | 2.1 | -2.0 | -0.7 | -0.1 | -0.6 | -0.3 |
| 145 | SOX21A | 1.0 | 1.3 | 1.6 | 1.5 | 0.7 | -3.1 | 0.2 | 0.3 | -0.9 | -1.9 |
| 146 | COL1A1B | -0.3 | 1.6 | 1.2 | 3.0 | 3.6 | -2.5 | -1.6 | -0.2 | 0.5 | 4.0 |
| 147 | COL4A1 | 1.4 | 0.9 | 0.8 | 2.0 | 2.1 | -1.8 | 2.0 | 1.8 | 0.1 | 1.1 |
| 148 | COL4A2 | -0.3 | 1.1 | -1.1 | 1.4 | 2.0 | -2.9 | -1.9 | -1.3 | -1.6 | 0.1 |
| 149 | COL4A4 | 5.1 | 4.7 | 6.1 | 3.5 | 5.6 | -3.1 | -5.8 | -4.0 | -5.3 | -7.0 |
| 150 | COL4A6 | -1.1 | 0.8 | 1.4 | 2.4 | 3.4 | -2.4 | -1.9 | -0.7 | 0.0 | 2.0 |
| 151 | COL5A1 | 1.3 | 2.6 | 2.7 | 4.2 | 4.5 | -2.1 | -1.5 | -0.4 | 0.2 | 2.4 |
| 152 | COL5A2A | 0.4 | 0.5 | 0.3 | 2.9 | 3.5 | -1.6 | -1.0 | 0.0 | -0.2 | 2.8 |
| 153 | COL6A3 | 0.2 | 0.1 | 0.1 | 1.0 | 2.1 | -2.8 | 0.2 | 0.4 | -0.7 | 1.4 |
| 154 | COL7A1 | 1.2 | 1.4 | 1.5 | 2.9 | 2.6 | -2.6 | 1.1 | 1.6 | 0.5 | 1.6 |
| 155 | COL8A1B | 1.1 | -0.6 | 1.0 | 1.8 | 1.5 | -2.5 | 3.2 | 2.5 | 1.1 | 0.9 |
| 156 | COL8A2 | 2.1 | 0.9 | 0.7 | 0.6 | 2.2 | -1.2 | 0.7 | 1.0 | -0.2 | 0.9 |
| 157 | COL9A1B | 1.6 | 0.7 | 0.0 | 0.2 | 2.4 | -0.4 | 0.9 | 1.7 | 0.4 | 1.7 |
| 158 | COL9A3 | 0.4 | 1.9 | 4.7 | 4.1 | 3.9 | -2.9 | 0.6 | -1.2 | -1.7 | -1.3 |
| 159 | COL10A1A | 0.7 | 0.8 | 0.3 | 7.5 | 8.1 | -2.5 | 3.3 | 2.9 | 1.4 | 11.2 |
| 160 | COL10A1B | 0.7 | 0.7 | -0.4 | 2.1 | 4.9 | -1.6 | 2.1 | 1.7 | 0.3 | 8.3 |
| 161 | COL11A1A | 1.8 | 1.7 | 2.0 | 2.1 | 3.2 | 0.5 | -0.9 | -0.2 | -0.4 | 1.0 |
| 162 | COL11A1B | 0.3 | 0.9 | 0.3 | 2.6 | 3.3 | -1.0 | 1.3 | 1.1 | 0.8 | 4.3 |
| 163 | COL12A1A | 0.6 | 1.1 | 1.1 | 2.8 | 2.7 | -2.1 | -0.2 | -0.1 | 0.0 | 2.4 |
| 164 | COL13A1 | -0.4 | -0.2 | 0.2 | 1.8 | 3.6 | -3.4 | -2.7 | -0.8 | -1.9 | 1.8 |
| 165 | COL15A1B | 1.2 | 1.6 | 1.2 | 1.4 | 3.1 | 0.2 | -2.2 | -1.7 | -2.5 | -1.6 |
| 166 | COL17A1B | 0.5 | 1.2 | 1.3 | 1.7 | 2.1 | -2.5 | -2.5 | -1.5 | -0.9 | -1.0 |
| 167 | COL18A1B | 0.8 | 0.8 | 0.6 | 0.6 | 2.2 | -0.7 | 0.0 | 1.0 | 0.0 | 1.0 |
| 168 | LOC100330651 | 1.6 | 0.0 | -0.3 | 1.0 | 2.0 | -1.8 | -2.6 | -2.2 | -2.5 | -1.4 |
| 169 | COL25A1 | 1.1 | 0.3 | 1.0 | 2.1 | 4.1 | -1.1 | -0.9 | 0.1 | 0.2 | 2.0 |
| 170 | COL27A1A | 1.5 | 2.4 | 2.1 | 3.3 | 5.0 | 0.1 | 1.0 | 1.6 | 2.4 | 4.5 |
| 171 | COL27A1B | 1.7 | 1.1 | 1.1 | 1.8 | 3.7 | -1.9 | -1.5 | -0.8 | -0.3 | 1.0 |
| 172 | COL28A2A | 1.7 | 0.9 | 1.4 | 1.6 | 4.7 | -2.6 | -1.1 | 0.8 | 0.4 | 1.4 |
| 173 | COL28A2B | 3.6 | 0.9 | 2.5 | 2.8 | 3.4 | 1.0 | -0.1 | -0.1 | 0.8 | 1.4 |
| 174 | TM1 | 2.3 | 0.5 | 0.7 | 2.0 | 1.2 | -2.5 | -1.5 | -0.3 | -0.2 | -0.9 |
| 175 | TM2 | 3.0 | 0.1 | 0.0 | 2.7 | 1.7 | -1.9 | -2.7 | -0.1 | 0.3 | 0.1 |
| 176 | TM3 | 1.6 | 0.7 | 0.2 | 2.8 | 1.5 | -2.1 | -1.7 | -0.8 | -1.1 | -1.3 |
| 177 | TM4 | 1.4 | 0.3 | 0.6 | 2.3 | 1.8 | -2.6 | -0.6 | -0.3 | 0.2 | -0.7 |
| 178 | TM5 | 0.7 | 0.9 | 0.9 | 3.1 | 2.1 | -3.6 | -0.4 | -0.5 | 0.2 | -0.8 |
| 179 | TM6 | 2.5 | 0.2 | -0.2 | 1.7 | 0.6 | -1.1 | -1.4 | -0.1 | -0.5 | -0.4 |
| 180 | TM8 | 1.2 | 0.6 | 1.0 | 2.8 | 2.1 | -3.6 | -0.2 | -0.6 | -0.1 | -1.5 |
| 181 | TM10 | 1.5 | 0.1 | 1.0 | 2.6 | 1.6 | -2.1 | -0.1 | 0.2 | 0.5 | -0.2 |
| 182 | TM11 | 0.9 | 0.2 | 0.7 | 2.7 | 2.3 | -2.6 | 0.2 | 0.3 | 1.1 | 0.6 |
| 183 | TM12 | 1.9 | 0.0 | 0.1 | 2.1 | 0.8 | -1.7 | -1.6 | -0.4 | -0.2 | -0.7 |
| 184 | TM13 | 1.1 | -0.1 | -0.3 | 1.6 | 1.1 | -2.2 | -0.5 | -0.3 | 0.3 | -0.8 |
| 185 | TM16 | 2.1 | 0.5 | 0.5 | 2.0 | 1.3 | -1.8 | -0.4 | -0.1 | 0.8 | 0.0 |
| 186 | TM17 | 0.7 | -0.6 | -0.5 | 1.0 | 0.0 | -3.4 | -1.1 | -1.7 | -1.0 | -1.8 |
| 187 | TM18 | 1.3 | -0.3 | 0.3 | 1.9 | 1.4 | -2.3 | -0.2 | 0.6 | 1.0 | 0.2 |
| 188 | TM19 | 3.8 | 2.8 | 3.5 | 2.8 | 2.6 | -1.7 | 1.1 | 0.6 | 0.9 | 0.6 |
| 189 | TM20 | 1.9 | 1.4 | 1.4 | 3.7 | 4.9 | -2.9 | -2.4 | -0.4 | -0.4 | -1.3 |
| 190 | TM21 | 1.4 | -0.2 | 1.0 | 1.5 | 2.5 | -0.1 | -1.0 | -0.1 | -0.6 | 0.2 |
| 191 | TM22 | 1.2 | 0.4 | 1.1 | 2.1 | 2.1 | 0.2 | 0.0 | 0.3 | 1.1 | 2.7 |
| 192 | TM23 | 1.6 | -0.2 | 1.2 | 1.9 | 3.4 | -2.3 | -2.2 | -1.3 | -0.7 | -1.2 |
| 193 | TM24 | 0.9 | -0.5 | 0.5 | 1.8 | 2.5 | -4.7 | -2.9 | -1.7 | -1.7 | -3.4 |
| 194 | TM25 | 2.0 | 0.9 | 1.0 | 3.4 | 3.7 | -0.4 | -1.4 | 0.7 | 0.8 | 1.2 |
| 195 | TM26 | 2.5 | 1.0 | 1.1 | 2.4 | 2.6 | -0.6 | -1.8 | 0.2 | 0.5 | 0.5 |
| 196 | TM27 | 1.2 | 0.7 | 1.1 | 2.3 | 2.2 | -1.5 | 0.1 | 1.1 | 1.7 | 0.6 |
| 197 | TM28 | 2.9 | 1.4 | 1.5 | 1.5 | 3.5 | -0.8 | -3.1 | -1.5 | -2.6 | -1.0 |
| 198 | TM29 | 4.8 | 2.2 | 3.1 | 4.8 | 4.0 | 0.8 | 0.4 | 0.9 | 1.5 | 1.4 |
| 199 | TM30 | 3.3 | 1.8 | 2.3 | 4.0 | 5.3 | -2.1 | -2.6 | -1.1 | -1.3 | -1.3 |
| 200 | TM31 | -0.9 | -1.2 | 0.2 | 1.6 | 1.9 | -3.2 | -0.8 | -0.2 | 0.4 | -0.4 |
| 201 | NOTCH1A | 0.6 | -0.6 | -0.5 | -0.4 | 0.1 | -1.6 | -1.3 | -1.2 | -1.5 | -1.8 |
| 202 | NOTCH1B | 0.0 | -1.0 | -0.5 | -0.6 | 0.5 | -1.4 | -1.3 | -0.9 | -1.6 | -2.7 |
| 203 | NOTCH2 | -1.0 | -2.4 | -3.1 | -3.7 | -0.3 | -1.3 | -0.5 | -0.4 | -0.9 | -1.2 |
| 204 | NOTCH3 | 1.8 | 1.1 | 1.0 | 0.8 | 1.9 | -1.4 | -1.0 | -0.5 | -0.8 | -1.4 |
| 205 | NOTCHL | 1.9 | 0.9 | 2.3 | 2.0 | 1.9 | -2.9 | -1.5 | -0.3 | -1.7 | -1.6 |
| 206 | JAG1A | 0.5 | 1.4 | 1.4 | 0.8 | 1.4 | -0.7 | -0.3 | -0.2 | -0.3 | -1.2 |
| 207 | JAG1B | 0.2 | 0.1 | -0.2 | -1.2 | 0.1 | -1.3 | -0.9 | -1.1 | -1.5 | -1.8 |
| 208 | JAG2A | 1.1 | 0.1 | 1.0 | 0.9 | 2.2 | -0.7 | -0.8 | 0.2 | -1.2 | -1.7 |
| 209 | JAG2B | 0.3 | 0.7 | 0.5 | -0.2 | 0.6 | -1.7 | -1.9 | -1.3 | -1.7 | -3.0 |
| 210 | DLL4 | -0.5 | 1.3 | 1.6 | 1.3 | 2.2 | -2.4 | -1.3 | -0.2 | -0.8 | -1.7 |
| 211 | DLA | 2.1 | 2.5 | 5.1 | 5.3 | 3.7 | -3.1 | -2.0 | -0.2 | -1.3 | -1.7 |
| 212 | DLC | 2.1 | 2.4 | 3.3 | 2.4 | 3.1 | -2.7 | -1.9 | -0.6 | -2.0 | -1.5 |
| 213 | DLD | 1.0 | 2.5 | 3.7 | 2.2 | 2.2 | -2.6 | -1.2 | 0.0 | -1.2 | -0.7 |
| 214 | DLK1 | 0.0 | 2.0 | 2.7 | 2.6 | 3.4 | -1.2 | -0.1 | 1.0 | 0.1 | 0.2 |
| 215 | NRARPA | 1.5 | 1.7 | 2.6 | 1.4 | 1.7 | -1.3 | -1.5 | -1.3 | -1.5 | -1.6 |
| 216 | NRARPB | 1.9 | 2.9 | 3.7 | 2.3 | 3.1 | -2.3 | -2.1 | -1.5 | -1.9 | -2.1 |
| 217 | SBN01 | 1.1 | 1.9 | 0.8 | 0.7 | 2.1 | -1.4 | -1.6 | -1.1 | -1.3 | -1.7 |
| 218 | SBN02 | 0.9 | 1.7 | 0.8 | 0.2 | 1.7 | -1.2 | -1.8 | -1.5 | -1.6 | -1.8 |
| 219 | CO1 | 1.3 | 1.6 | 1.6 | 1.8 | 3.5 | -1.6 | -1.1 | -0.4 | -0.2 | -1.0 |
| 220 | CO2 | 1.5 | 1.8 | 2.2 | 2.4 | 3.7 | -1.6 | -1.4 | -0.2 | 0.0 | -0.6 |
| 221 | CYB | 1.6 | 1.8 | 2.6 | 2.5 | 4.1 | -1.1 | -1.1 | 0.3 | 0.6 | -0.2 |
| 222 | ND1 | 1.8 | 2.0 | 2.3 | 2.5 | 4.1 | -0.7 | -0.3 | 1.2 | 0.8 | 0.1 |
| 223 | ND4 | 1.4 | 1.7 | 2.1 | 2.1 | 3.8 | -1.3 | -1.1 | 0.1 | 0.2 | -0.3 |
| 224 | ND4L | 1.4 | 2.0 | 2.4 | 2.4 | 4.2 | -1.7 | -1.3 | 0.0 | 0.0 | -0.3 |
| 225 | ND5 | 1.2 | 1.8 | 2.0 | 1.9 | 3.8 | -1.3 | -0.8 | 0.4 | 0.2 | -0.2 |
| 226 | ND6 | 1.4 | 1.5 | 2.4 | 2.1 | 3.7 | -1.3 | -0.6 | 0.9 | 0.5 | -0.1 |

Meanwhile, aging-associated genes such as *igf1*, *tert*, *hsp70*, and *atm* showed reduced induction in older fish, reflecting compromised growth signaling, genomic maintenance, and stress resilience. Similarly, transport- and signaling-related genes such as *slc25a1a*, *slc25a1b*, and *tmem9* were differentially expressed, further implicating impaired metabolic and membrane homeostasis in age-related regenerative decline. These molecular differences align with the morphological and behavioral findings, highlighting that age-associated attenuation of regenerative signaling networks contributes to reduced fin regrowth and functional recovery in zebrafish.

Notably, mitochondrial genes encoding electron transport chain components (*mt-nd1, mt-nd4, mt-nd4L, mt-nd5, mt-nd6, mt-co1, mt-co2,* and *mt-cyb*) exhibited dynamic expression patterns during regeneration in young zebrafish (Figure 3 and 4). The prominent regulation of these mitochondrial genes during regenerative progression prompted us to investigate the functional role of mitochondrial activity in this process. To examine this, mitochondrial function was pharmacologically inhibited using rotenone, a Complex I inhibitor. Following rotenone treatment, these mitochondrial genes displayed reduced or dysregulated expression patterns similar to those observed in aged zebrafish, suggesting that mitochondrial dysfunction contributes to impaired regenerative responses.

**Figure 4.**
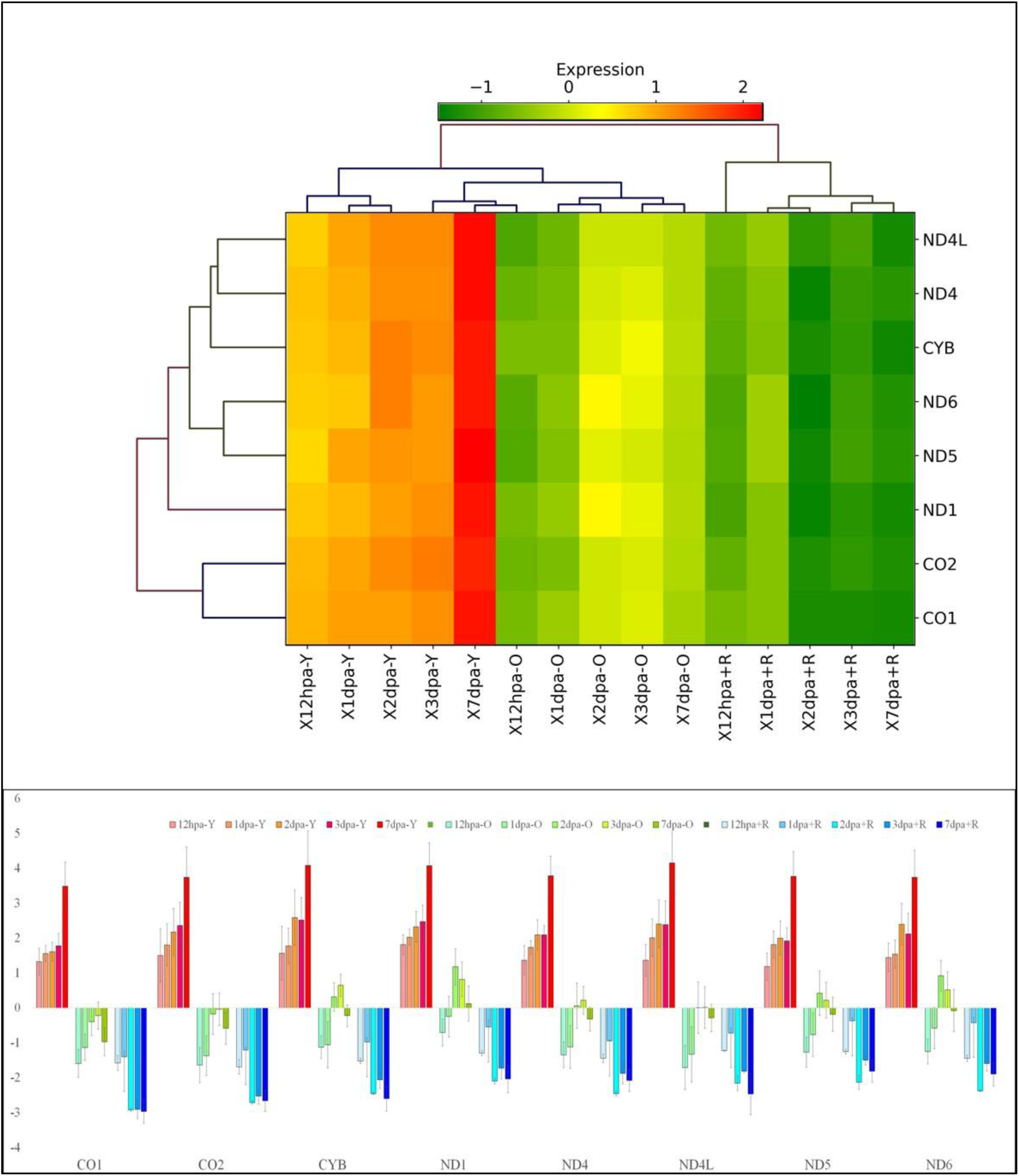
Differential expression of mitochondrial genes during regeneration. Comparative gene expression analysis of mitochondrial genes in young (<12 months), old (>36 months), and rotenone-treated young zebrafish (a) Heatmap showing expression profiles of mitochondrial genes associated with caudal fin regeneration. (b) Bar graph representation of differential gene expression.

### STRING network analysis reveals functional interactions and enrichment of regenerative gene families

To characterize the functional relationships among the differentially expressed regenerative genes, STRING-based protein-protein interaction and functional enrichment analysis was performed using the selected gene families (Figure 5). The STRING network revealed a highly interconnected interaction map linking transcription factors, signaling molecules, extracellular matrix components, transporters, and aging-associated genes, underscoring the coordinated molecular framework underlying caudal fin regeneration.

**Figure 5.**
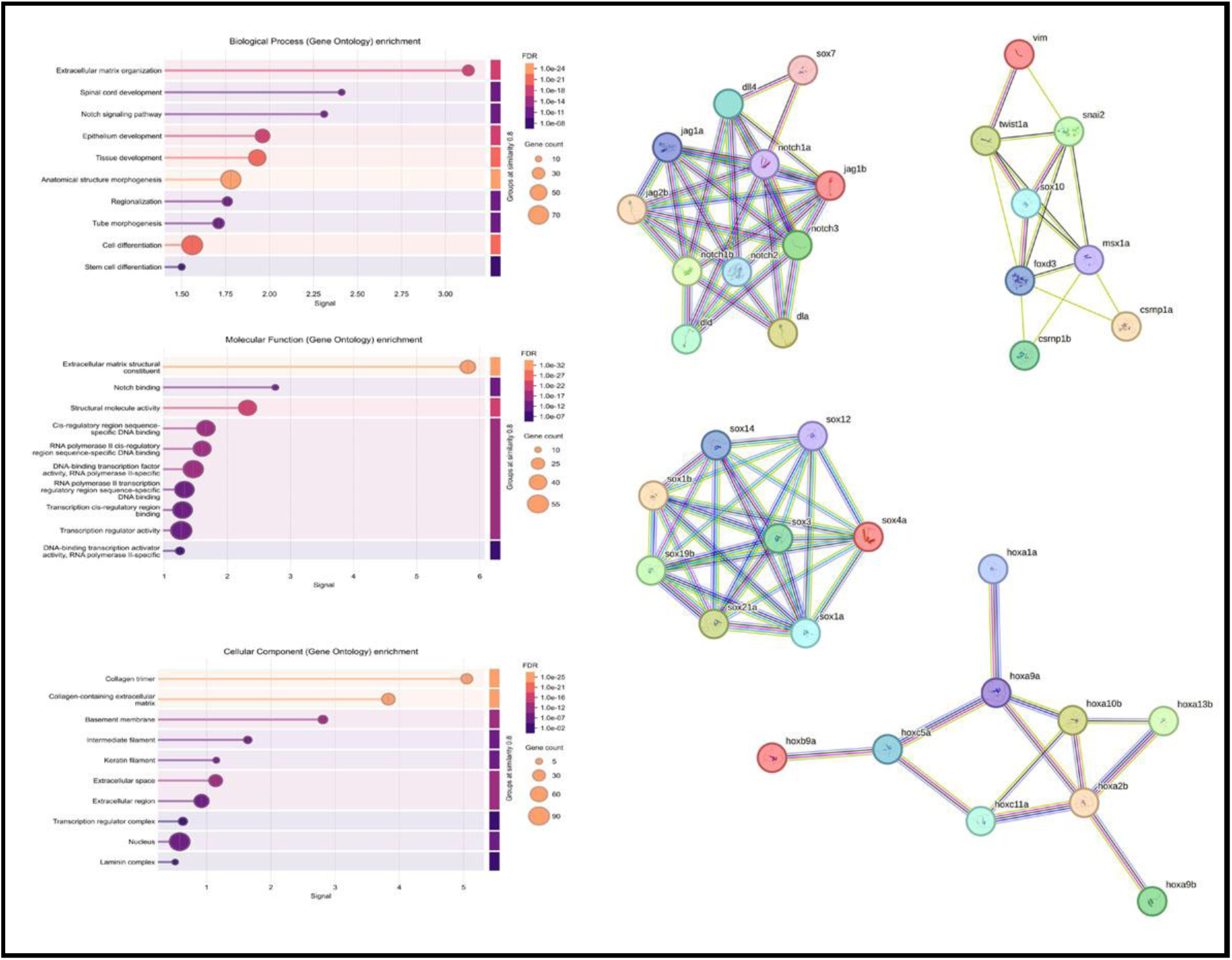
Gene network and functional enrichment analysis. Functional classification and network analysis of genes associated with caudal fin regeneration. (a) Biological processes. (b) Molecular functions. (c) Cellular components. (d) Interaction networks derived from the selected gene set.

Functional enrichment analysis highlighted significant clustering of genes involved in key biological processes, including developmental patterning, regulation of transcription, cell differentiation, extracellular matrix organization, cell proliferation, and response to stress. Prominent transcriptional regulators such as HOX, MSX, and SOX family members were strongly associated with processes related to morphogenesis, positional identity, and progenitor cell regulation. Notch signaling components and kinase-associated genes were enriched in pathways governing cell fate determination and signal transduction, emphasizing their regulatory role during regenerative progression.

Analysis of molecular function enrichment revealed overrepresentation of DNA-binding transcription factor activity, protein kinase activity, structural molecule activity, and transporter activity. Genes encoding keratins, collagens, and laminins clustered within structural and extracellular matrix–related functions, reflecting their role in tissue integrity, wound epidermis formation, and regenerative outgrowth. Solute carrier (SLC) family members were enriched for transmembrane transporter activity, highlighting their contribution to metabolic support and ionic homeostasis during regeneration.

At the level of cellular components, enriched terms included nucleus, cytoplasm, plasma membrane, extracellular matrix, and cell junctions, indicating the spatial distribution of regenerative gene activity across intracellular, membrane-associated, and extracellular compartments. Aging-associated genes such as *igf1*, *tert*, *hsp70*, and *atm* were linked to pathways involved in growth regulation, proteostasis, telomere maintenance, and DNA damage response, integrating stress resilience and cellular maintenance mechanisms into the regenerative network.

Collectively, the STRING interaction and enrichment analysis demonstrates that the selected gene families participate in an integrated molecular network encompassing transcriptional regulation, signaling cascades, extracellular matrix remodeling, transport processes, and cellular stress responses, all of which are essential for successful caudal fin regeneration.

### Proteomic profiling reveals proteomics reveals age-dependent alterations in regenerative protein dynamics

To investigate age-associated changes in protein expression during caudal fin regeneration, quantitative proteomic analysis was performed using both label-free quantification (LFQ) and iTRAQ-based approaches. Regenerating fin tissues from young and old zebrafish were analyzed at three key regenerative stages: early (1dpa), mid (2dpa), and late (7dpa) against 0hpa as the baseline control. Proteins identified across samples showed robust peptide coverage and high confidence, enabling reliable quantitative comparisons between age groups (Table 2). Notably, all 32 proteins identified across the regenerative stages using the iTRAQ-labeled approach were consistently detected in the LFQ proteomic dataset as well, demonstrating strong concordance between the two independent quantification methods. This overlap highlights the robustness and reproducibility of the proteomic analysis and strengthens confidence in the identified regeneration-associated protein signatures.

**Table 2:**
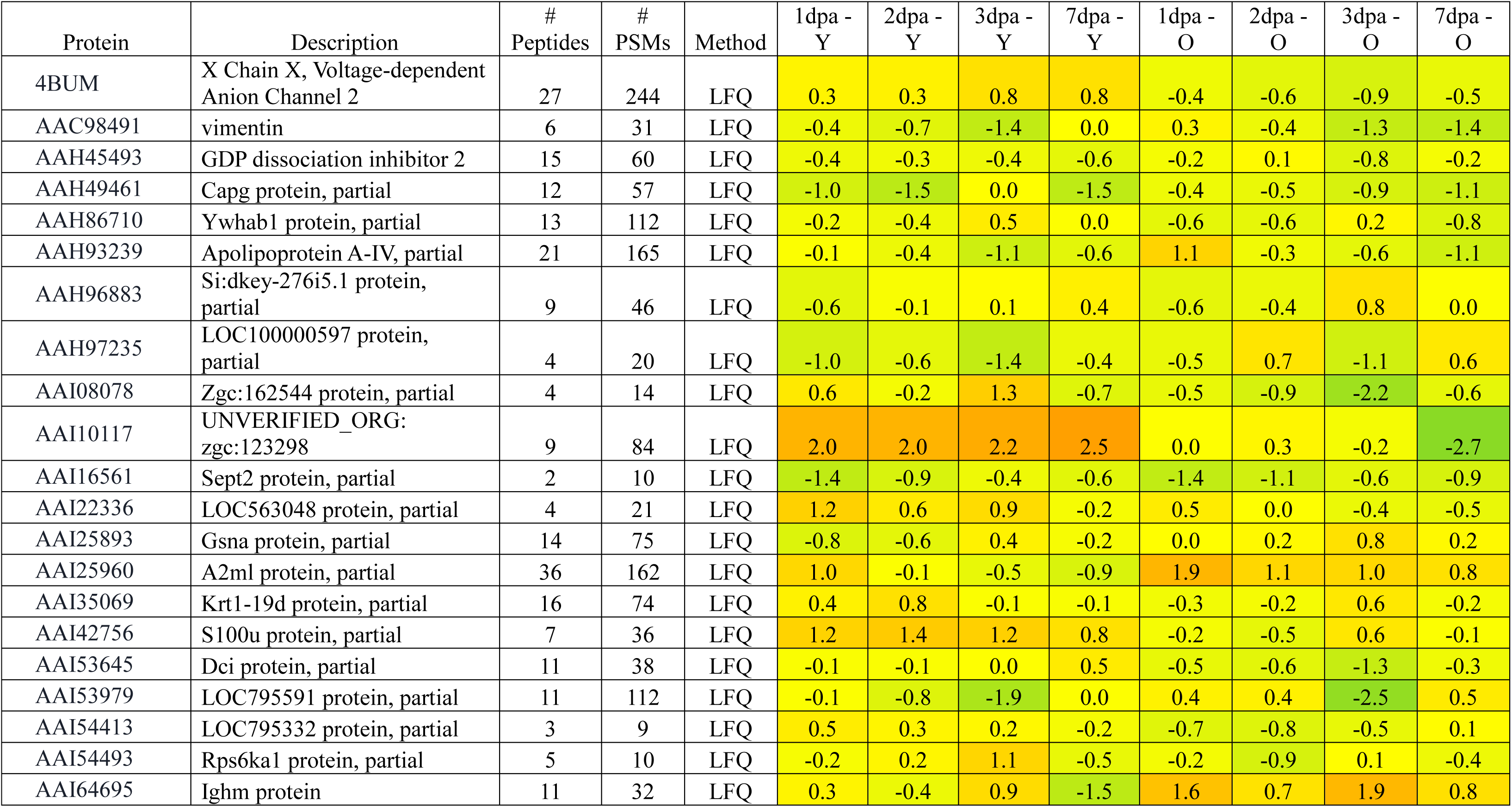
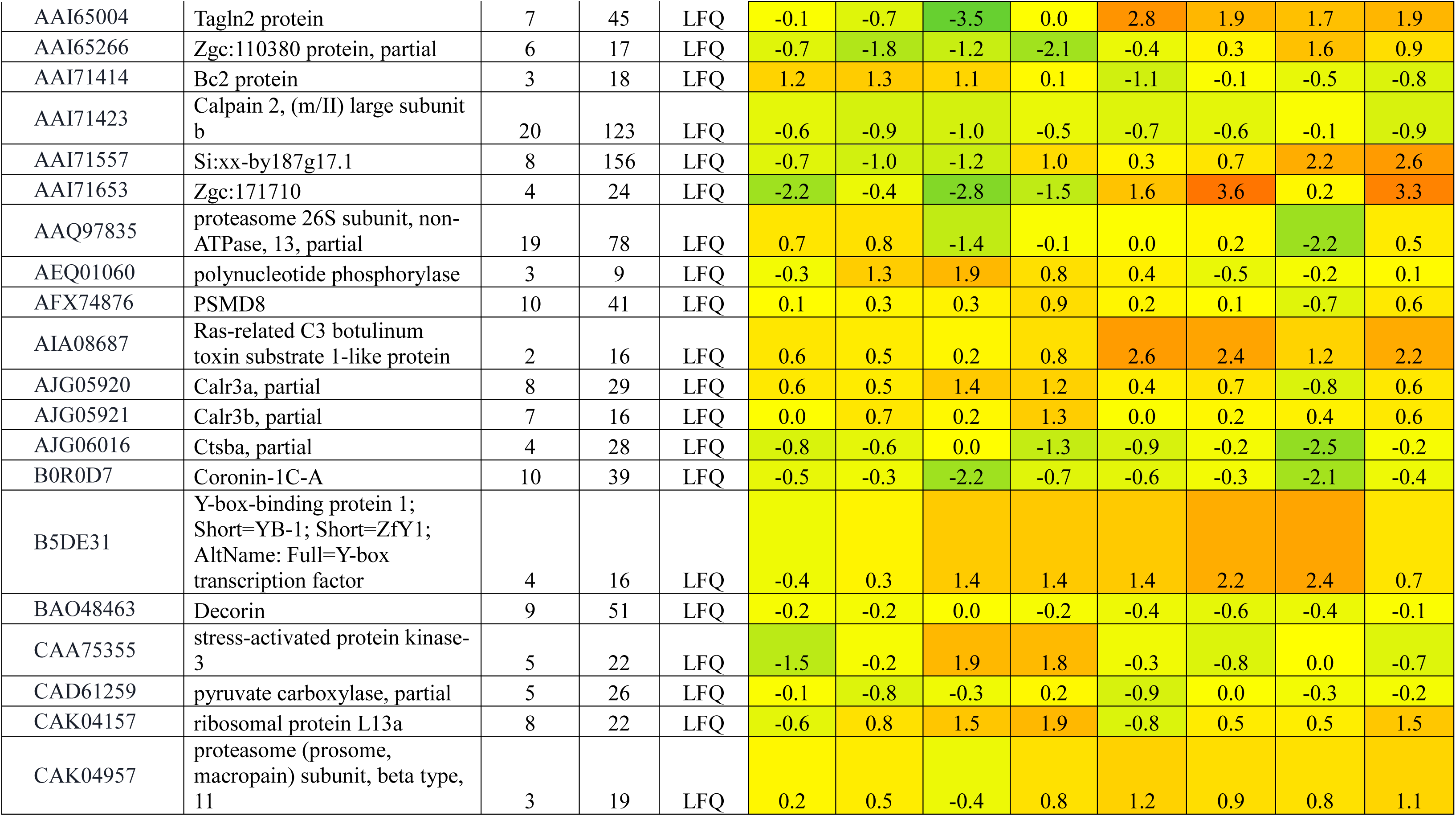
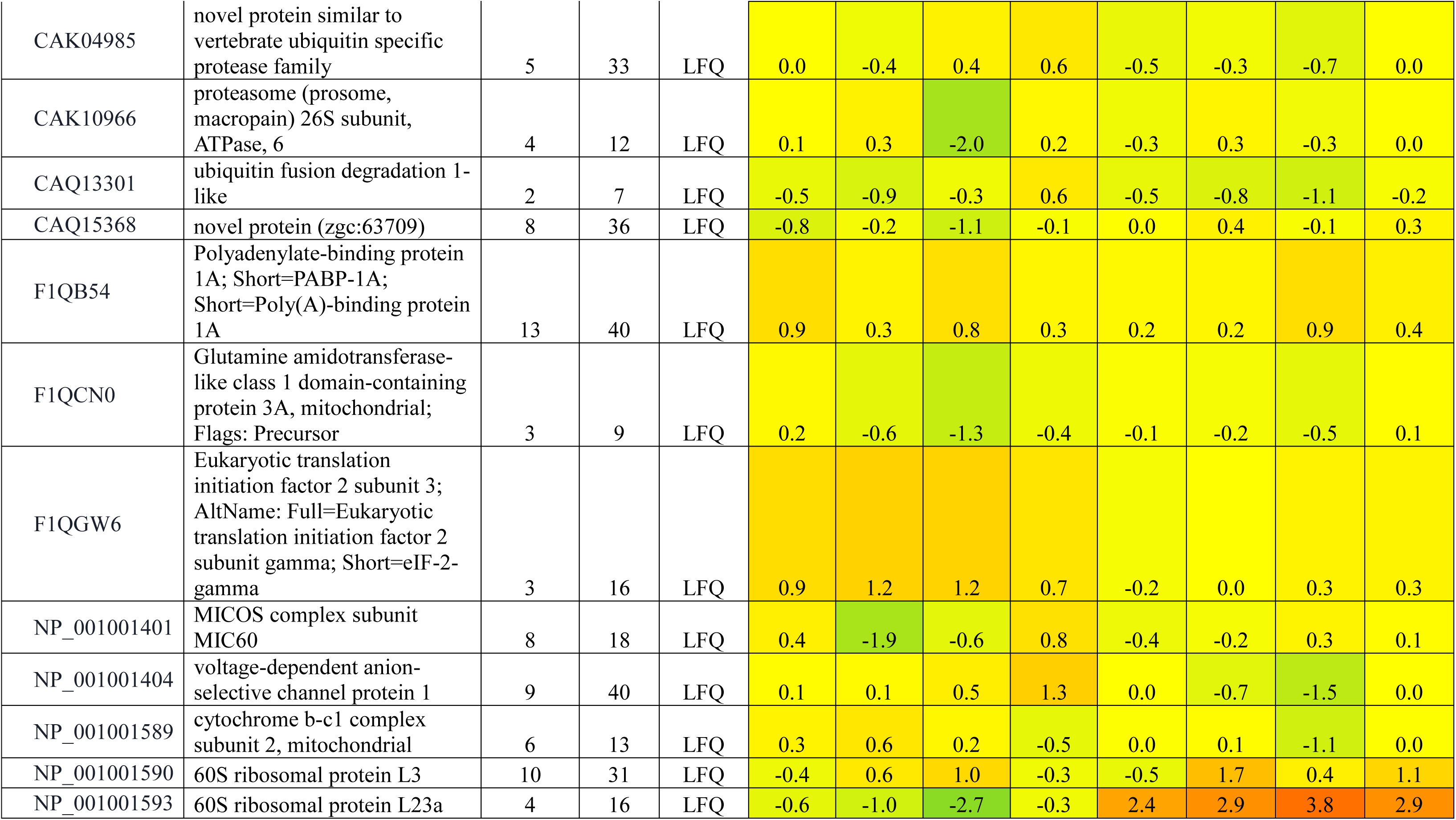
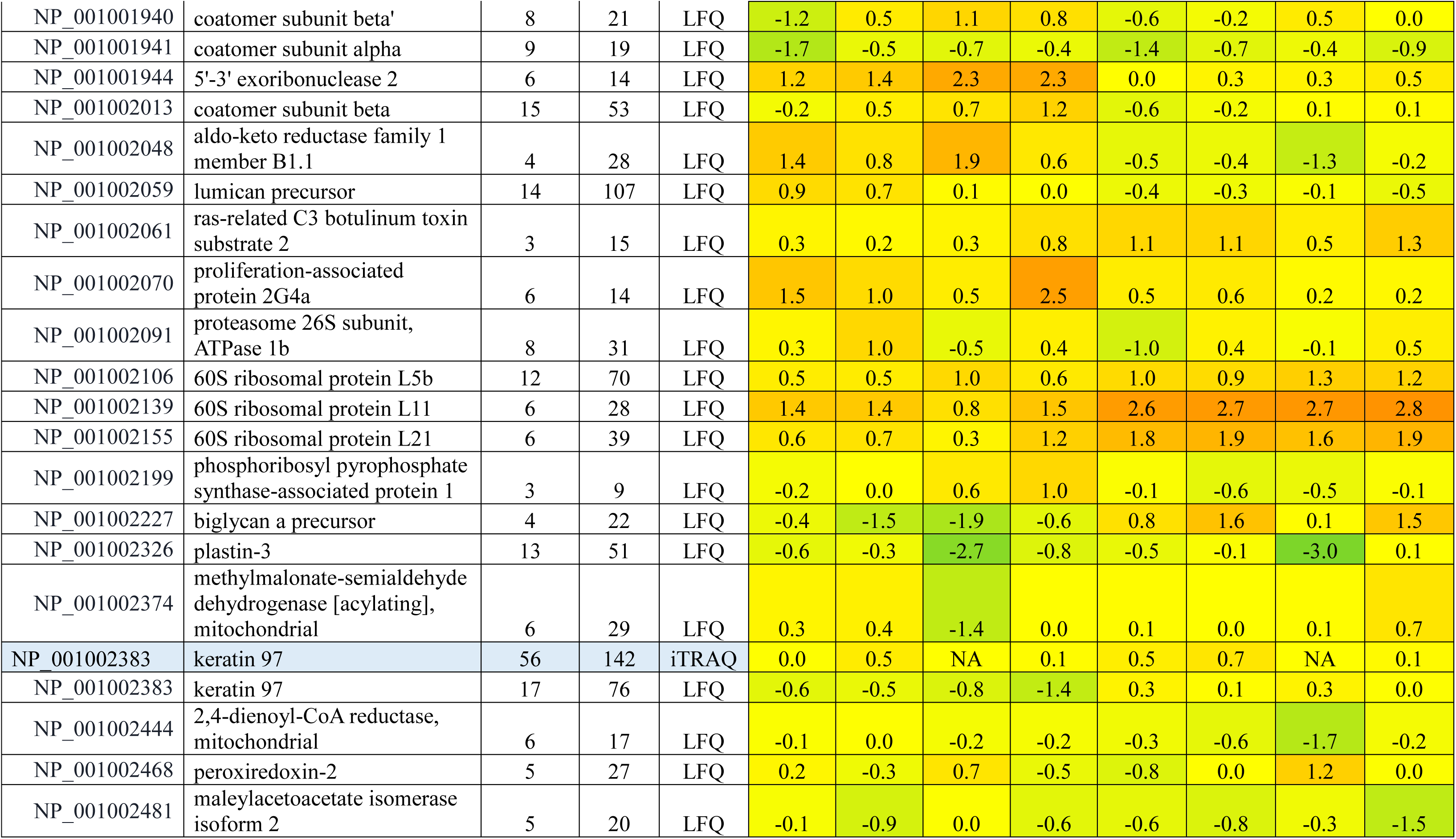
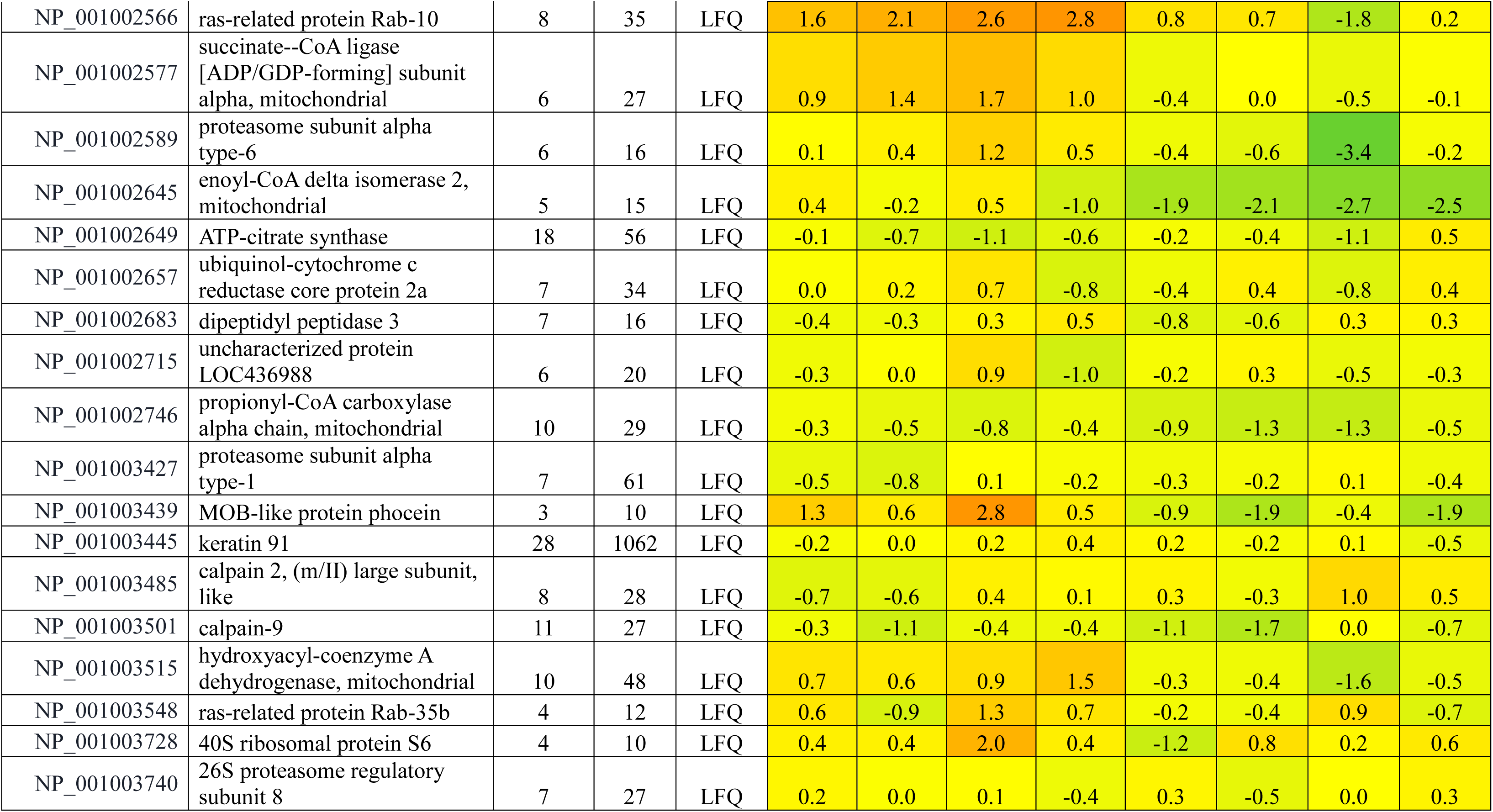
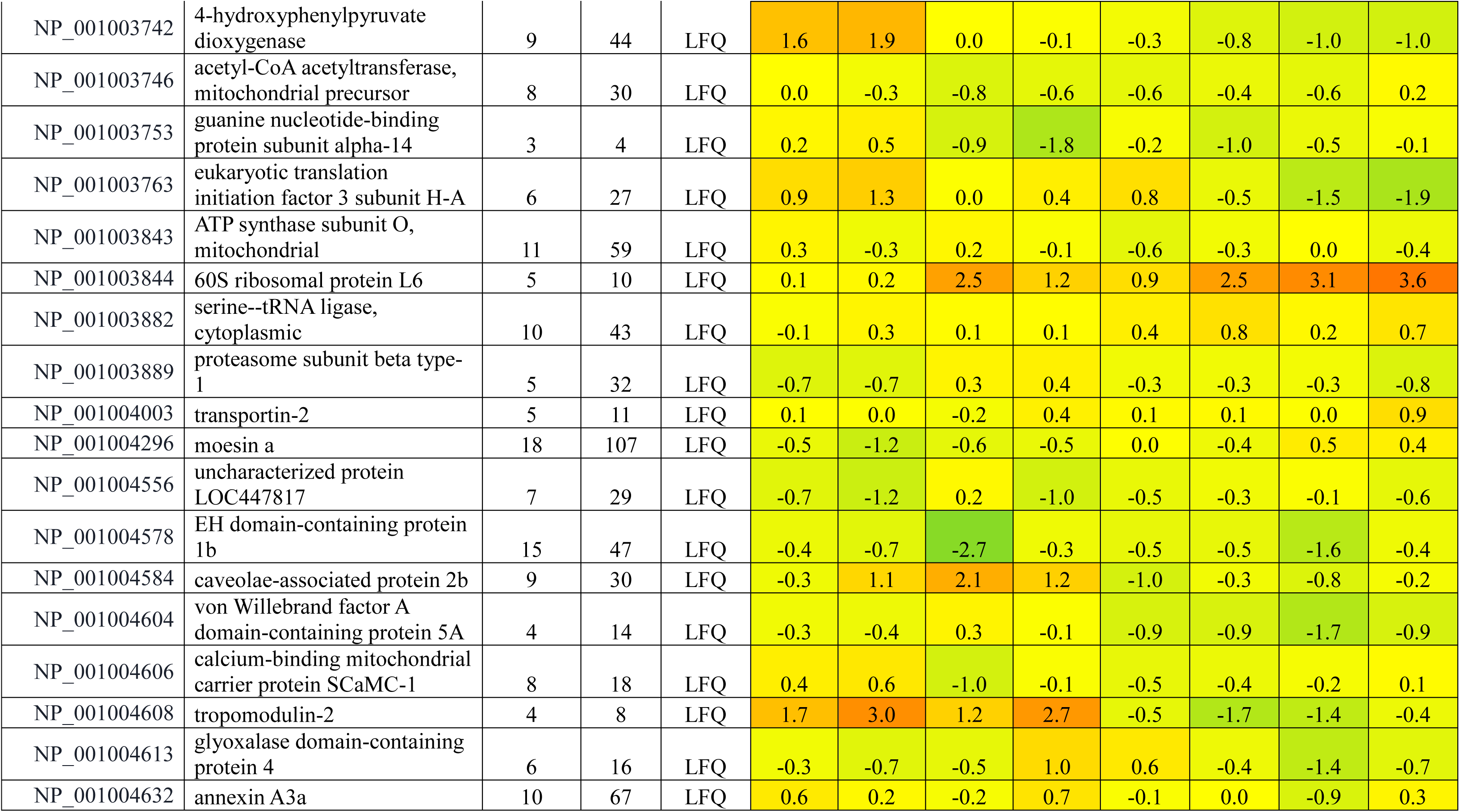
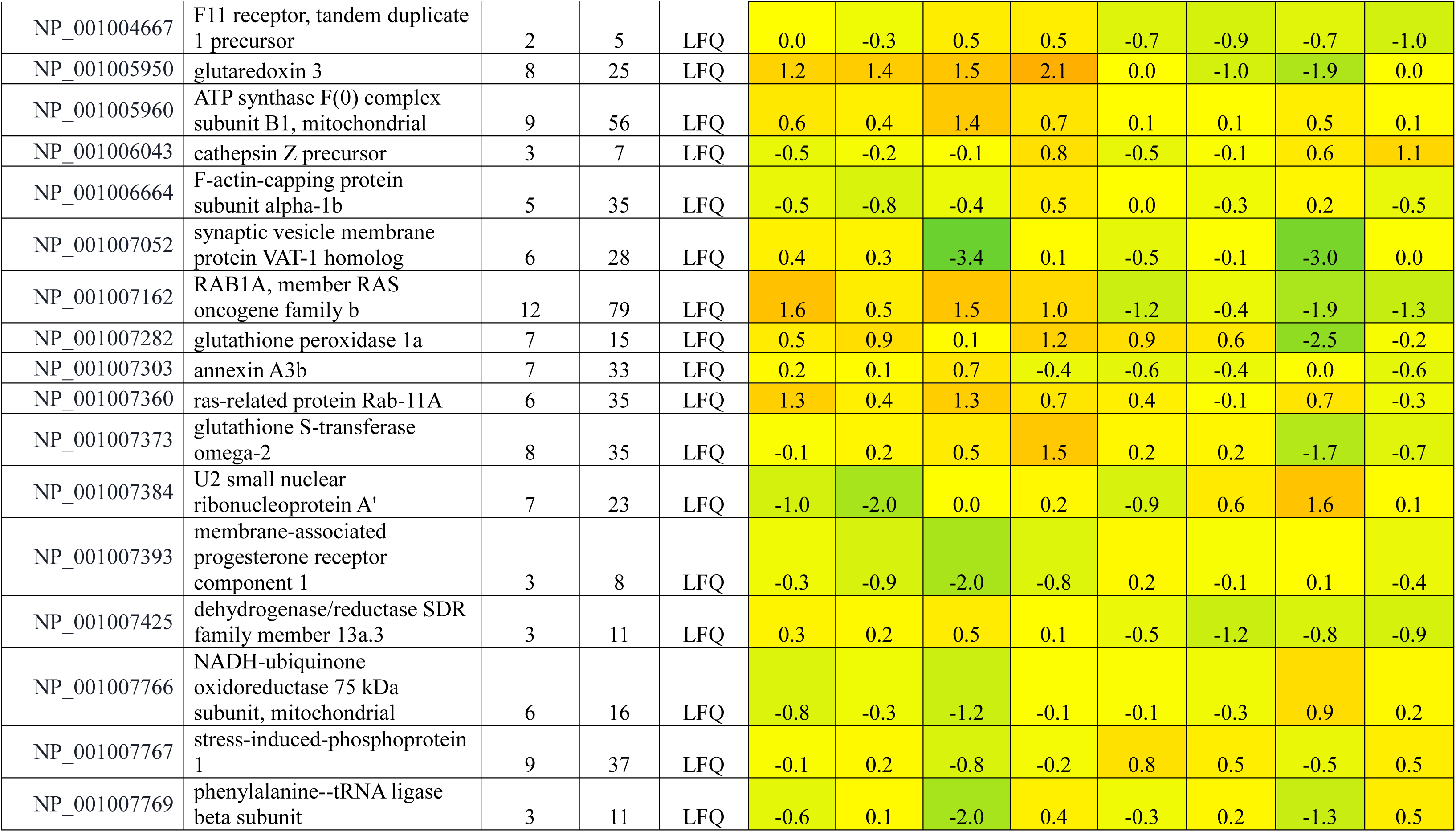
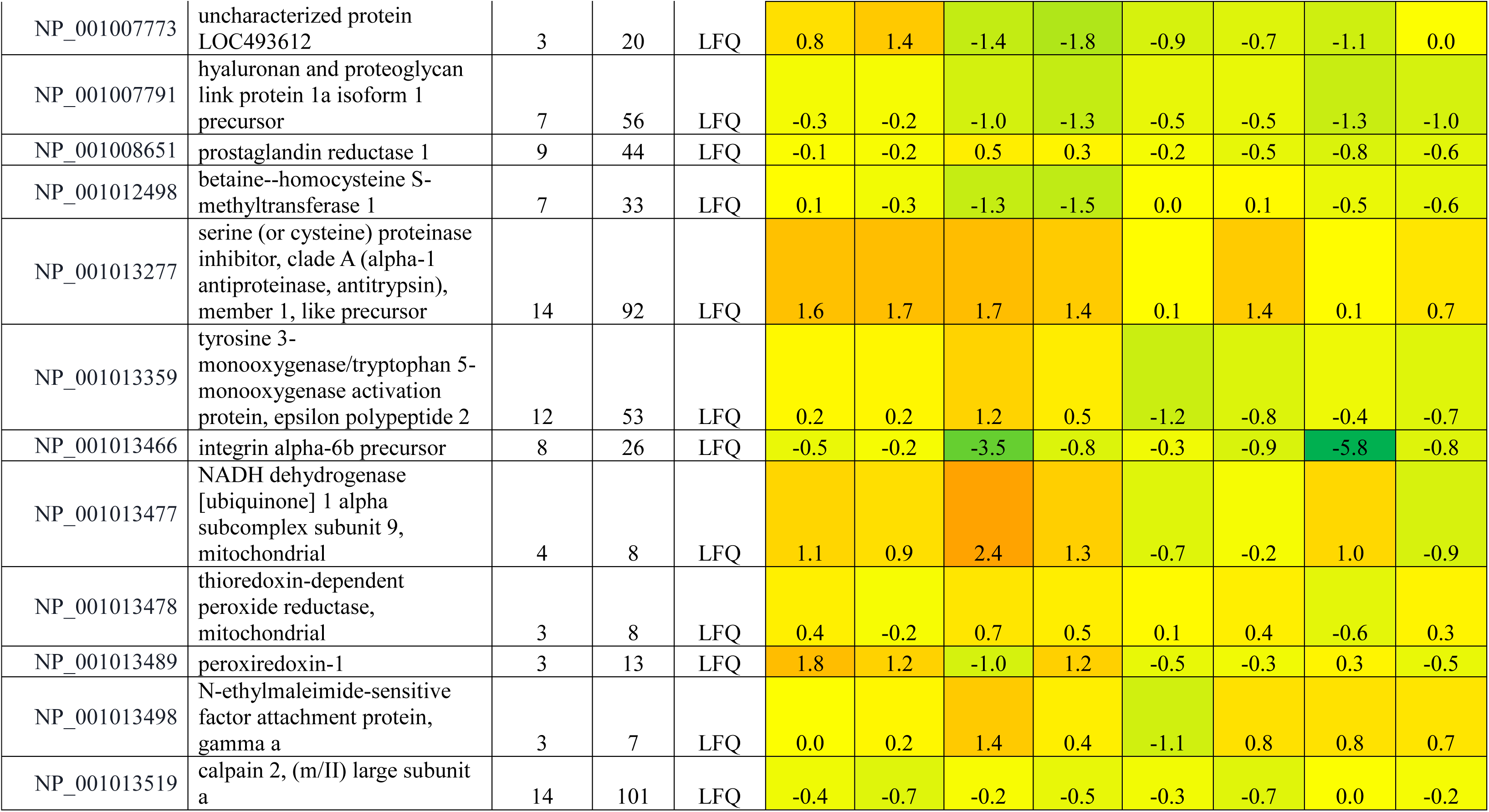
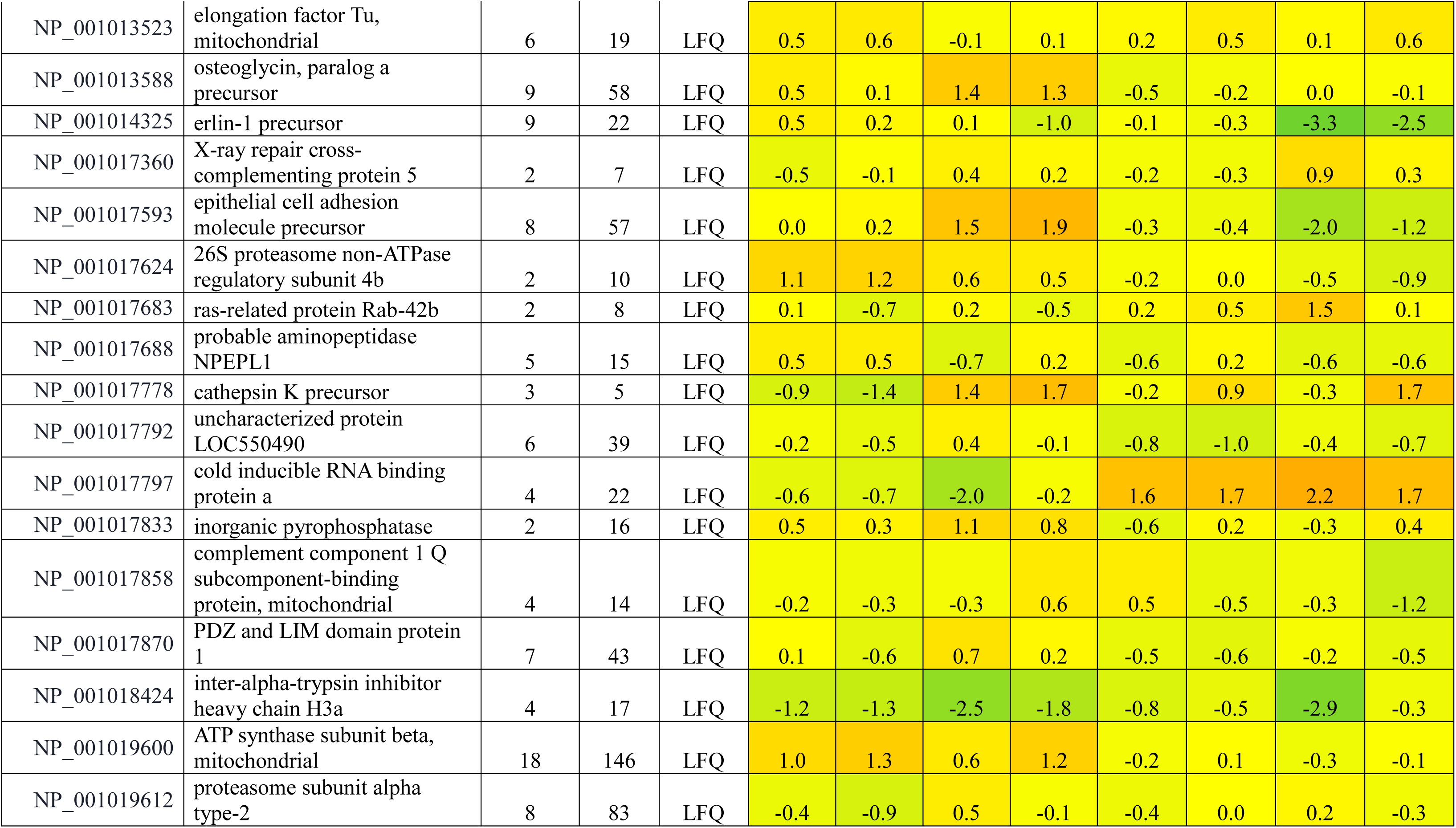
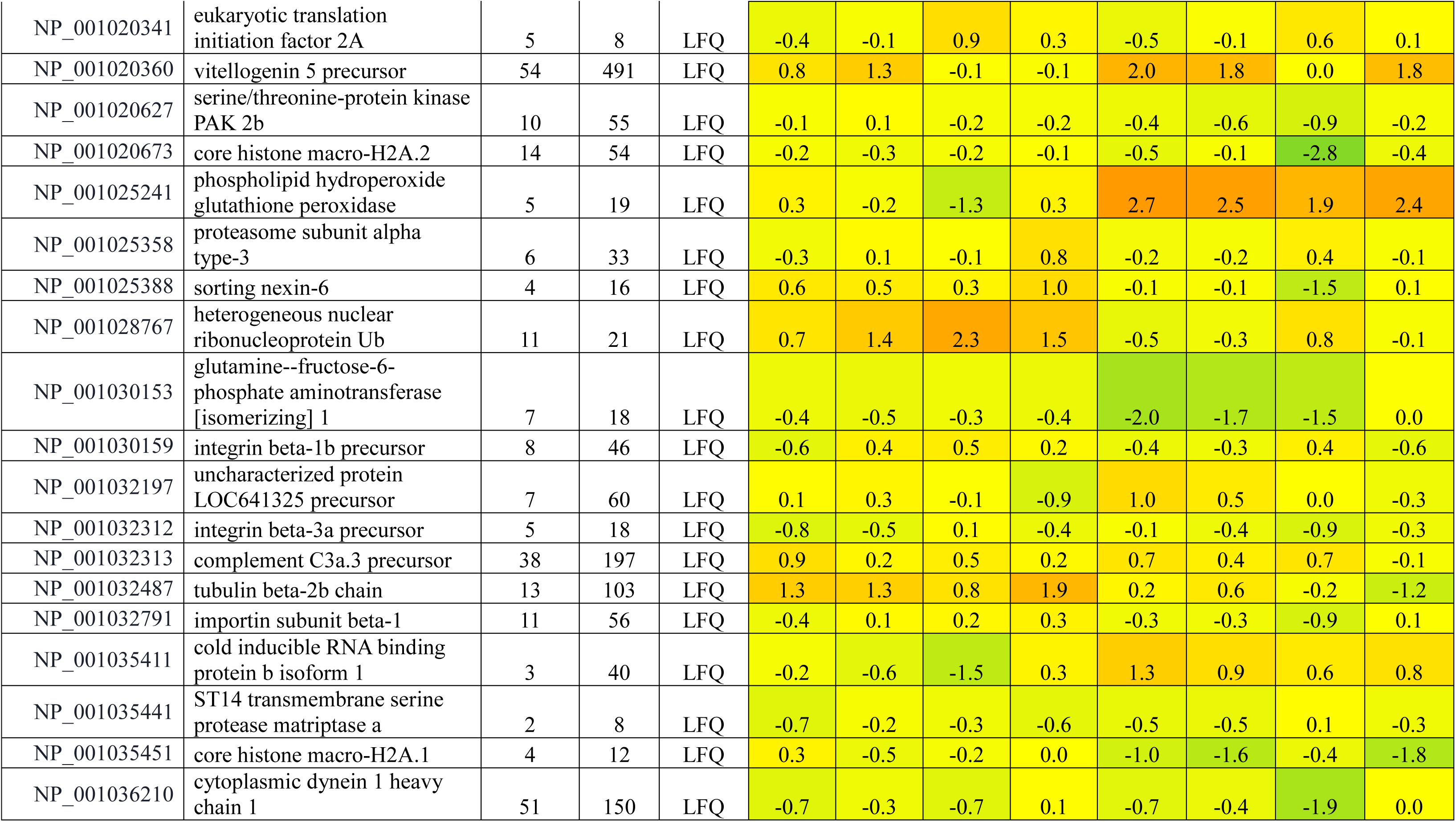
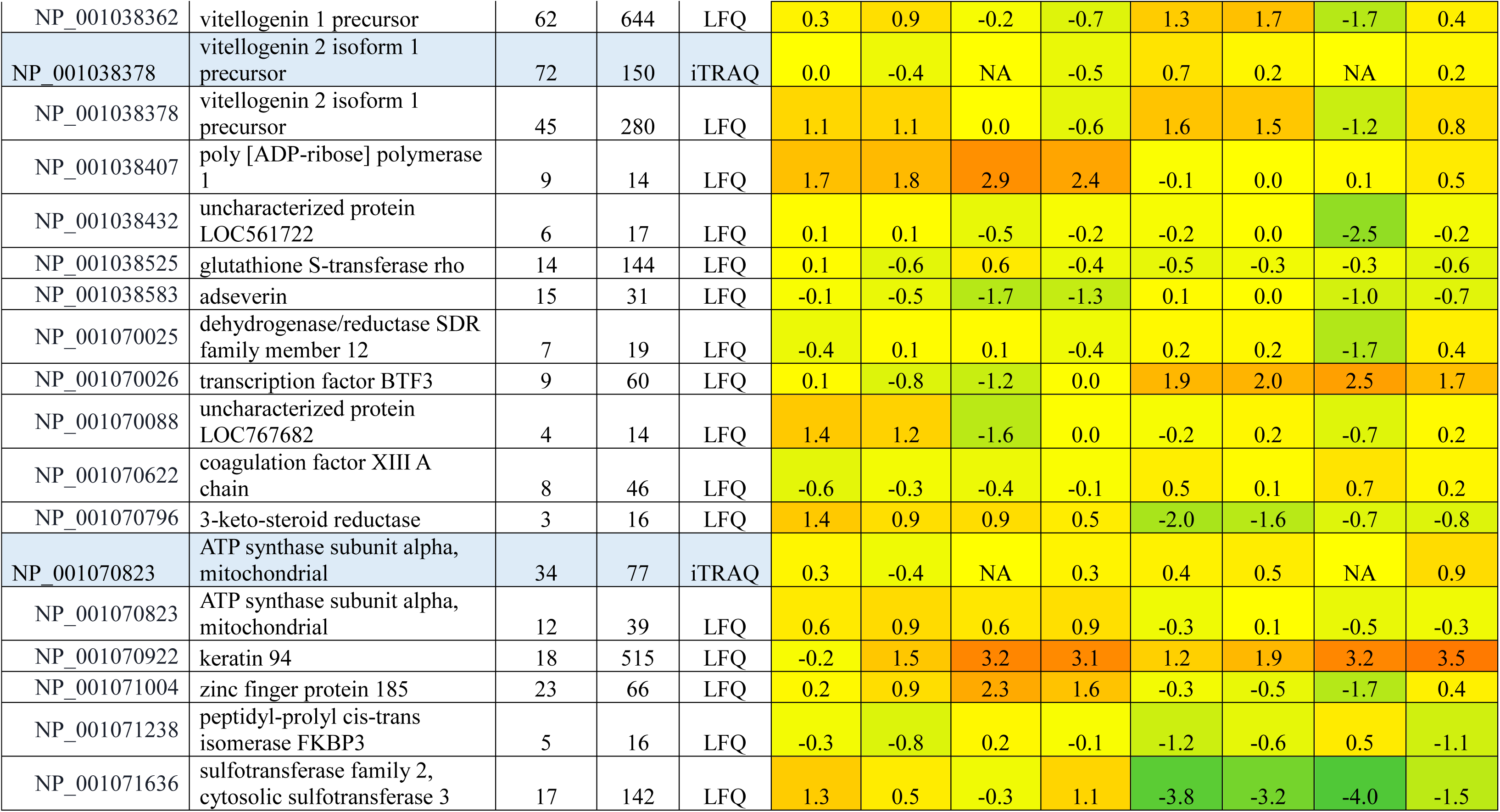
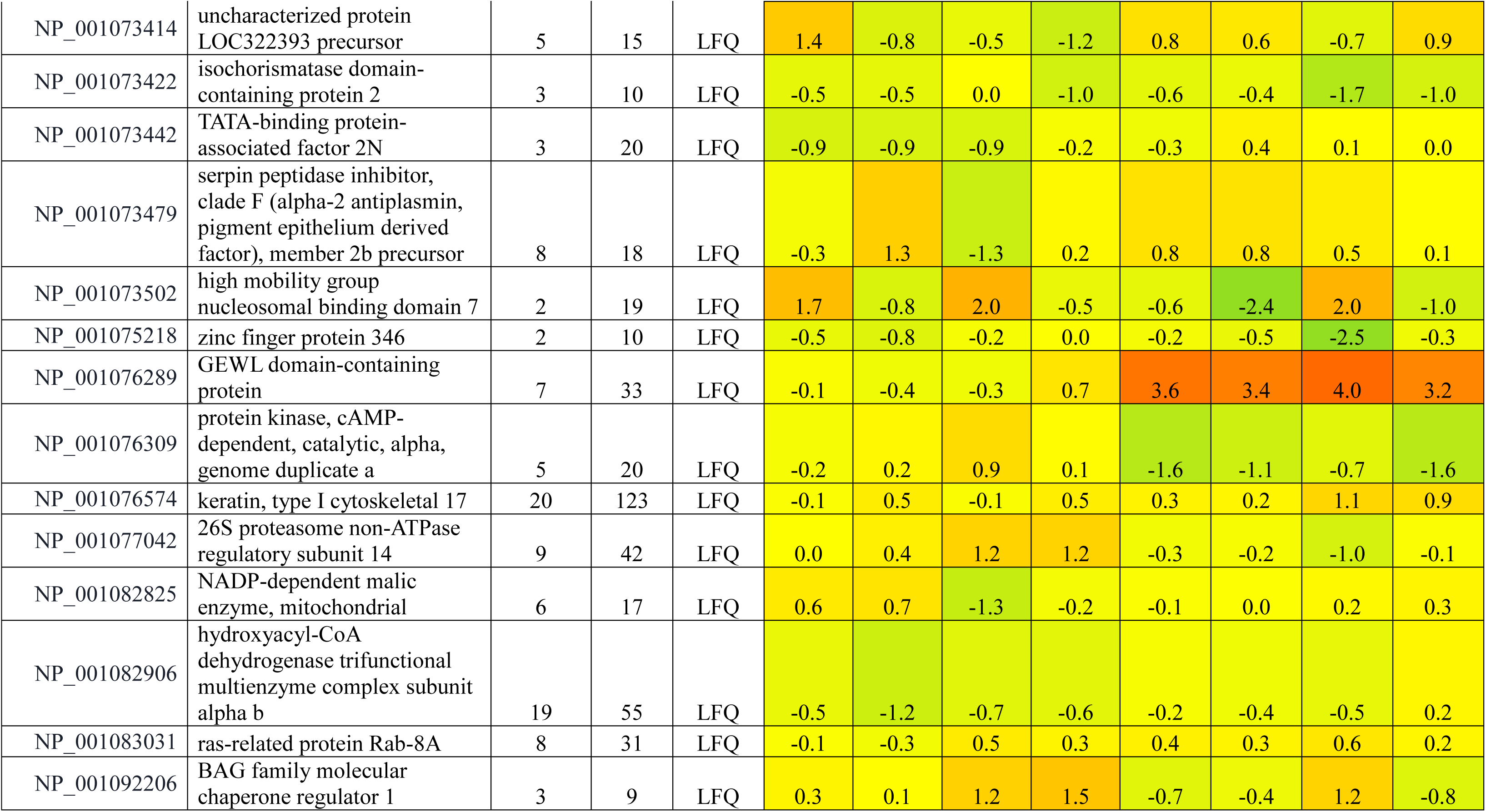
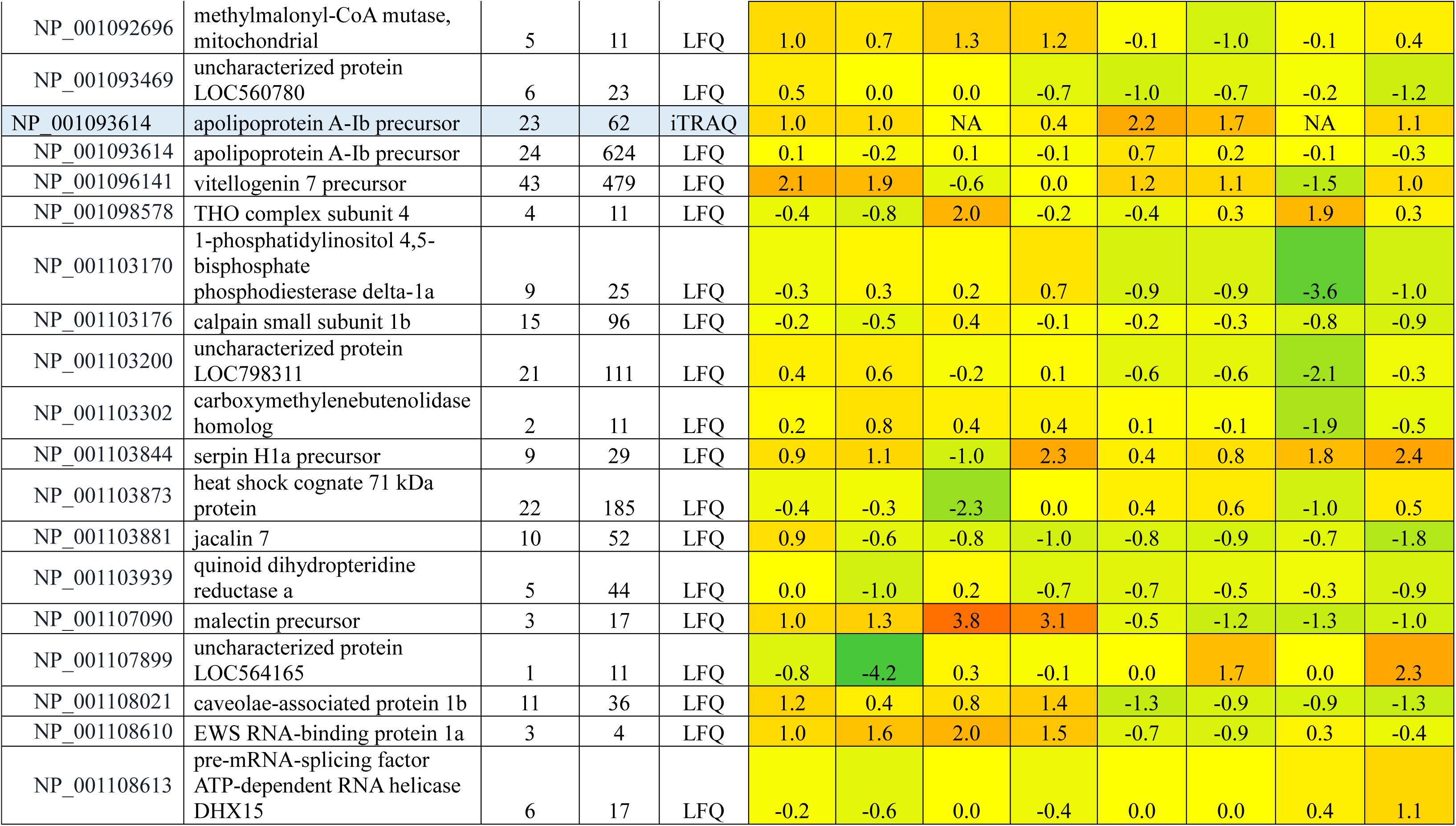
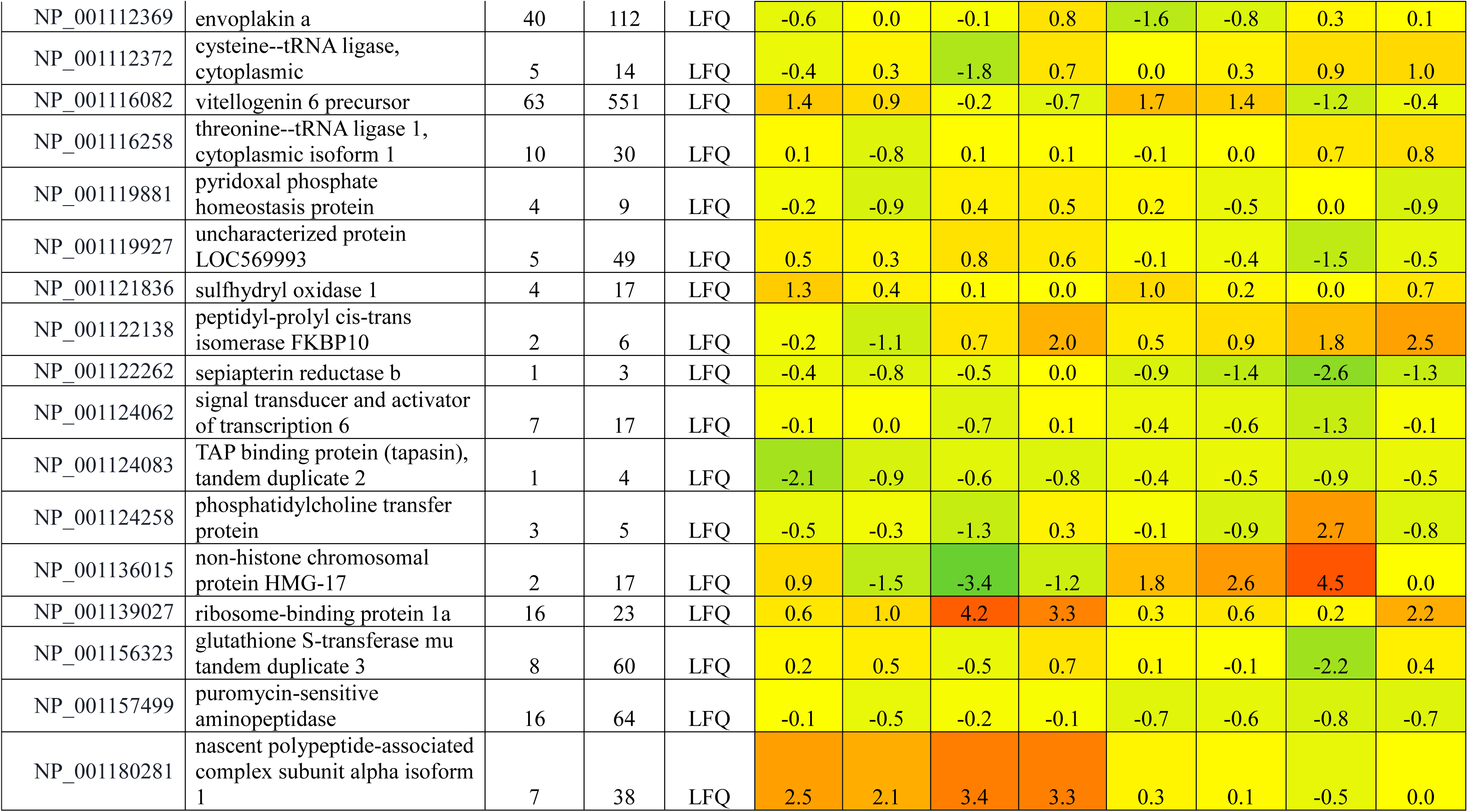
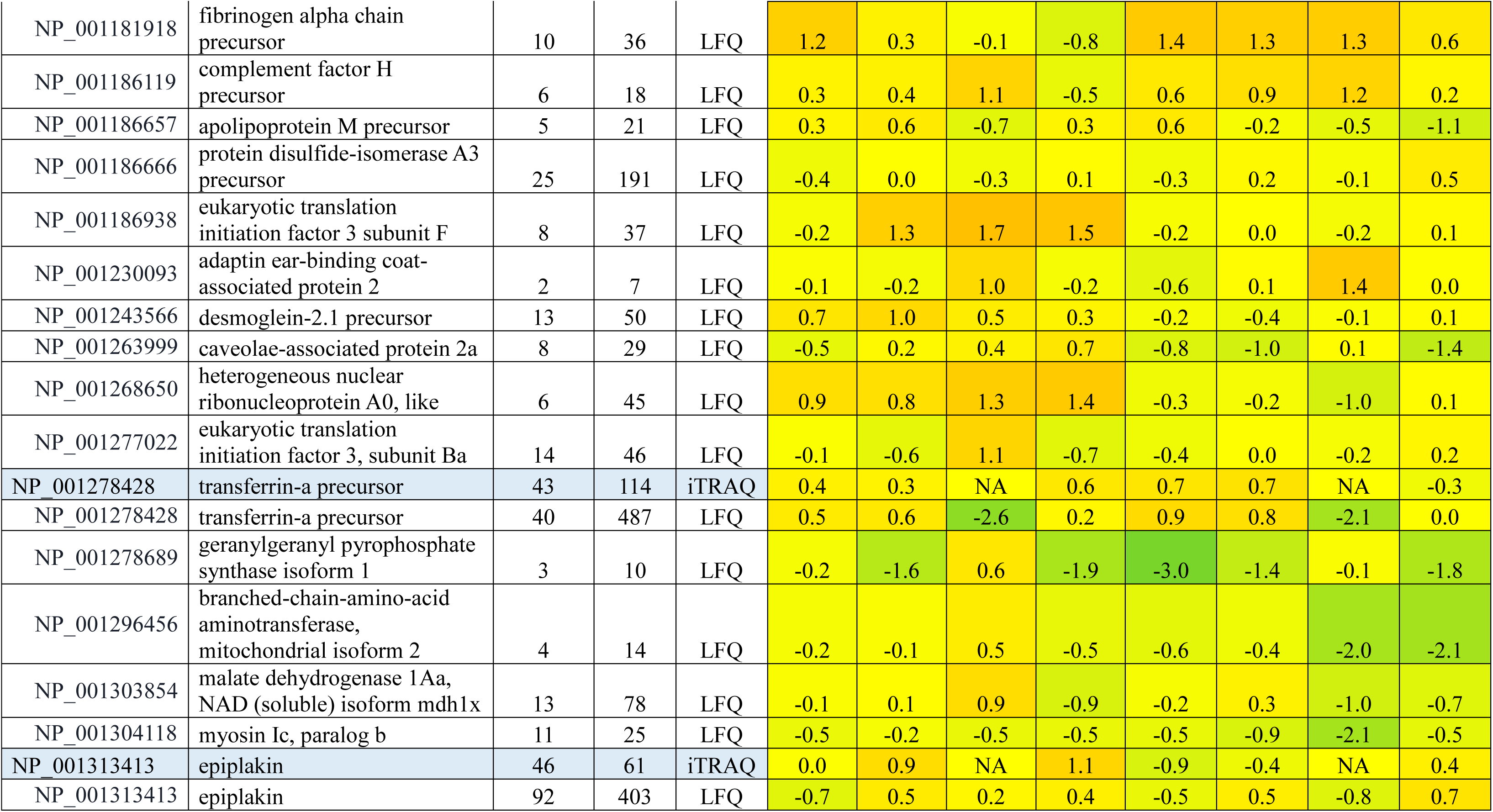
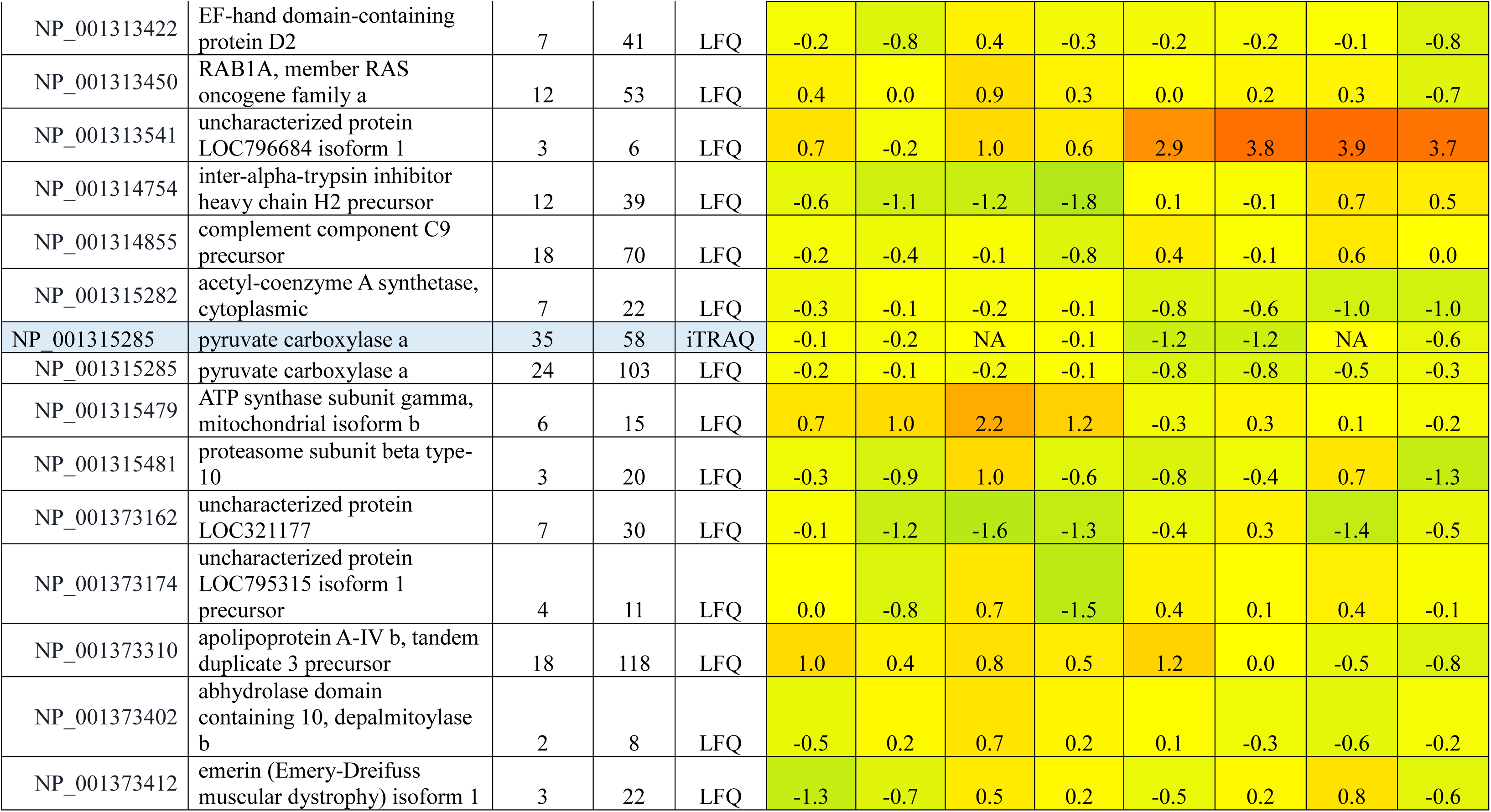
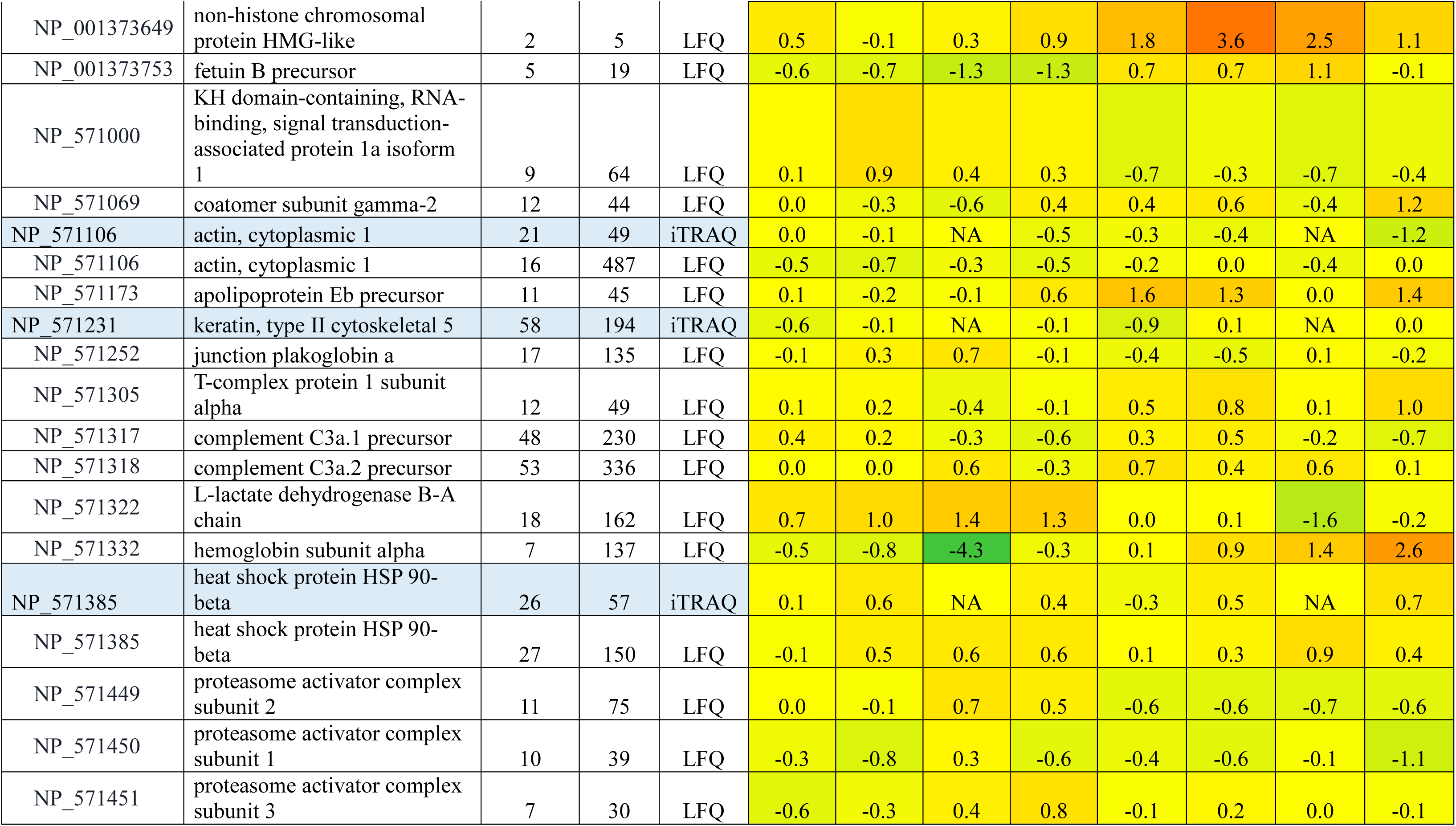
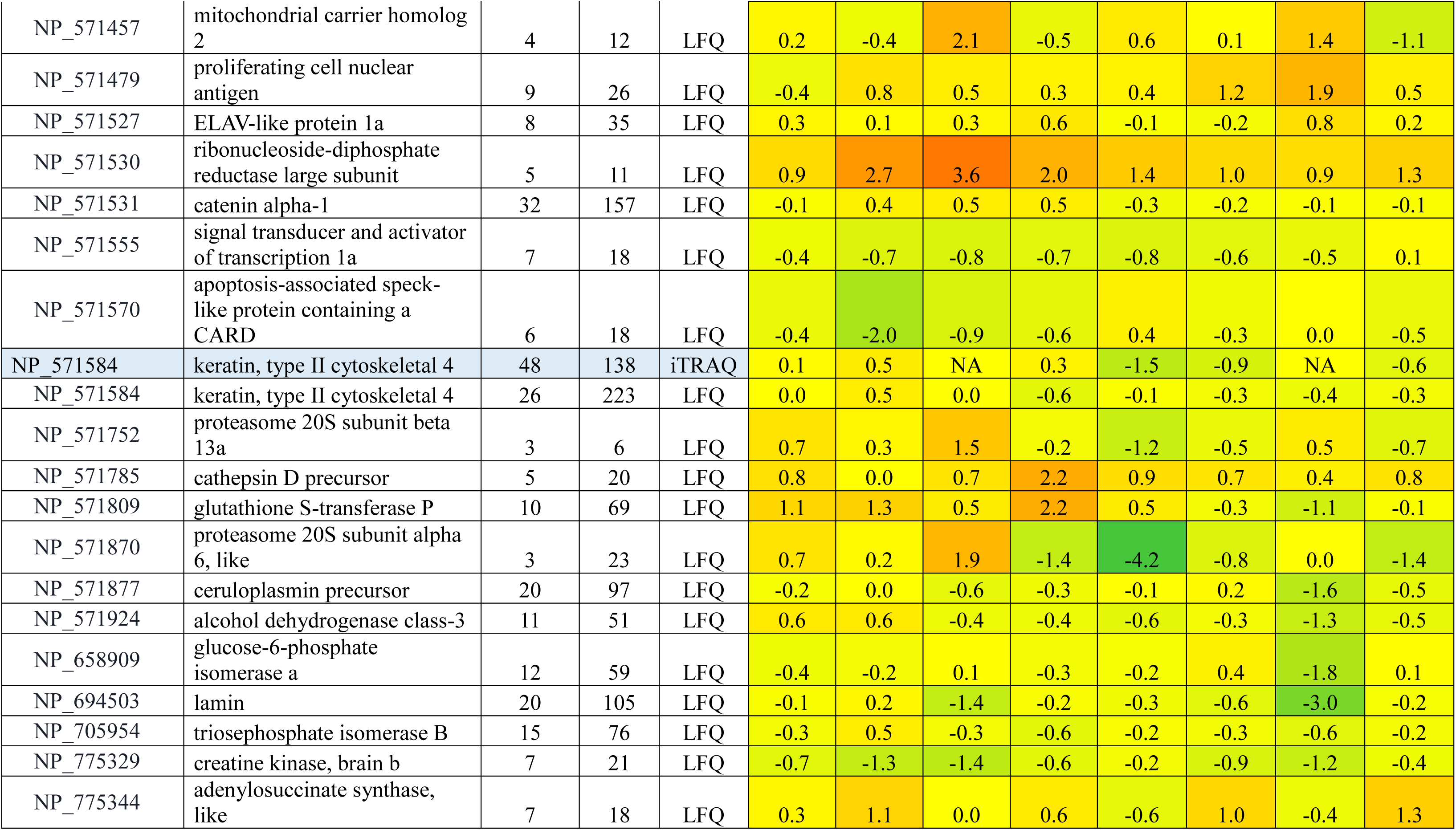
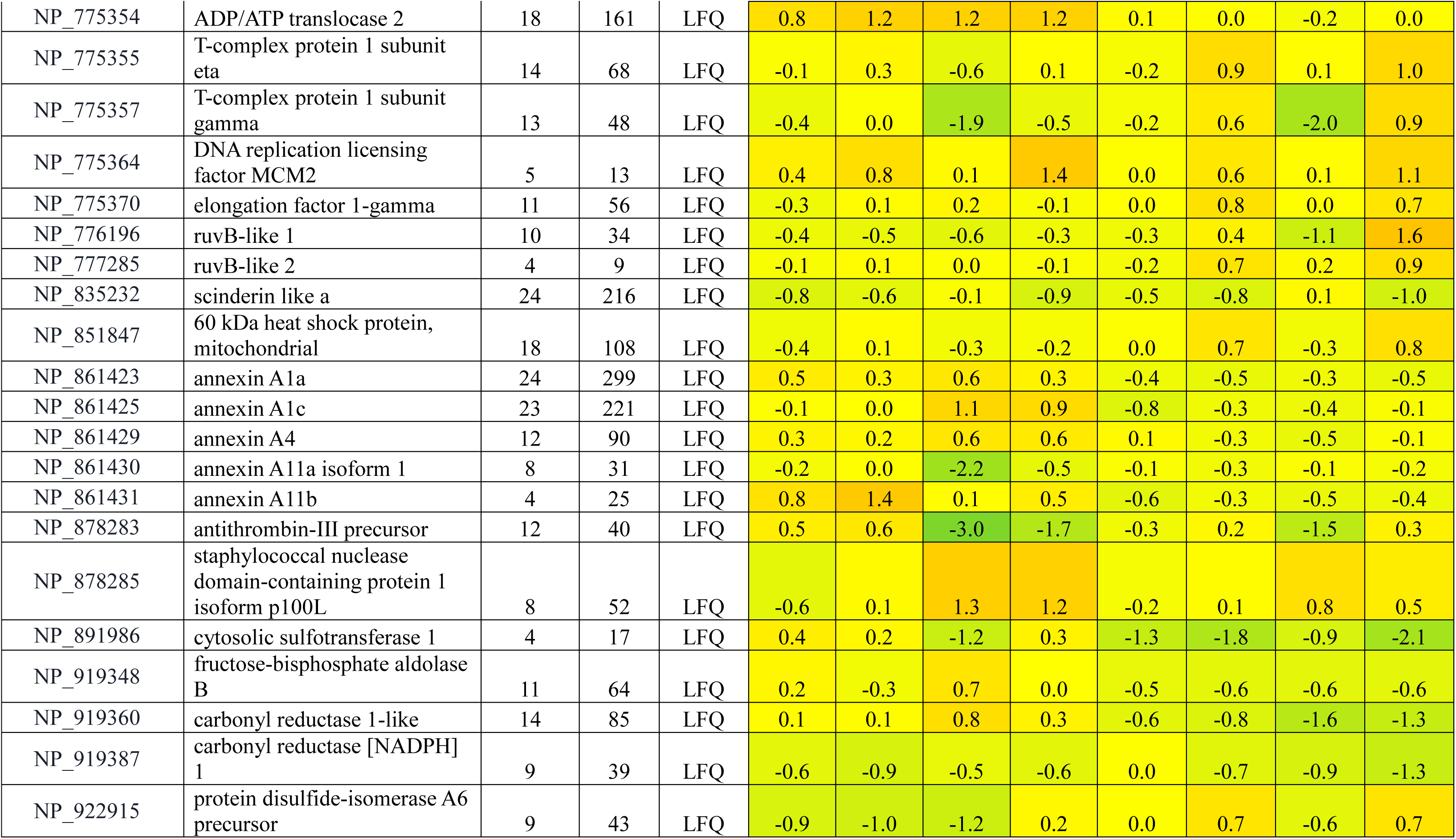
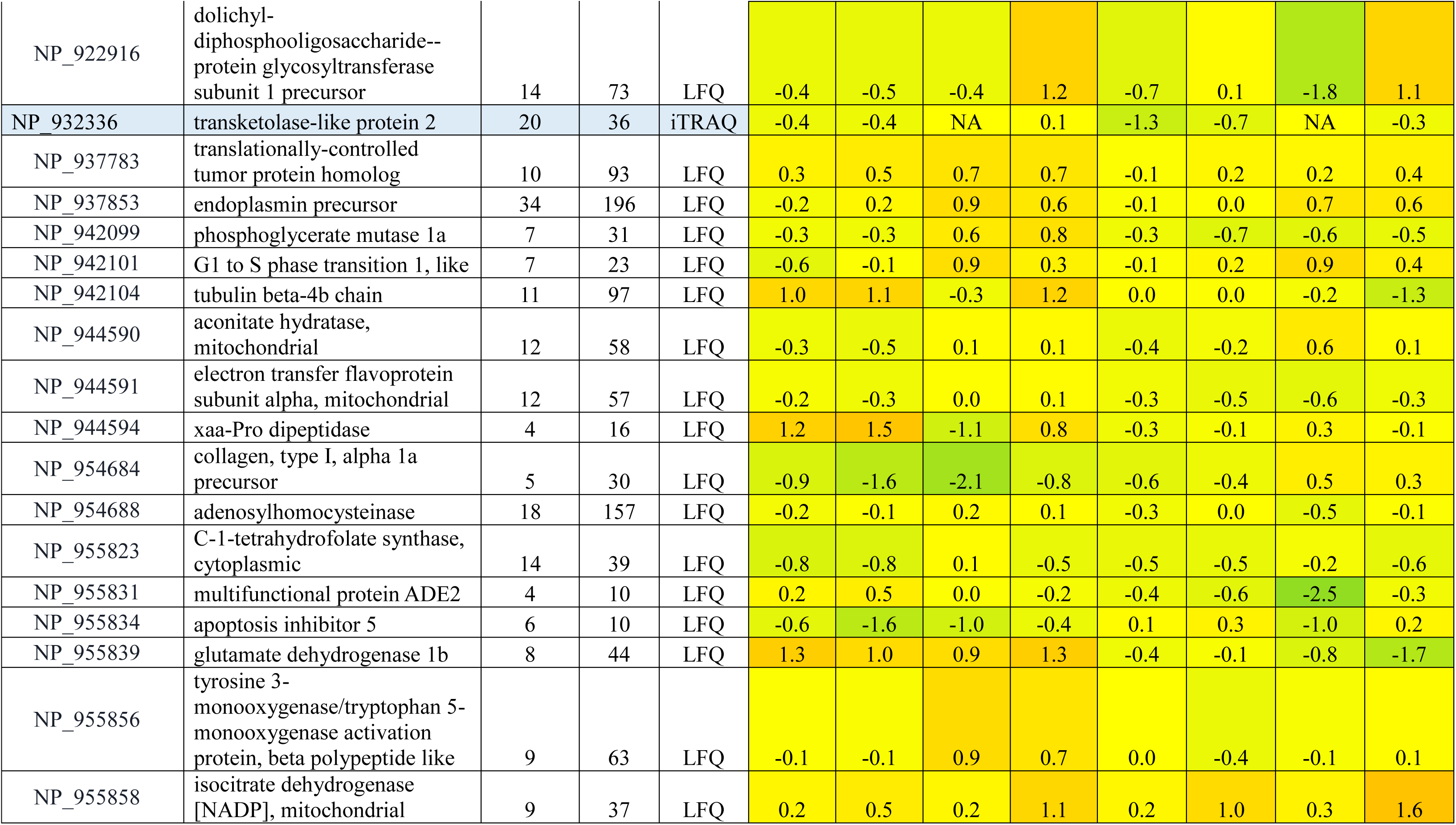
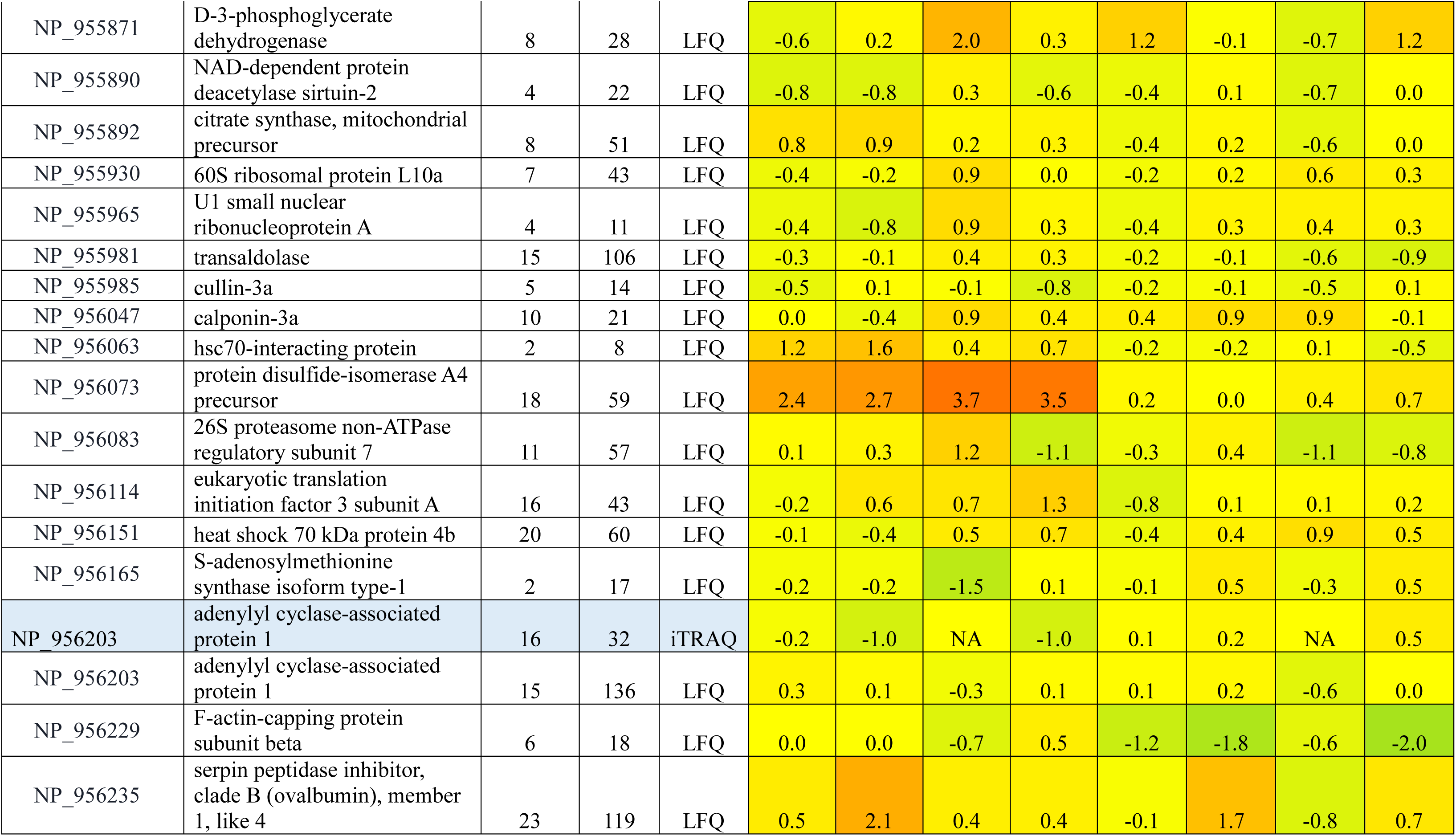
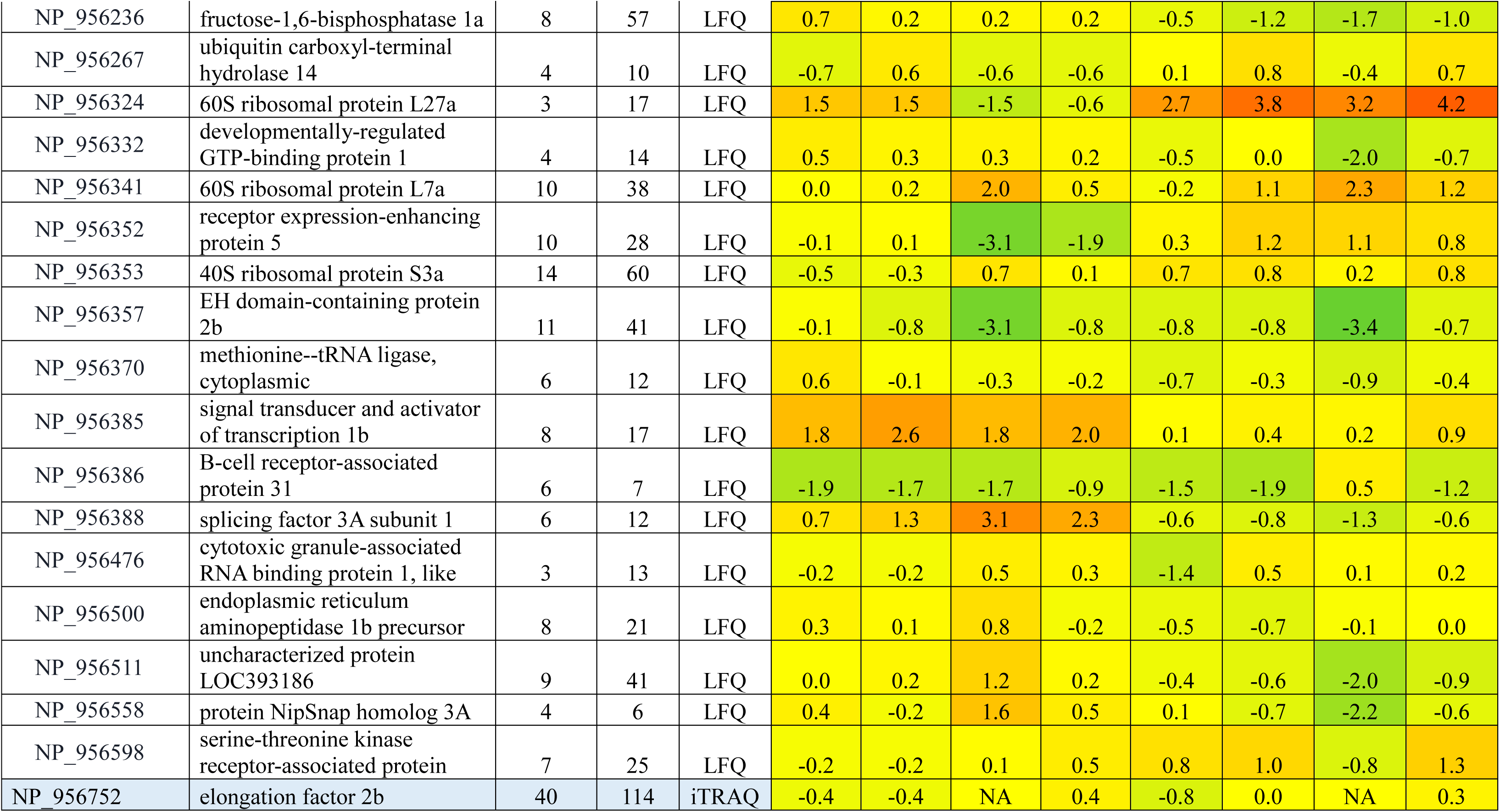
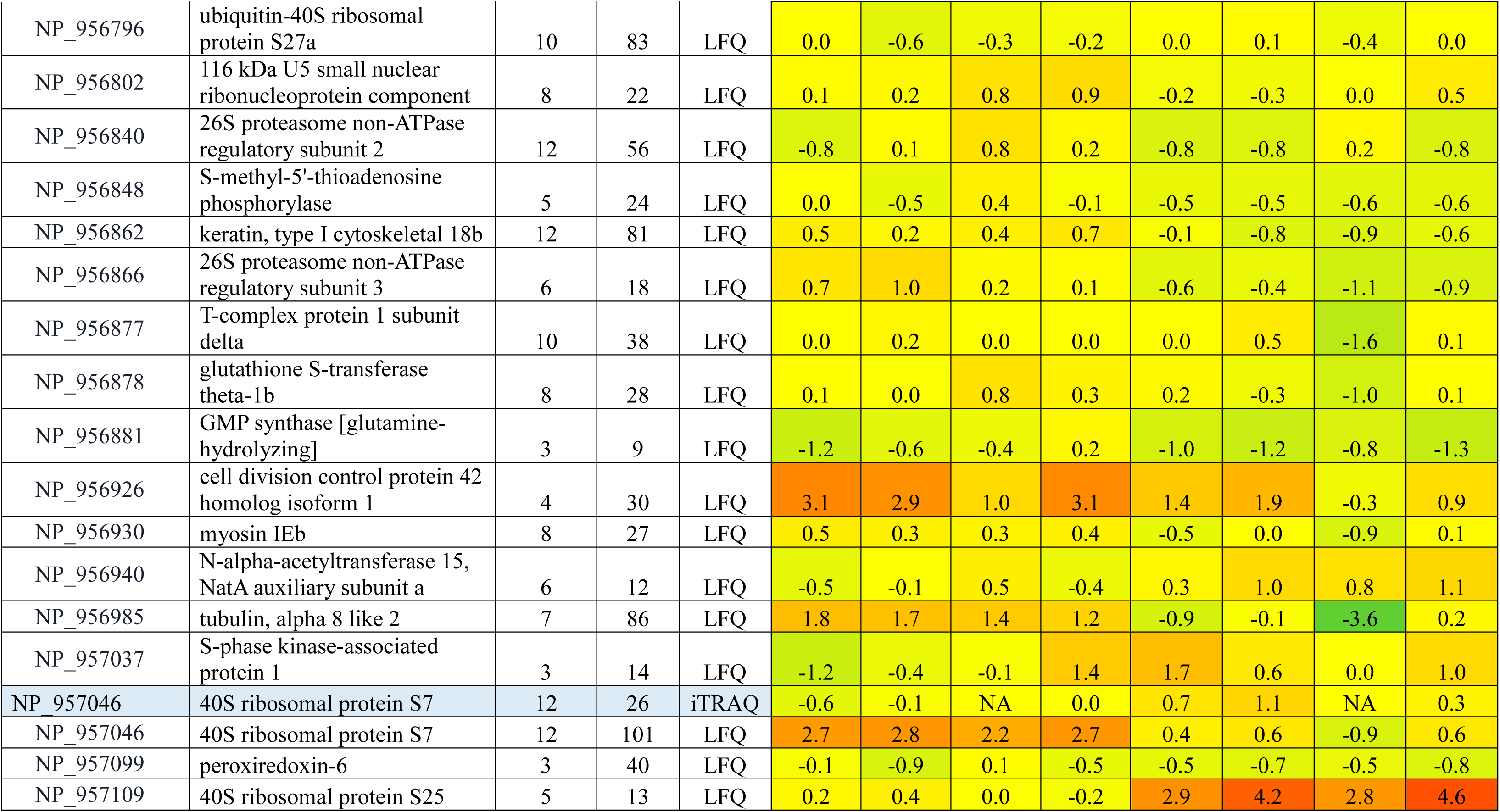
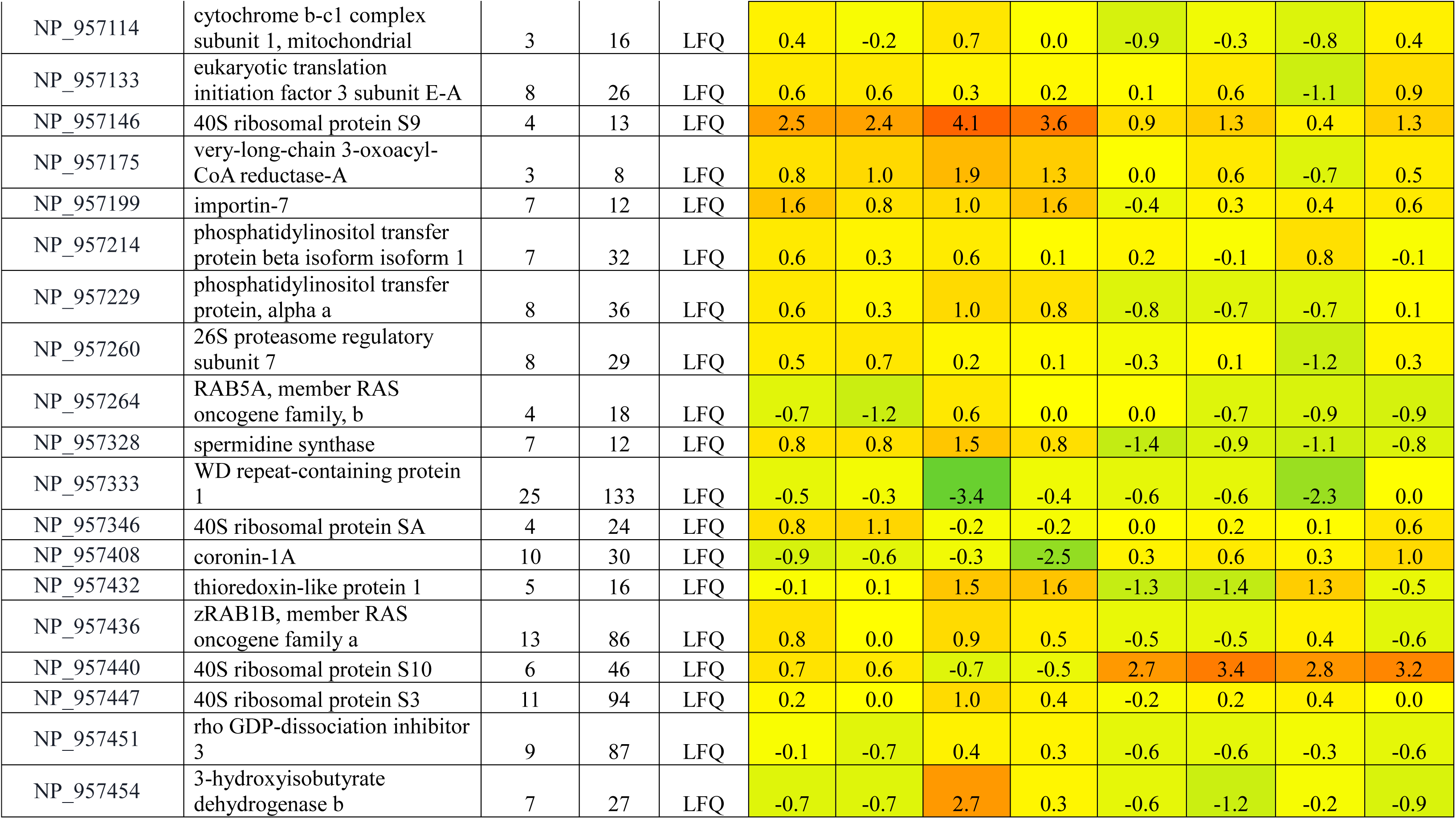
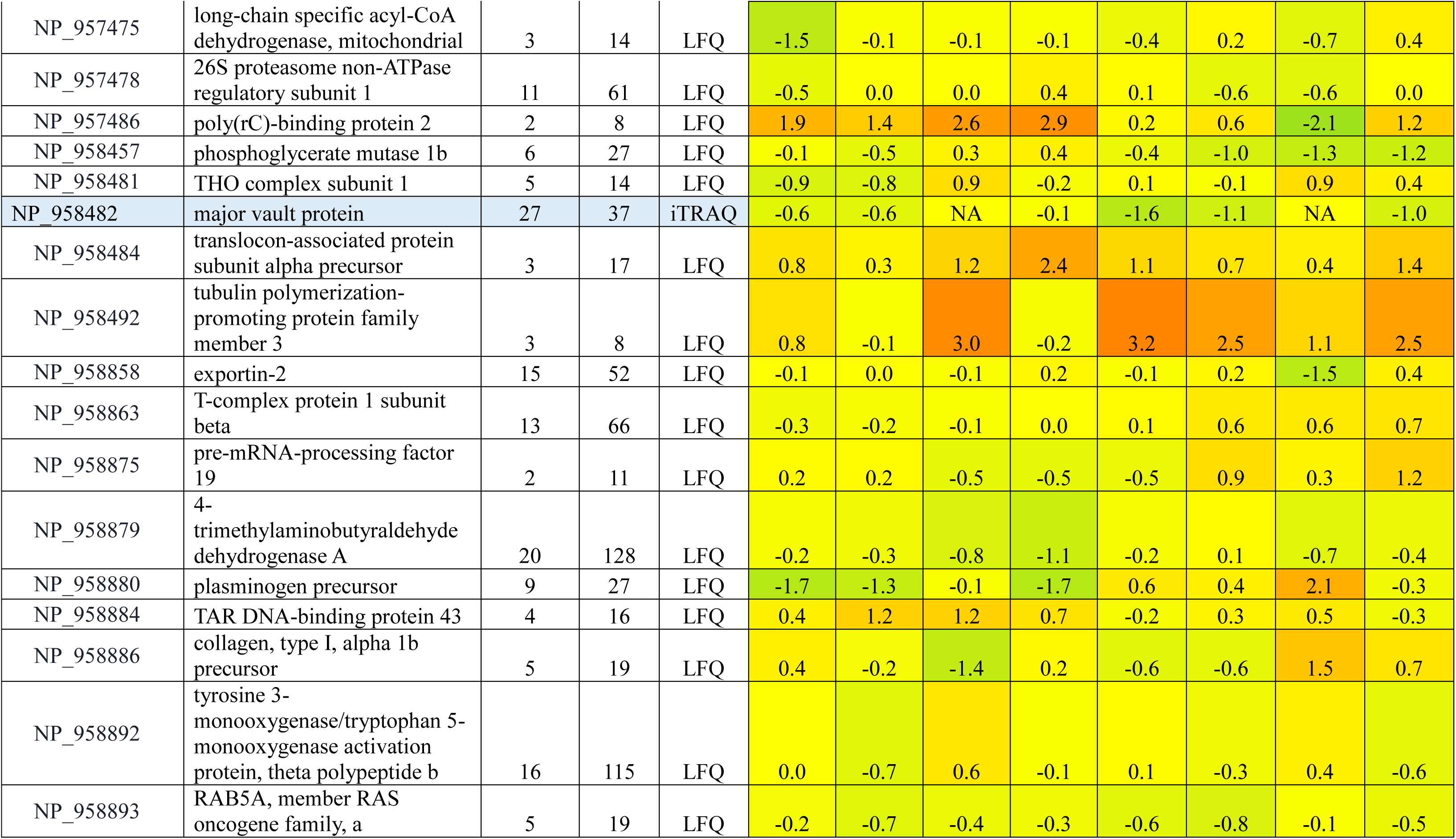
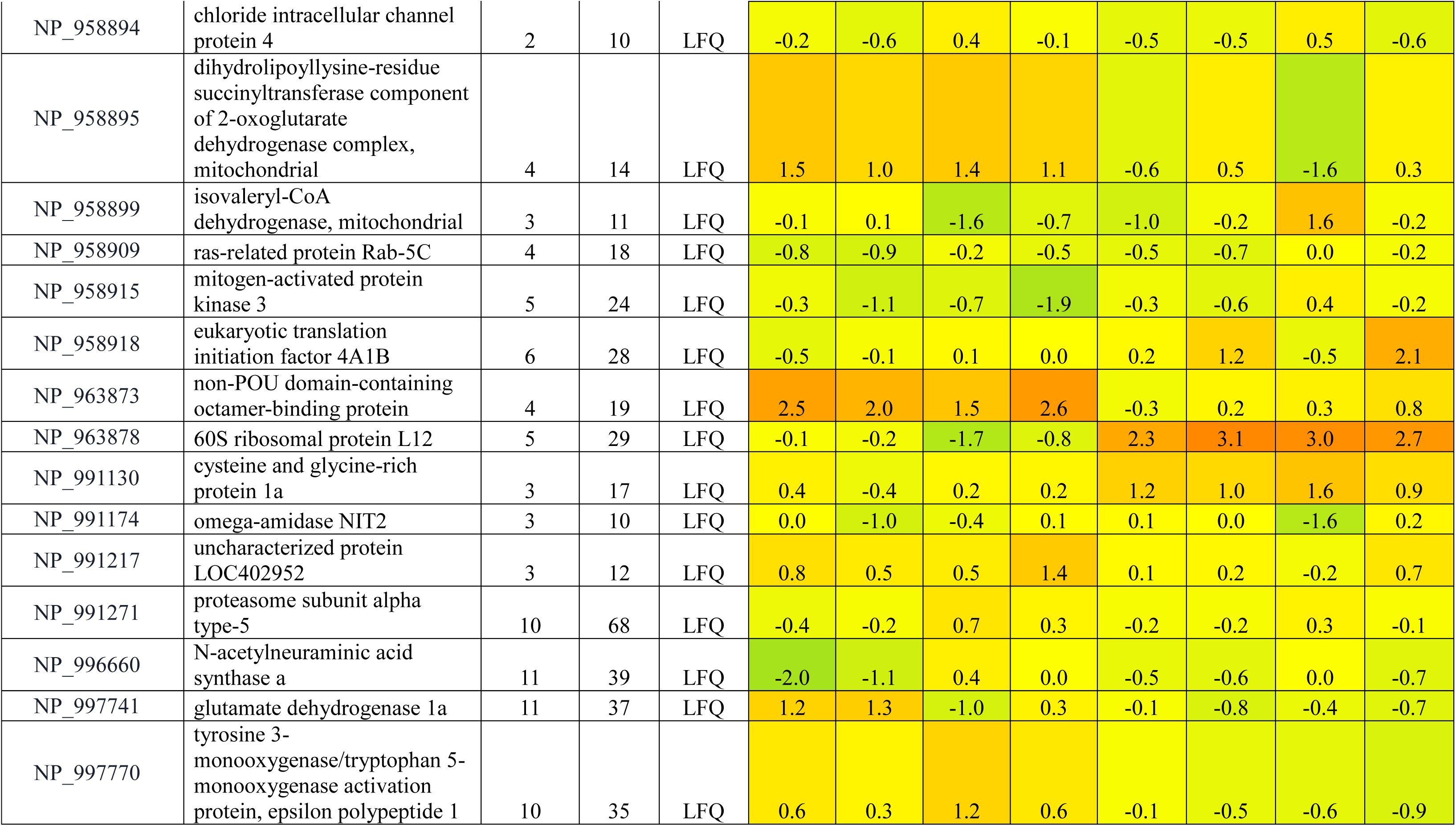
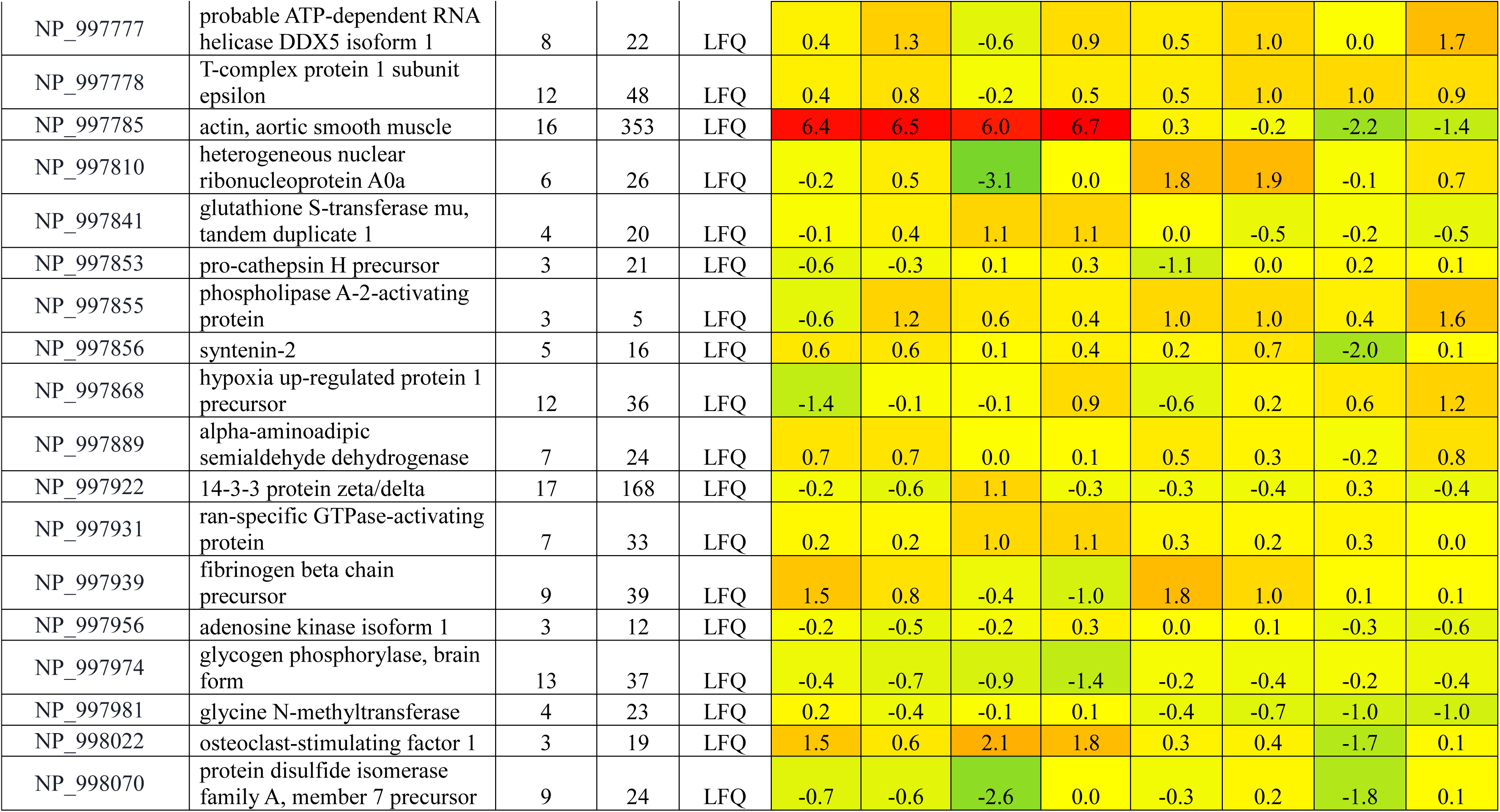
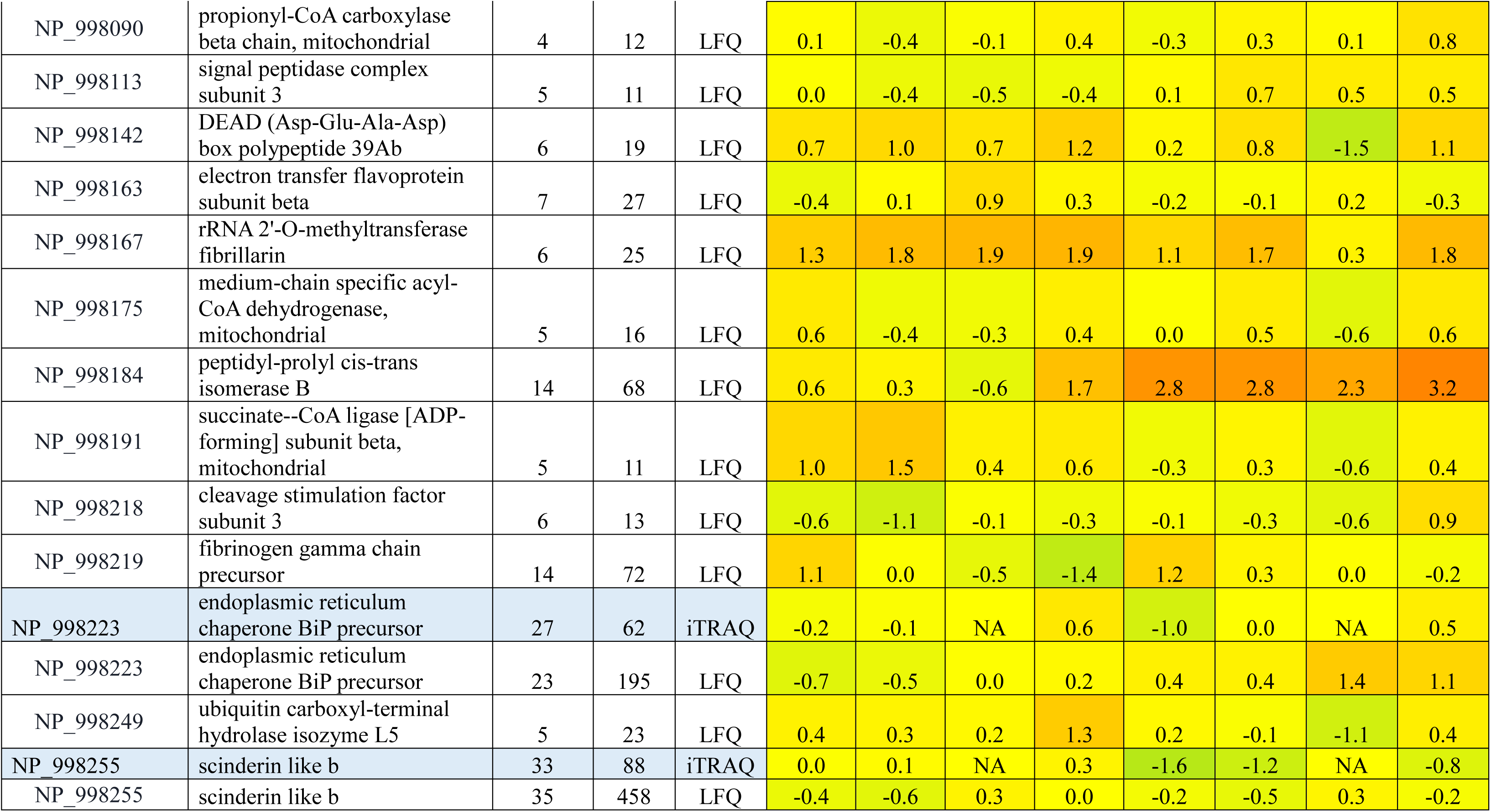
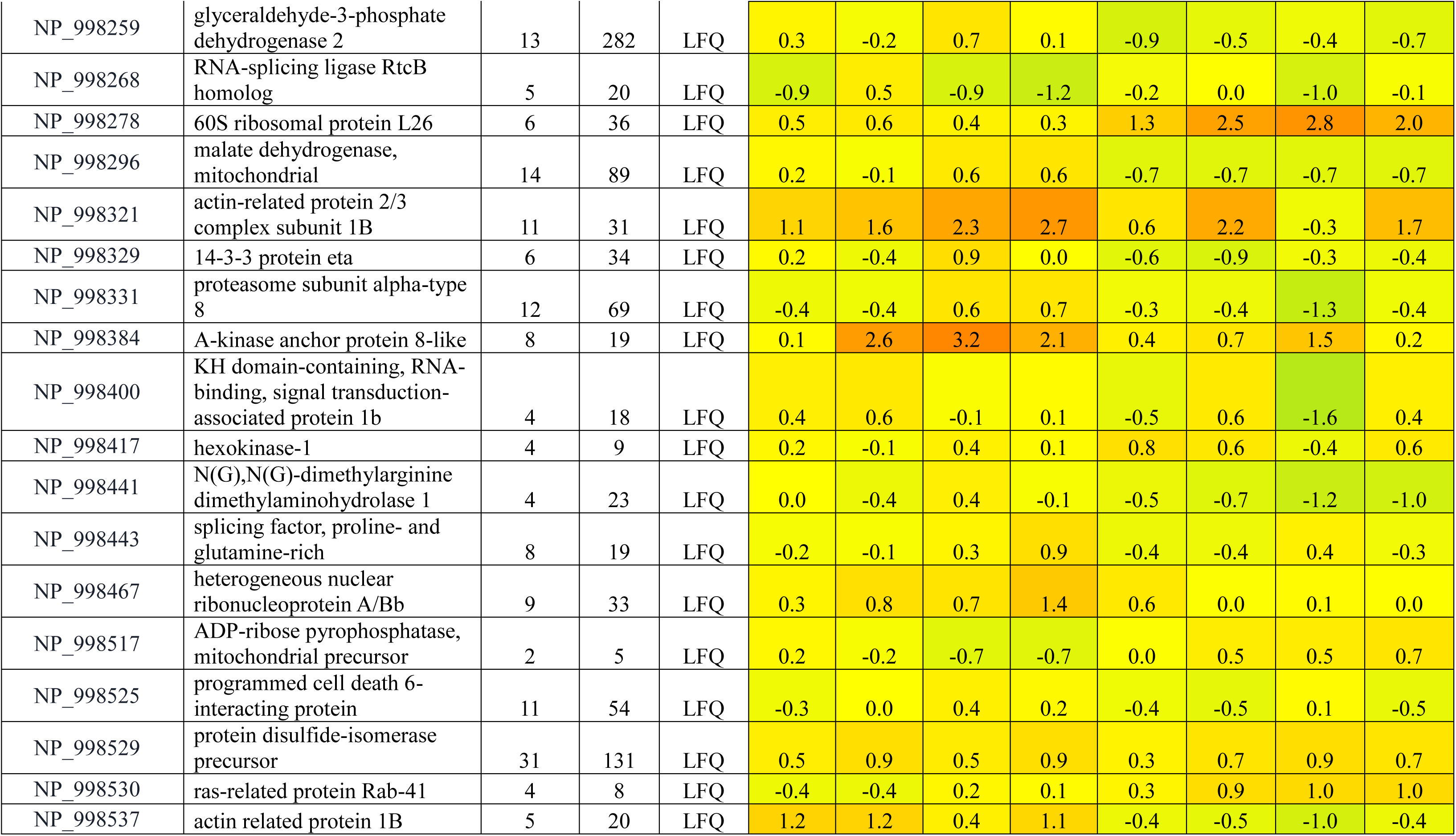
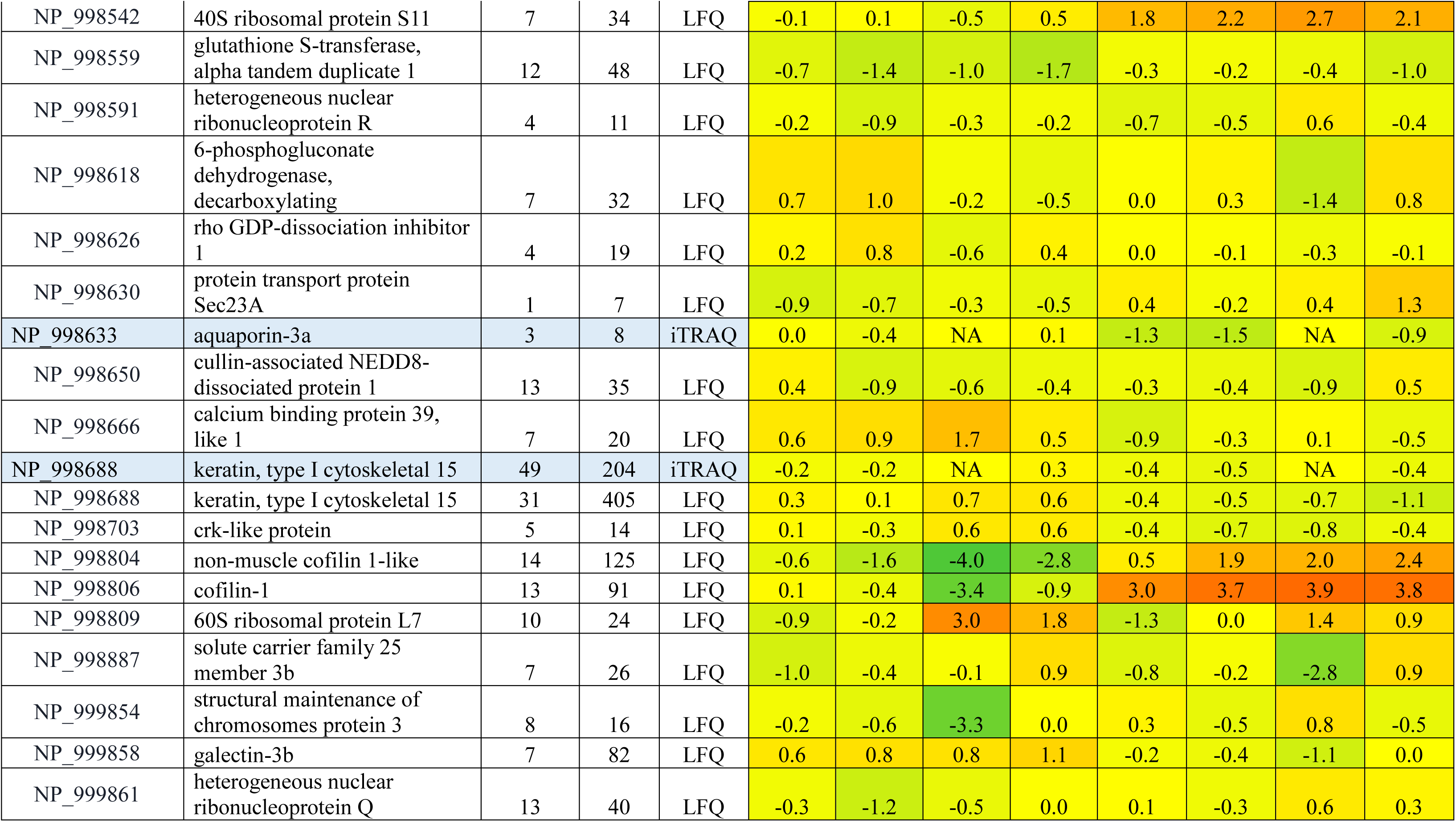
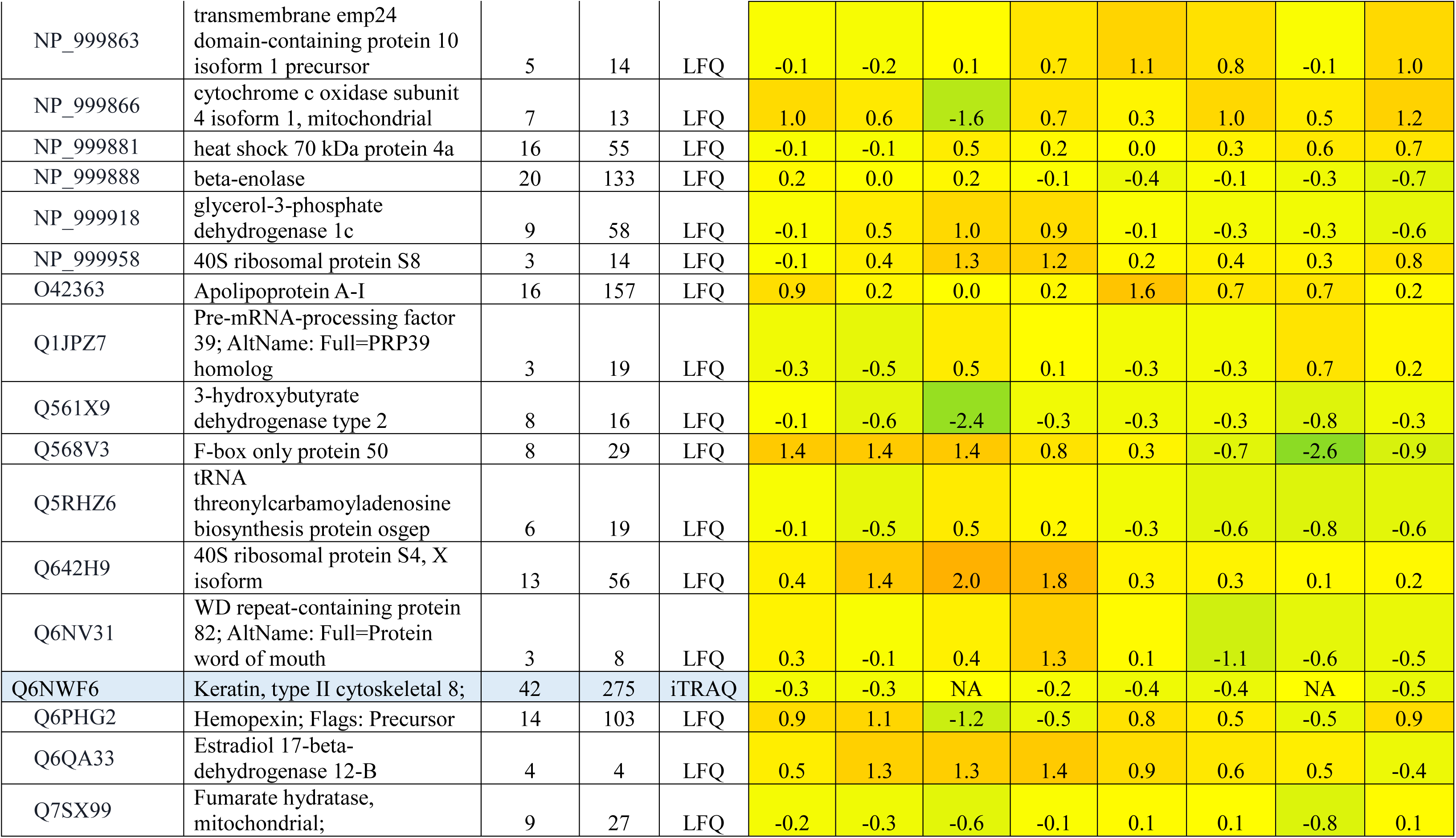
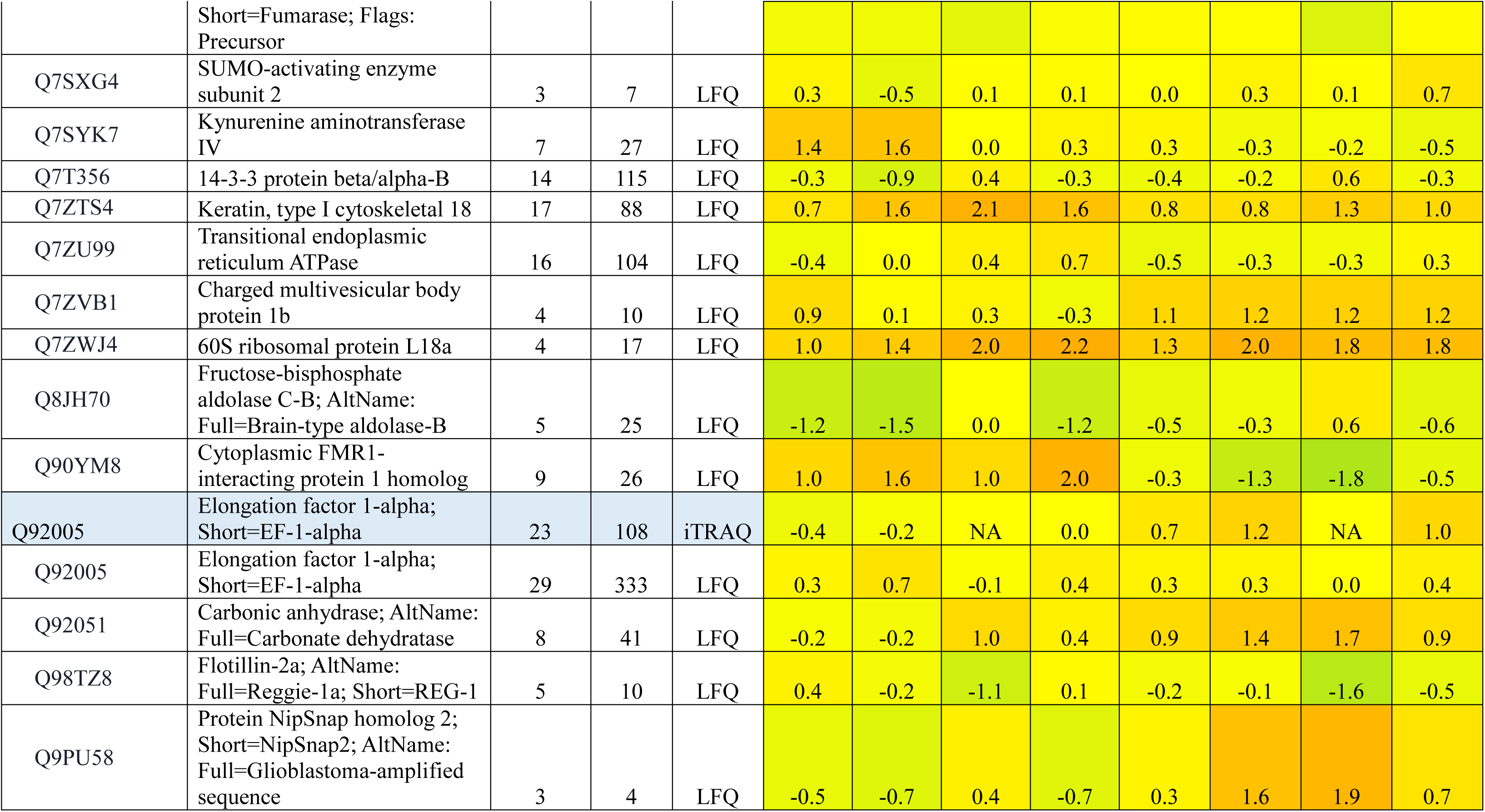
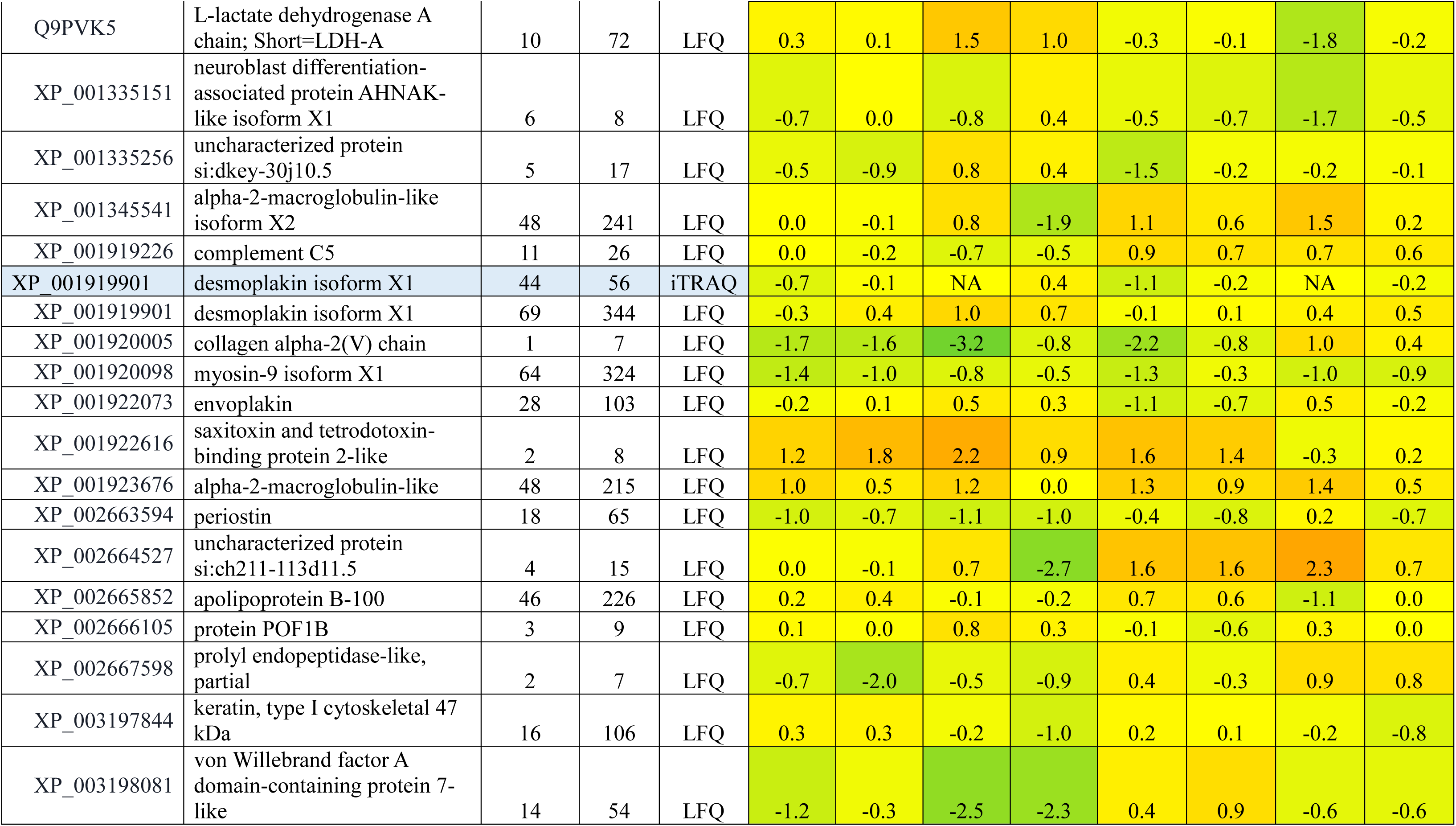
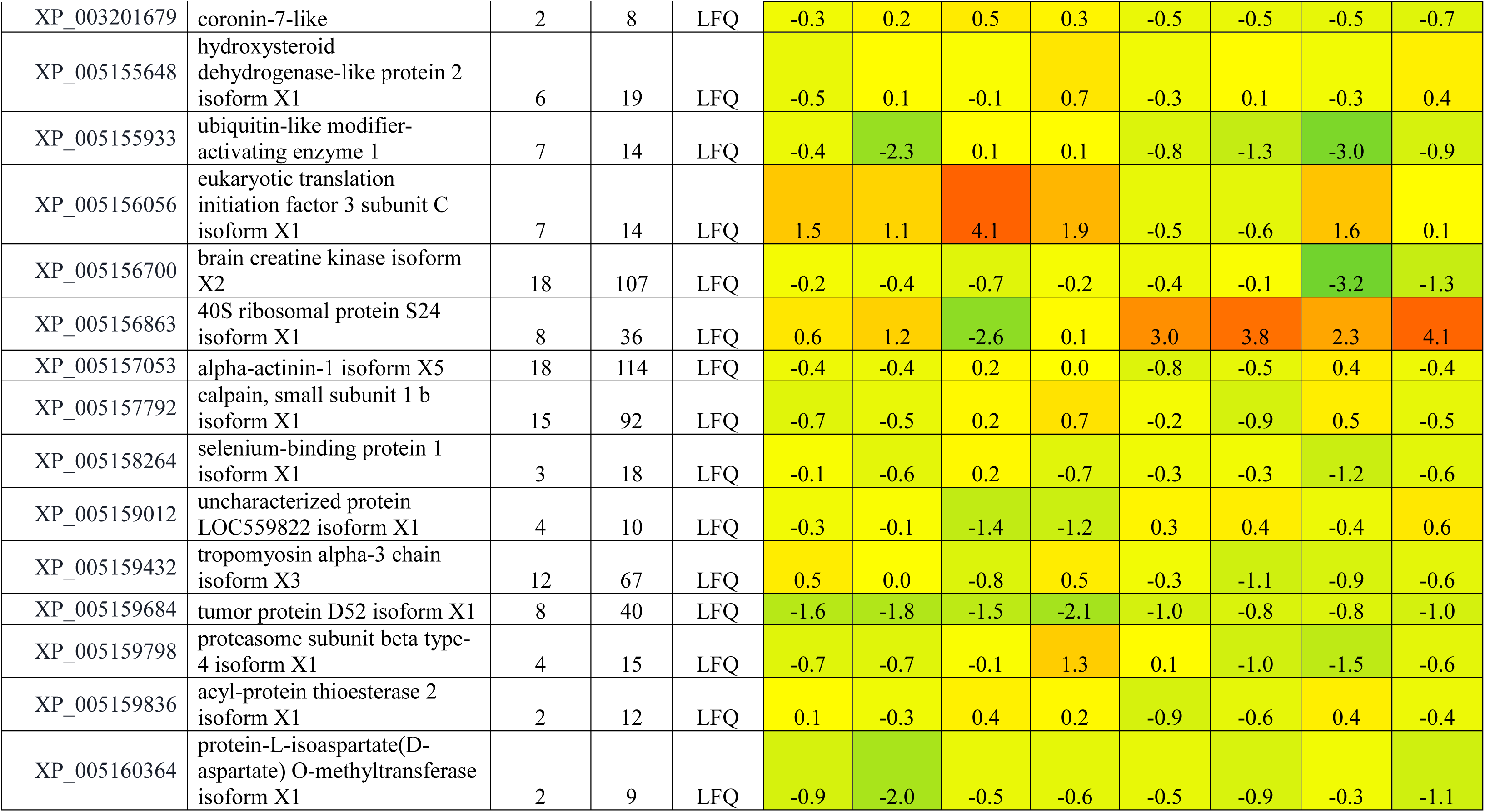
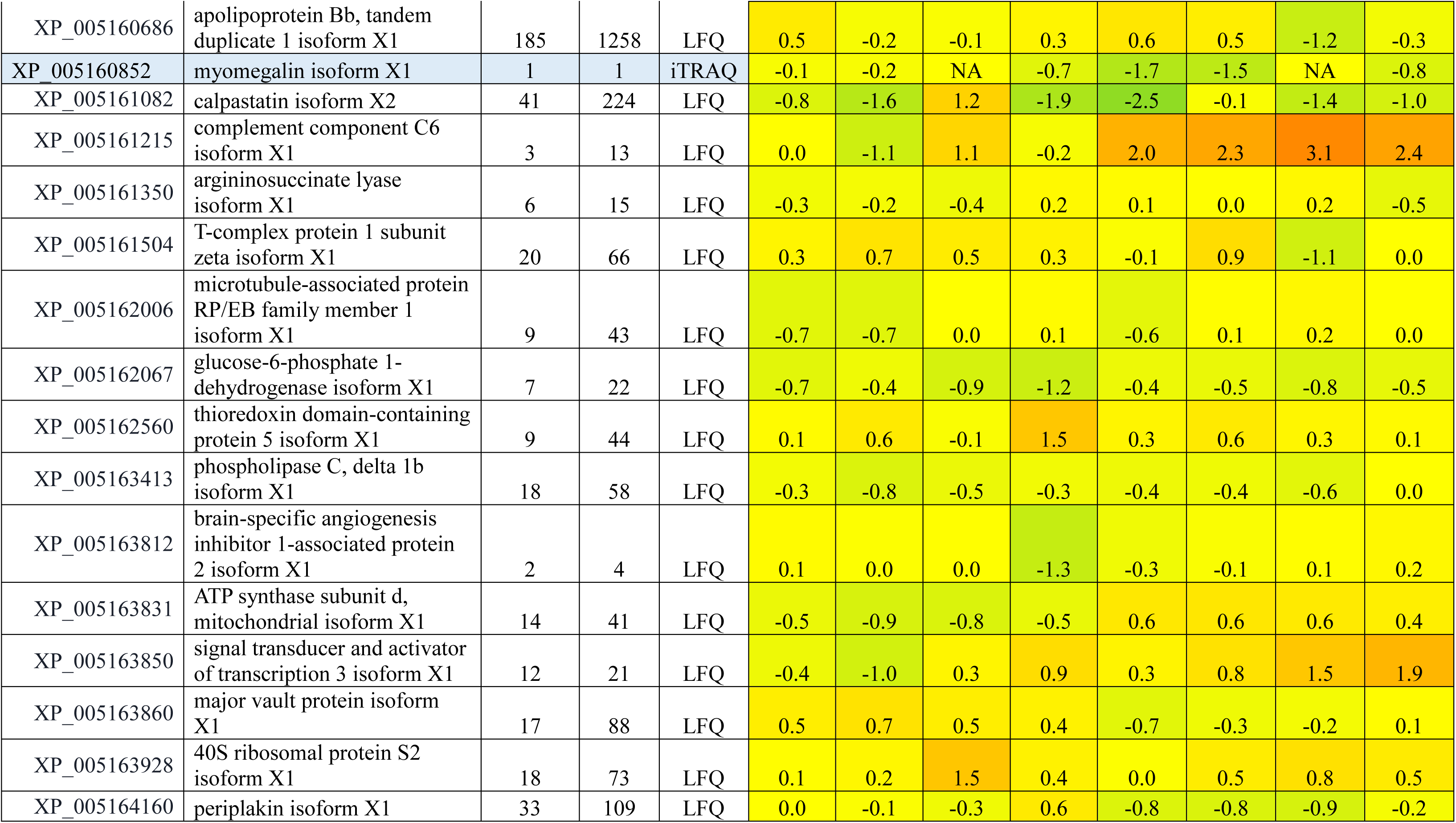
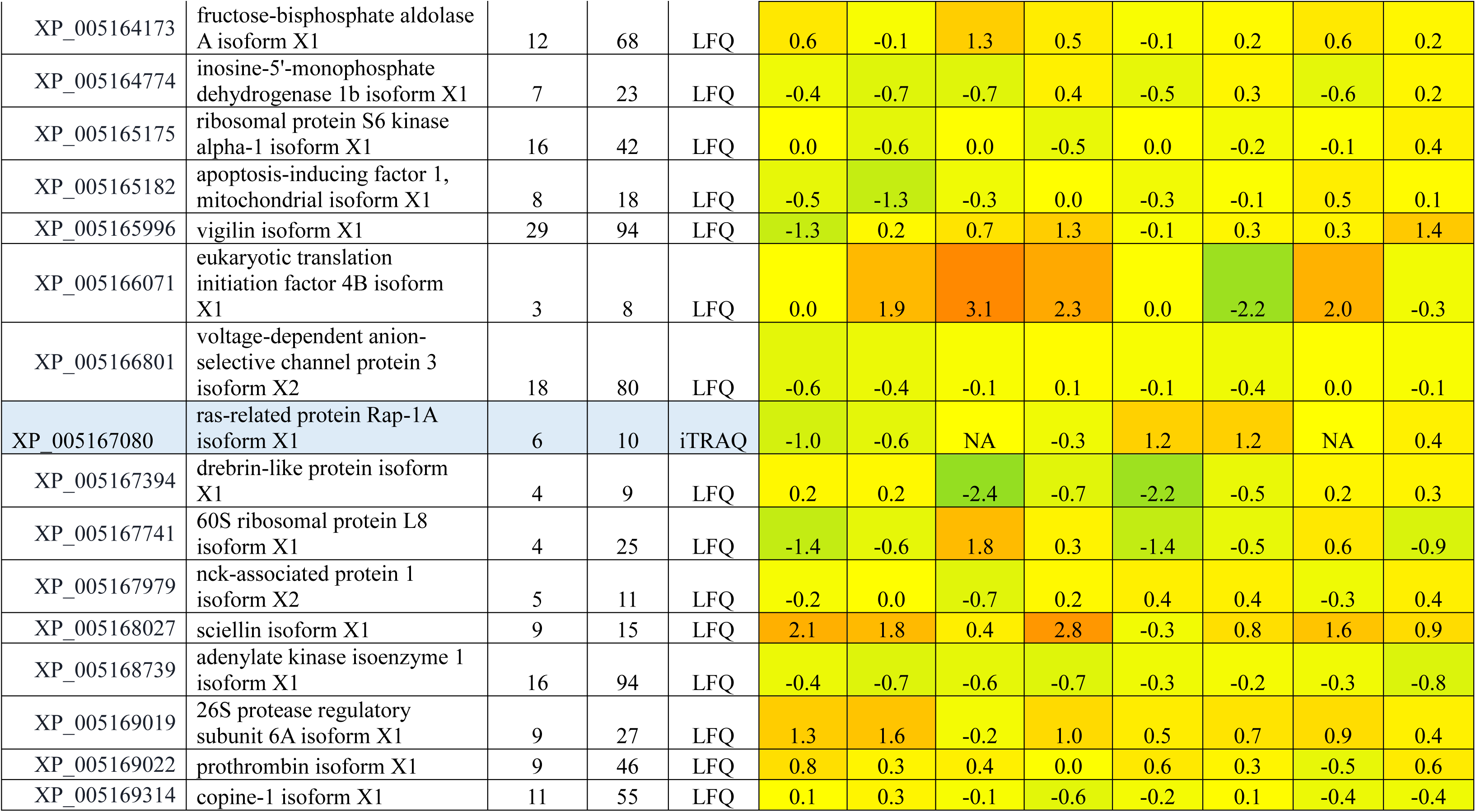
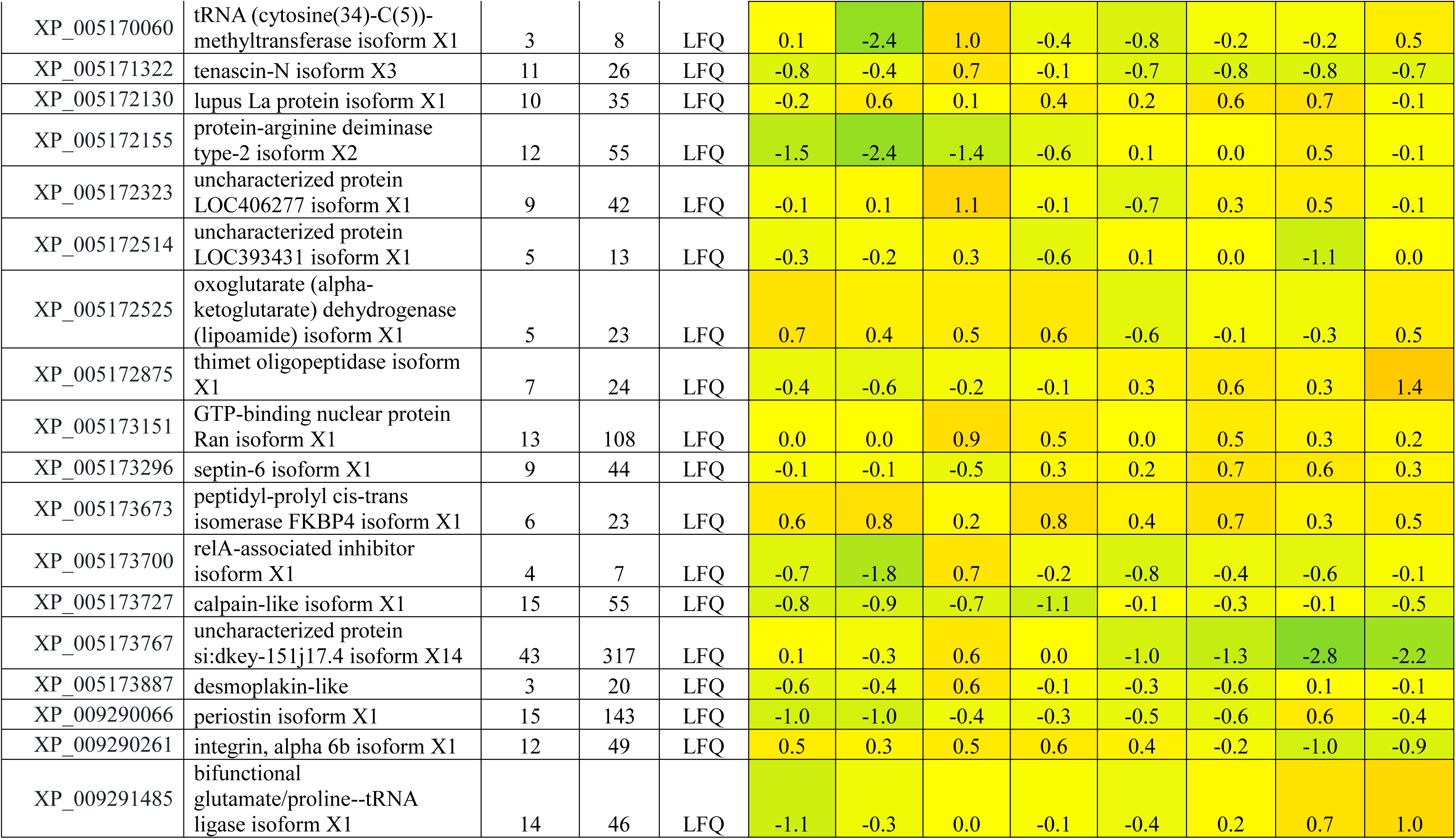
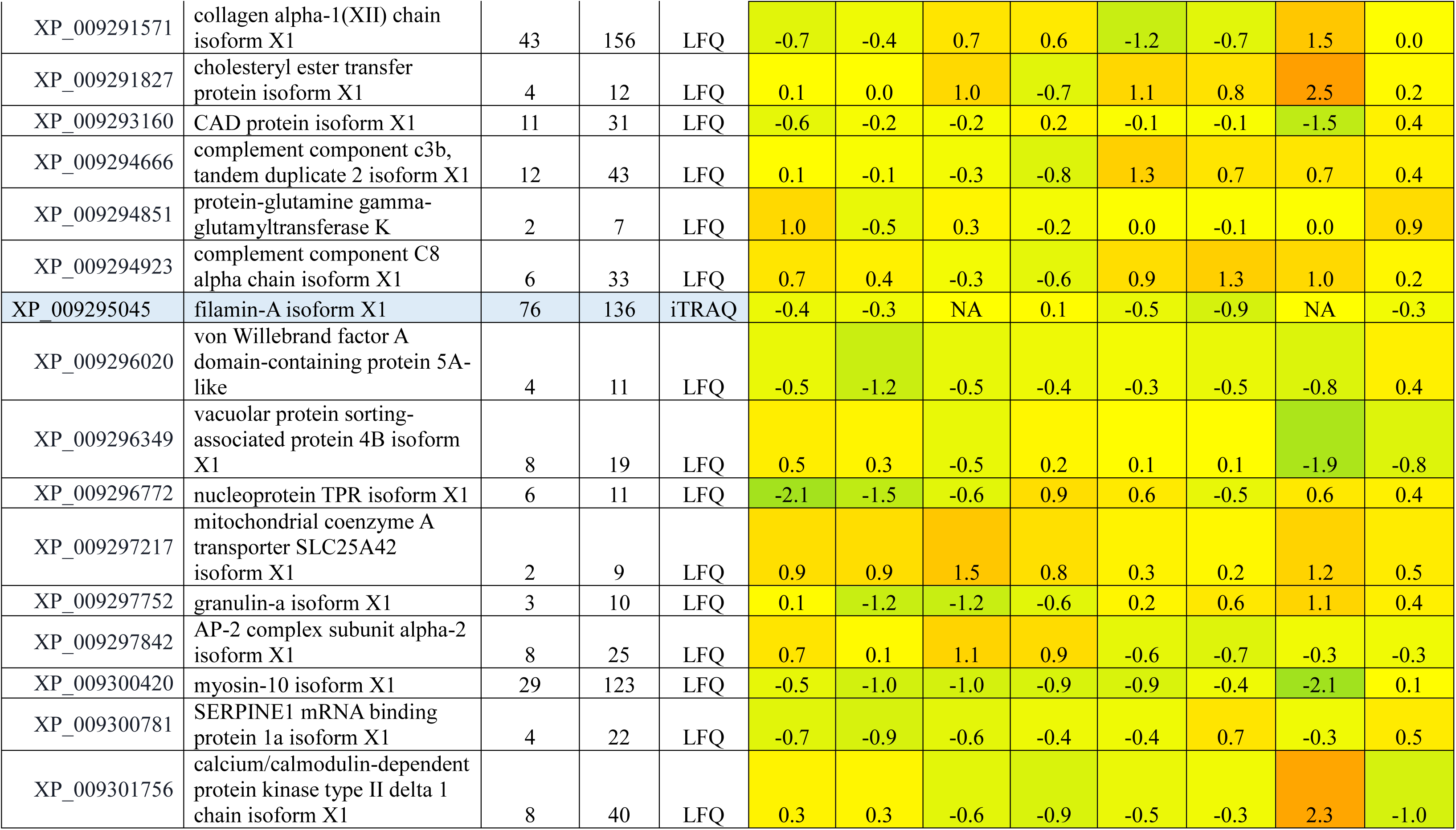
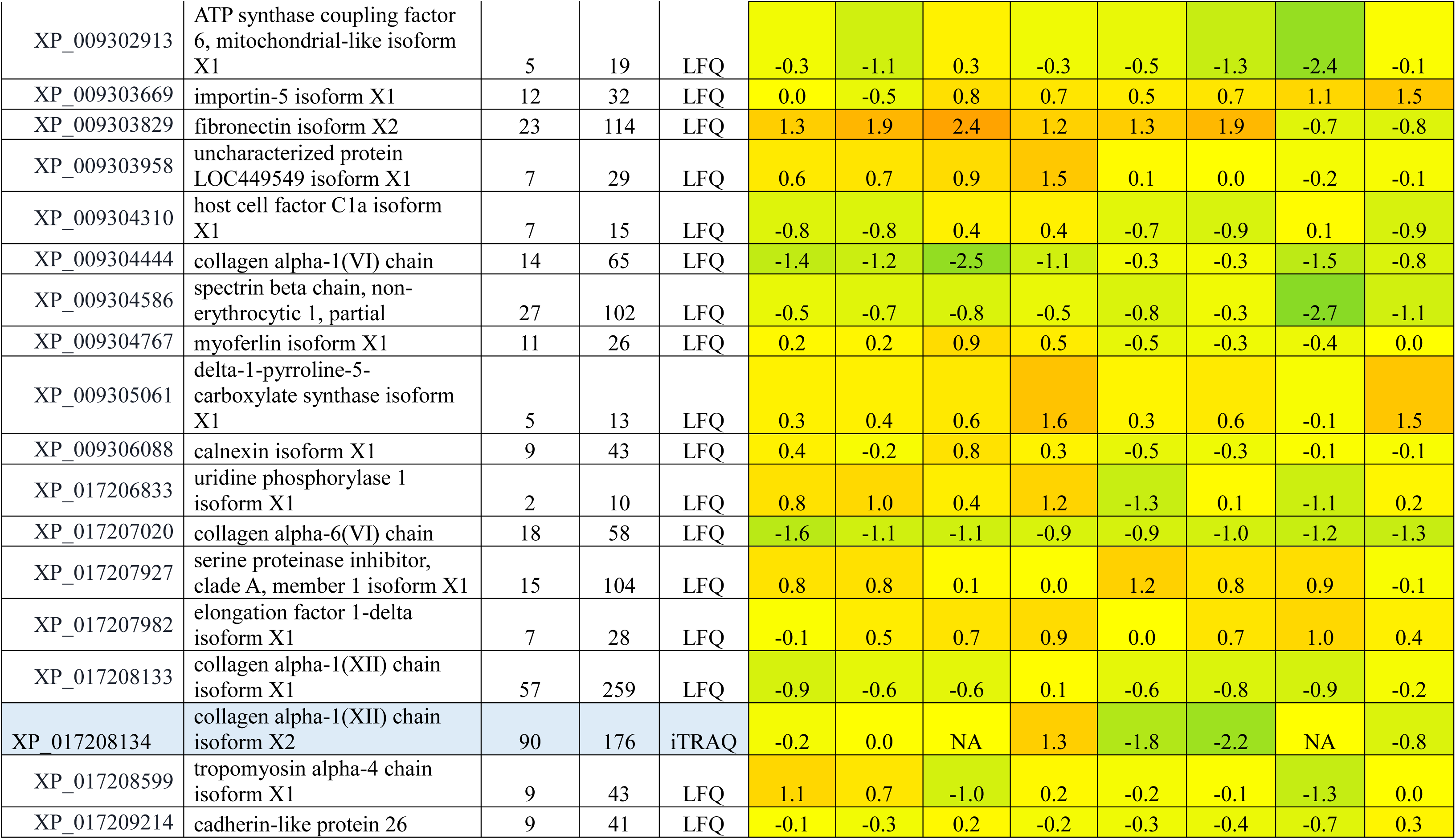
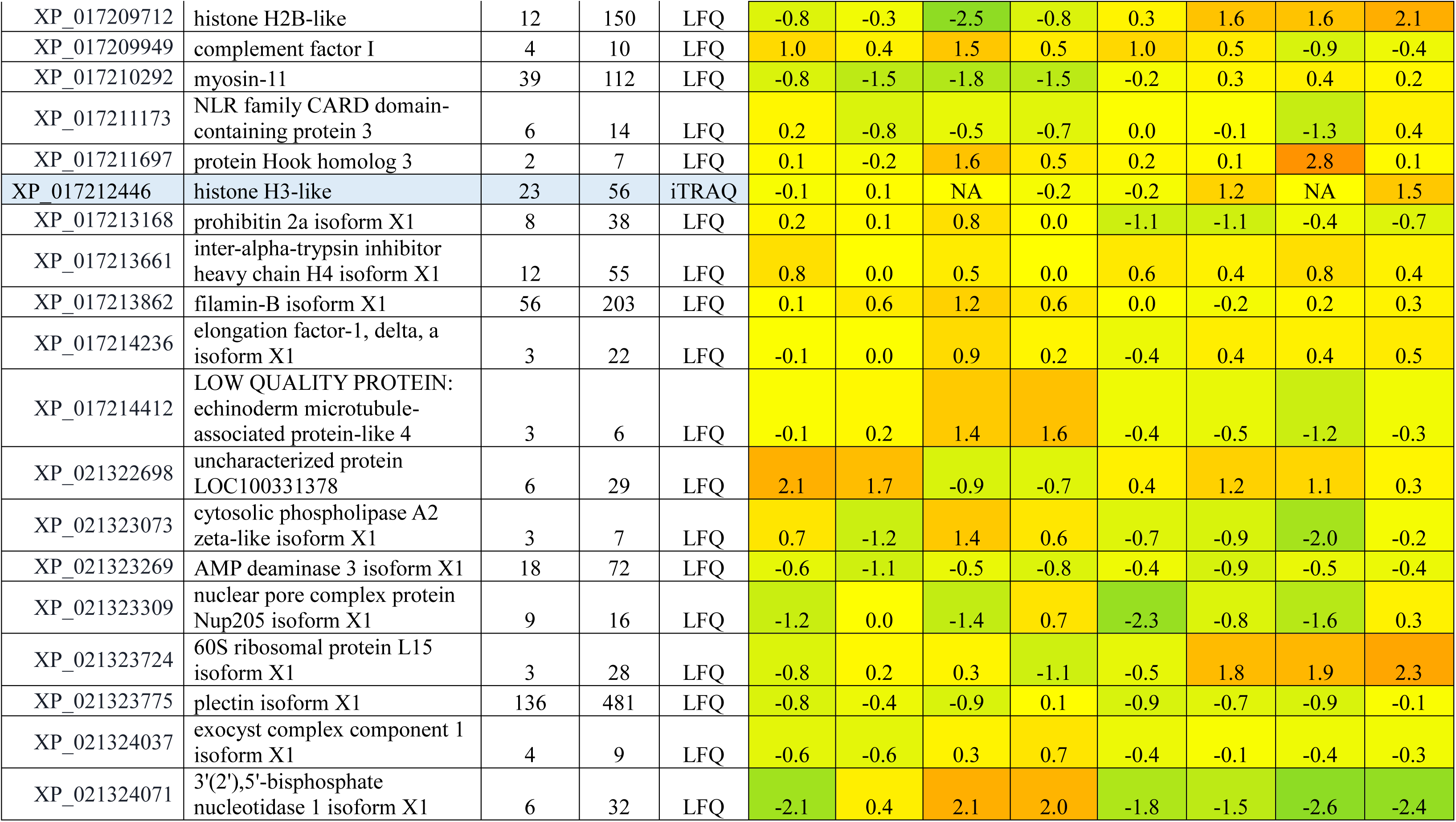
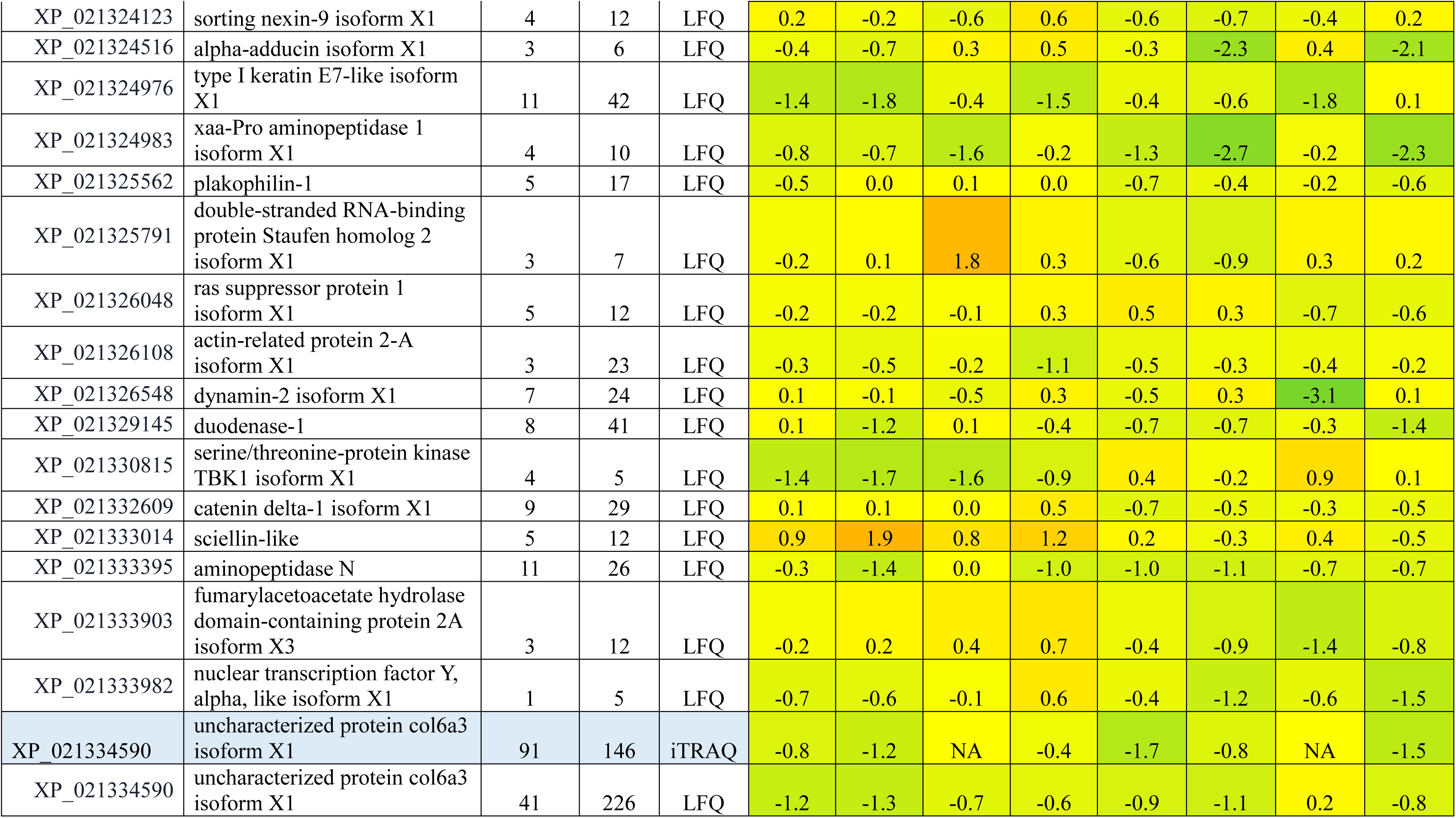
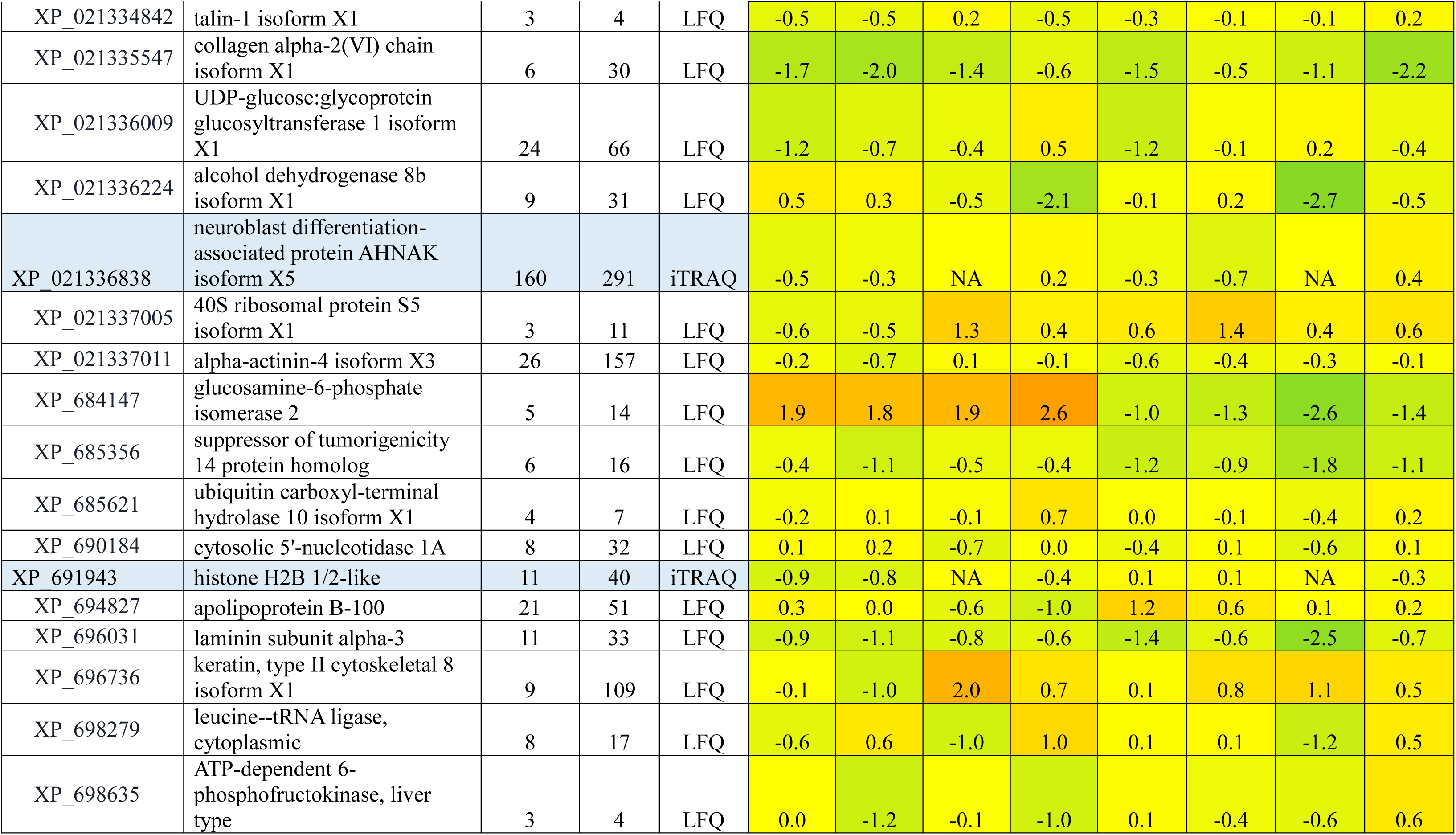
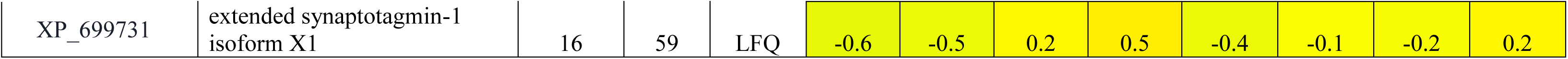
List of Proteins differentially expressed based on LFQ and iTRAQ analysis among young and old regenerating caudal fin tissue.

Differential protein expression analysis revealed distinct temporal proteomic signatures between young and aged zebrafish across all regeneration stages. During the early regenerative phase (1dpa), young zebrafish exhibited relatively stable or modest changes in proteins associated with cytoskeletal organization, extracellular matrix (ECM) components, and cellular architecture. Structural and cytoskeletal proteins including actin (actin, cytoplasmic 1), filamin-A, desmoplakin, and several keratin isoforms (keratin type II cytoskeletal 4, keratin type II cytoskeletal 5, and keratin type I cytoskeletal 15) were consistently detected, supporting early wound stabilization and epithelial remodeling. In contrast, aged zebrafish showed a pronounced reduction in the abundance of several structural and ECM-related proteins, including collagen alpha-1(XII) and collagen type VI alpha-3 (col6a3), along with cytoskeleton-associated proteins such as filamin-A and keratin family members. This reduction suggests an early deficit in extracellular matrix organization and structural support required for efficient wound healing and blastema initiation.

At the mid-regenerative stage (2 dpa), when blastema proliferation and tissue remodeling are prominent, young zebrafish displayed sustained or increased expression of proteins involved in cytoskeletal dynamics, cell adhesion, and intracellular signaling. Proteins such as epiplakin, desmoplakin, scinderin-like b, and adenylyl cyclase-associated protein 1 were prominently represented, indicating active cytoskeletal reorganization and membrane-associated signaling processes during blastema expansion. Additionally, metabolic and biosynthetic proteins including transketolase-like protein 2, pyruvate carboxylase a, and translation-associated factors such as elongation factor 1-alpha and elongation factor 2b were detected, reflecting increased metabolic and translational activity during regenerative tissue growth. Conversely, aged zebrafish continued to show reduced expression of key ECM and structural proteins, including collagen isoforms and keratin family members, along with altered abundance of proteins associated with cellular scaffolding and vesicle transport such as major vault protein and AHNAK. These differences suggest impaired structural remodeling and reduced cellular coordination during blastema proliferation in aged fins.

By the late regenerative stage (7 dpa), young zebrafish exhibited increased expression of proteins associated with tissue maturation, differentiation, and structural reinforcement.

Several cytoskeletal and epithelial structural proteins including keratin 8, keratin 97, and filamin-A were enriched, consistent with progressive fin tissue organization and restoration of epithelial integrity. Proteins involved in cellular metabolism and stress responses, such as heat shock protein HSP90-beta and endoplasmic reticulum chaperone BiP, were also detected, indicating active protein folding and stress adaptation during late-stage tissue maturation. Additionally, mitochondrial and metabolic proteins such as ATP synthase subunit alpha suggested increased energetic demands associated with regenerative outgrowth.

In aged zebrafish, however, the abundance of several ECM, cytoskeletal, and regulatory proteins remained comparatively lower or showed incomplete recovery during late regeneration. Proteins involved in cellular scaffolding, epithelial integrity, and metabolic support including keratin family members, colslagen isoforms, and metabolic enzymes displayed divergent expression patterns between young and aged fish, reflecting delayed or incomplete tissue remodeling during late-stage regeneration.

Notably, the identification of mitochondrial metabolic proteins, including ATP synthase subunit alpha, together with altered expression of metabolic enzymes such as pyruvate carboxylase a and transketolase-like protein 2, suggested potential involvement of mitochondrial energy metabolism in the regenerative process. These observations, combined with our transcriptomic findings showing dynamic regulation of mitochondrial electron transport chain genes during regeneration in young zebrafish, prompted further investigation into the role of mitochondrial function in regenerative efficiency.

### Rotenone-mediated mitochondrial inhibition alters mitochondrial protein expression

To further validate the involvement of mitochondrial activity during fin regeneration, RTPCR for mitochondrial complex genes was performed. Compared with young control zebrafish, rotenone-treated fish exhibited a marked reduction of mitochondrial genes levels during the regenerative process (Figure 4). Given that rotenone is a well-established inhibitor of mitochondrial complex I, down regulation of mitochondrial genes indicates disruption of electron transport chain function and compromised mitochondrial activity. These findings provide evidence supporting the effective inhibition of mitochondrial respiration following rotenone exposure and further reinforce the role of mitochondrial function in sustaining regenerative capacity.

### Ultrastructural analysis reveals mitochondrial abnormalities during impaired regeneration

Transmission electron microscopy (TEM) was performed to examine mitochondrial ultrastructure in regenerating fin tissues from young, aged, and rotenone-treated zebrafish (Figure 6). Mitochondria in regenerating tissues of young zebrafish displayed intact outer membranes and well-organized cristae architecture, consistent with structurally healthy mitochondria supporting active metabolic activity during regeneration. In contrast, mitochondria from aged zebrafish exhibited clear ultrastructural abnormalities, including elongation, irregular morphology, and partial disruption of cristae organization, suggesting compromised mitochondrial integrity during aging.

**Figure 6.**
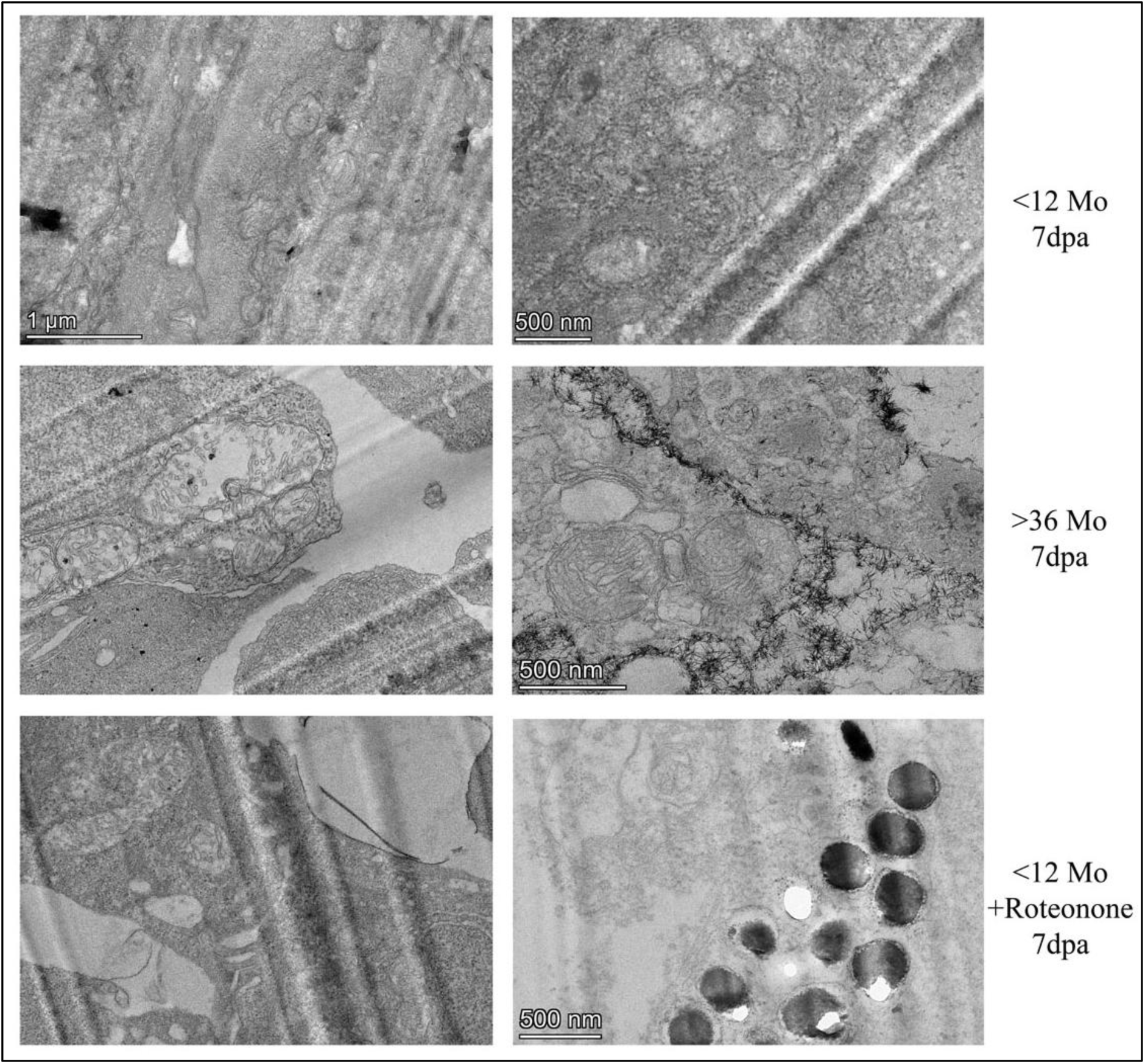
Ultrastructural analysis of regenerating tissue. Transmission electron microscopy images of zebrafish caudal fin tissue at 7 dpa. (a) Young. (b) Old. (c) Young treated with rotenone.

Rotenone-treated fish displayed more pronounced mitochondrial alterations. In addition to mitochondrial enlargement, many mitochondria appeared elongated and exhibited severely disrupted or fragmented cristae structures. Notably, electron-dense dark regions were frequently observed within the mitochondrial matrix, indicative of mitochondrial stress and structural damage. These pronounced ultrastructural defects indicate severe mitochondrial dysfunction following complex I inhibition. Collectively, these observations demonstrate that both aging and pharmacological inhibition of mitochondrial activity compromise mitochondrial structural integrity during regeneration, further supporting the critical role of mitochondrial function in sustaining regenerative capacity.

Together, these findings establish mitochondrial integrity and respiratory activity as critical determinants of efficient tissue regeneration, and suggest that mitochondrial dysfunction represents an important mechanistic driver of age-associated regenerative decline.

## Discussion

Aging is a critical biological factor that influences tissue homeostasis, repair capacity, and functional recovery. In this study, we demonstrate that aging significantly compromises caudal fin regeneration in zebrafish, affecting not only the rate and extent of structural regrowth but also behavioral recovery and the underlying molecular programs governing regeneration. By integrating morphological, behavioral, transcriptomic, proteomic, and mitochondrial functional analyses, our findings provide a comprehensive view of how aging alters regenerative competence in a well-established vertebrate regeneration model.

### Age-dependent decline in fin regenerative capacity

The attenuated fin regrowth observed in aged zebrafish is consistent with previous reports demonstrating a progressive decline in regenerative efficiency with advancing age across multiple tissues in zebrafish and other vertebrates. Reduced blastema formation, delayed wound closure, and incomplete tissue patterning in older fish suggest that aging interferes with early regenerative events that are critical for successful outgrowth [27]. The blastema is a transient, highly proliferative structure composed of lineage-restricted progenitors, and its formation requires coordinated activation of wound epidermis signaling, positional reprogramming, and cell cycle re-entry. Aging-associated impairments in any of these processes can lead to reduced regenerative output, as observed in our morphological analyses. Our findings further demonstrate that pharmacological inhibition of mitochondrial activity using rotenone phenocopies the regenerative defects observed in aged fish, resulting in delayed wound closure and reduced regenerative outgrowth. This observation suggests that mitochondrial dysfunction may represent a key mechanistic contributor to age-associated regenerative decline.

### Behavioral analysis reflects functional consequences of impaired regeneration

Beyond structural regeneration, our behavioral data reveal that aging influences functional recovery following fin amputation. Using the Novel Tank Test, we observed delayed locomotor recovery and reduced exploratory behavior in aged zebrafish during intermediate regenerative stages. Previous studies have established that fin integrity is essential for efficient swimming performance, maneuverability, and energy expenditure in zebrafish [28]. Thus, impaired fin regeneration in aged fish likely contributes directly to altered swimming dynamics.

Importantly, the Novel Tank Test is also a validated paradigm for assessing anxiety-like behavior in zebrafish. The reduced exploratory activity observed in aged fish during early and intermediate regeneration phases may therefore reflect heightened stress or anxiety responses following injury. Zebrafish behavioral phenotypes show translational relevance to mammalian anxiety and stress responses [26]. Aging is known to exacerbate stress sensitivity and reduce behavioral adaptability, potentially through dysregulation of neuroendocrine and inflammatory pathways [29]. Our findings suggest that regenerative impairment in aged zebrafish is accompanied by altered behavioral states, emphasizing that regeneration is not merely a morphological process but is closely linked to organismal physiology and behavior.

### Dysregulation of developmental and regenerative gene networks with aging

At the molecular level, our gene expression analysis revealed pronounced age-associated alterations in key developmental and regenerative gene families. In young zebrafish, robust induction of transcription factors such as *hoxa13b*, *msxb*, and *sox9a* was observed, consistent with their established roles in positional identity, progenitor maintenance, and lineage specification during fin regeneration [9, 10, 11]. The attenuated or delayed expression of these genes in aged fish suggests impaired reactivation of developmental programs necessary for blastema organization and patterning.

HOX and MSX genes are central regulators of positional memory and regenerative outgrowth. Their reduced expression in aged fins indicates compromised re-patterning capacity, which may underlie the incomplete structural restoration observed morphologically. Similar age-related declines in homeobox gene reactivation have been reported in other regenerative contexts, supporting the notion that aging limits the plasticity of adult tissues by restricting access to embryonic gene regulatory networks.

### Altered extracellular matrix remodeling and signaling homeostasis

Efficient regeneration requires extensive extracellular matrix (ECM) remodeling to support cell migration, proliferation, and differentiation. The reduced induction of keratin, collagen, and laminin genes in aged zebrafish suggests impaired epidermal integrity and ECM dynamics. Previous studies have demonstrated that keratins and extracellular matrix components play essential roles in wound epidermis formation and blastema signaling during fin regeneration, with keratin expression marking epidermal reorganization and provisional matrices rich in hyaluronic acid, fibronectin, and tenascin C contributing to cell migration and blastema dynamics in zebrafish [6].

Consistent with the idea that extracellular matrix dynamics underpin successful appendage regeneration, studies have shown that alterations in ECM-associated proteins such as FNDC3A lead to fin development and regeneration defects in zebrafish, implicating matrix remodeling as an essential component of proper tissue regrowth and mechanical support during regeneration [30]. Age-related ECM stiffening and altered matrix composition may therefore create a non-permissive environment for regeneration.

Similarly, differential expression of signaling and transporter genes such as *slc25a1a*, *slc25a1b*, and *tmem9* points toward disrupted metabolic and membrane homeostasis in aged regenerating tissue. SLC transporters regulate metabolite flux and mitochondrial function, processes that are increasingly recognized as key determinants of cellular metabolic state and regenerative potential [31]. Downregulation of *tmem9*, previously implicated in membrane-associated signaling, may further impair regenerative signal transduction [5].

### Aging-associated genes link regeneration, stress response, and genomic stability

The diminished induction of aging-associated genes such as *igf1*, *tert*, *hsp70*, and *atm* in aged zebrafish highlights a convergence between aging, stress response, and regenerative decline. IGF signaling promotes cell proliferation and survival during fin regeneration, while telomerase activity and DNA damage repair pathways are essential for maintaining genomic integrity in rapidly dividing blastemal cells [27]. Reduced expression of these genes may limit the proliferative capacity of progenitor cells and increase susceptibility to cellular stress, thereby constraining regenerative outcomes.

### Functional interaction networks reveal aging-sensitive regenerative modules

The STRING-based interaction and enrichment analysis provides a systems-level view of the molecular architecture underlying zebrafish caudal fin regeneration and highlights gene families whose coordinated activity is essential for successful tissue restoration. The enrichment of biological processes related to developmental patterning, transcriptional regulation, extracellular matrix organization, signal transduction, and stress response reflects the reactivation of embryonic-like programs during regeneration, a phenomenon well documented in zebrafish and other regenerative models [1],[2].

Importantly, many of the functional modules identified in our analysis, including Notch signaling, kinase-mediated pathways, ECM remodeling, and transmembrane transport, are known to be particularly vulnerable to aging-related dysregulation. Aging often disrupts the timing and coordination of transcriptional networks rather than eliminating them entirely, resulting in delayed or incomplete regenerative responses [8]. The enrichment of extracellular matrix components such as collagens and laminins highlights the importance of a dynamically remodeled regenerative niche, which is increasingly recognized as compromised with age due to altered ECM composition, reduced cell-matrix interactions, and impaired mechanotransduction [13,32]. The association of solute carrier (SLC) transporters and TMEM proteins with metabolic and membrane-related functions further suggests that efficient regeneration requires tight control of cellular homeostasis and energy balance. Aging is known to disrupt mitochondrial function, nutrient transport, and ionic regulation, which can indirectly constrain blastema cell proliferation and differentiation [33]. The integration of aging-associated genes such as *igf1*, *tert*, *hsp70*, and *atm* within this network further highlights the close coupling between regenerative potential and cellular maintenance mechanisms, including growth factor signaling, proteostasis, telomere stability, and DNA damage response.

Together, the functional interaction landscape depicted in figure 5 supports the concept that regeneration is governed by a highly interconnected molecular network, in which transcriptional regulators, signaling pathways, structural components, and stress-response systems operate in concert.

### Proteomic alterations reveal age-dependent disruption of structural and metabolic regeneration programs

While transcriptomic analyses provide valuable insights into regulatory mechanisms, successful tissue regeneration ultimately depends on the coordinated synthesis and function of proteins that execute cellular processes. Our integrative proteomic analysis, combining both label-free quantification (LFQ) and iTRAQ approaches, provides direct evidence that aging disrupts the protein-level architecture underlying zebrafish caudal fin regeneration.

Proteomic profiling across early (1 dpa), mid (2 dpa), and late (7 dpa) regenerative stages revealed age-associated alterations in proteins involved in extracellular matrix organization, cytoskeletal dynamics, metabolic regulation, stress responses, and translational control. Importantly, the concordance between LFQ and iTRAQ datasets, with all 32 proteins identified in the iTRAQ analysis also detected in the LFQ dataset, highlights the robustness of the identified regeneration-associated protein signatures.

Structural and cytoskeletal proteins such as actin, filamin-A, keratin isoforms, and desmoplakin were prominently represented in young regenerating fins, supporting early wound stabilization and epithelial reorganization. These proteins play key roles in maintaining cellular architecture, facilitating cell migration, and supporting blastema formation during appendage regeneration [34]. In contrast, aged fins displayed reduced abundance of these structural proteins, indicating compromised cytoskeletal organization and impaired tissue remodeling.

Similarly, extracellular matrix components such as collagen alpha-1(XII) and collagen VI were markedly reduced in aged regenerating tissue. FACIT collagens such as collagen XII regulate collagen fibril organization and mechanical stability of connective tissues. Reduced abundance of these ECM components may therefore compromise blastemal cell anchorage and matrix remodeling, ultimately limiting regenerative outgrowth [32].

Metabolic enzymes including pyruvate carboxylase and transketolase-like proteins were also differentially represented between young and aged fins. These enzymes participate in central metabolic pathways that regulate cellular energy production and biosynthetic processes [31]. Their altered expression suggests that aging disrupts metabolic reprogramming necessary to sustain rapid cellular proliferation during regeneration.

### Mitochondrial dysfunction emerges as a key determinant of regenerative decline

An important insight from our integrative analysis is the emerging role of mitochondrial function in regulating regenerative efficiency. Both transcriptomic and proteomic datasets revealed dynamic regulation of mitochondrial components during regeneration in young zebrafish, including genes encoding electron transport chain subunits and mitochondrial metabolic proteins such as ATP synthase.

Mitochondria are central regulators of cellular energy production, redox balance, and metabolic signaling, all of which are essential for supporting the rapid proliferation and differentiation of blastemal cells [14]. Aging is widely associated with mitochondrial dysfunction, characterized by reduced respiratory efficiency, increased oxidative stress, and impaired metabolic flexibility [33]. Our findings suggest that similar mitochondrial deficits may contribute to impaired regenerative responses in aged zebrafish.

### Functional validation of mitochondrial involvement in regeneration

To directly investigate the functional role of mitochondrial activity during regeneration, we pharmacologically inhibited mitochondrial complex I using rotenone. Rotenone treatment resulted in pronounced impairment of fin regeneration, closely resembling the regenerative defects observed in aged zebrafish. This phenocopy strongly supports the hypothesis that mitochondrial activity is a critical determinant of regenerative capacity. Complementary ultrastructural analysis using transmission electron microscopy revealed severe mitochondrial abnormalities in aged and rotenone-treated fish, including elongated mitochondria, disrupted cristae architecture, and electron-dense matrix deposits indicative of mitochondrial stress and dysfunction.

Collectively, these molecular and ultrastructural findings demonstrate that mitochondrial integrity and respiratory activity are essential for supporting the metabolic demands of regenerating tissues.

### Integrative perspective and significance

Taken together, our findings reveal that aging impairs zebrafish caudal fin regeneration through coordinated disruptions in developmental gene networks, extracellular matrix remodeling, metabolic regulation, and mitochondrial function. The convergence of morphological, behavioral, gene regulatory networks, proteomic and ultrastructural evidence points out mitochondrial dysfunction as a mechanistic driver linking aging to regenerative decline.

By building upon our previous transcriptomic analysis of young adult zebrafish, this study provides direct evidence that aging reshapes regeneration-associated gene expression programs, offering valuable insights into why regenerative capacity declines with age and demonstrates that the mitochondrial metabolic integrity is not merely a consequence of regeneration but an essential determinant of regenerative efficiency. Understanding how mitochondrial function influences tissue repair may therefore provide new avenues for enhancing regenerative capacity in aging organisms and may inform therapeutic strategies aimed at promoting tissue repair in age-related degenerative conditions.

### Conclusion

This study demonstrates that aging significantly impairs zebrafish caudal fin regeneration through coordinated disruptions in structural, molecular, and metabolic processes. In young zebrafish, caudal fin regeneration is supported by the coordinated activation of developmental transcription factors, dynamic extracellular matrix remodeling, and metabolic reprogramming, which collectively enable blastema formation and tissue outgrowth. These processes are sustained by intact mitochondrial function, providing the energetic and biosynthetic capacity required for rapid cell proliferation and differentiation. In aged zebrafish, however, dysregulation of regenerative gene networks, reduced abundance of structural and metabolic proteins, and impaired mitochondrial gene expression converge to create a metabolically constrained regenerative environment. Functional inhibition of mitochondrial activity using rotenone recapitulated many of the structural and molecular defects observed in aged animals, further supporting a causal role for mitochondrial dysfunction in limiting regenerative efficiency. Ultrastructural mitochondrial abnormalities observed in aged and rotenone-treated tissues further reinforce the idea that compromised mitochondrial integrity may underlie the metabolic insufficiency associated with aging-related regenerative decline.

By integrating morphological, behavioral, transcriptomic, proteomic, and ultrastructural analyses, this study provides a comprehensive framework for aging-associated decline of regenerative capacity in zebrafish. These findings extend current understanding of vertebrate regeneration by identifying mitochondrial metabolic integrity as a critical regulator of tissue repair. Future studies aimed at preserving mitochondrial function or enhancing cellular metabolic resilience may offer promising strategies for improving regenerative outcomes in aging organisms.

## Acknowledgement

The authors are thankful to Dr. B Raman and Ms. Y Kameshwari for their help at the CCMB Proteomic facility and Dr. Khushboo for the laboratory help.

## Abbreviations

hpa: hours post amputation
dpa: days post amputation;

